# Differential Brain Atrophy Patterns and Neurogenetic Profiles in Cognitively-Defined Alzheimer’s Disease Subgroups

**DOI:** 10.1101/2020.08.04.235762

**Authors:** Colin Groot, Michel J. Grothe, Shubhabrata Mukherjee, Irina Jelistratova, Iris Jansen, Anna Catharina van Loenhoud, Shannon L. Risacher, Andrew J. Saykin, Christine L. Mac Donald, Jesse Mez, Emily H. Trittschuh, Gregor Gryglewski, Rupert Lanzenberger, Yolande A.L. Pijnenburg, Frederik Barkhof, Philip Scheltens, Wiesje M. van der Flier, Paul K. Crane, Rik Ossenkoppele

**Affiliations:** Department of Neurology & Alzheimer Center, Amsterdam University Medical Center – Location VU University medical center, Amsterdam, the Netherlands; Unidad de Trastornos del Movimiento, Servicio de Neurología y Neurofisiología Clínica, Instituto de Biomedicina de Sevilla, Hospital Universitario Virgen del Rocío/CSIC/Universidad de Sevilla, Seville, Spain; German Center for Neurodegenerative Diseases (DZNE), Rostock, Germany; Department of Medicine, University of Washington, Seattle, WA, USA; Department of Complex Trait Genetics, Center for Neurogenomics and Cognitive Research, Amsterdam Neuroscience, VU University, Amsterdam, the Netherlands; Indiana University School of Medicine, Indianapolis, IN, USA; Neurological Surgery, University of Washington, Seattle, WA, USA; Department of Neurology, Boston University School of Medicine, Boston, MA, USA; Alzheimer’s Disease Center, Boston University School of Medicine, MA, USA; Psychiatry & Behavioral Science, University of Washington, Seattle, WA, USA; Veterans Affairs Puget Sound Health Care System, Geriatric Research, Education, & Clinical Center, Seattle, WA, USA; Department of Psychiatry and Psychotherapy, Medical University of Vienna, Vienna, Austria; Department of Radiology and Nuclear Medicine, Amsterdam University Medical Center – Location VU University medical center, Amsterdam, the Netherlands; University College London, Institutes of Neurology & Healthcare Engineering, London, United Kingdom; Epidemiology and Biostatistics, Amsterdam University Medical Center – Location VU University medical center, Amsterdam, the Netherlands; Lund University, Clinical Memory Research Unit, Lund, Sweden

**Author notes:** Colin Groot, Amsterdam UMC - VU University medical center, Boelelaan 1117, 1081 HV Amsterdam, the Netherlands. These authors contributed equally to this work (Paul K. Crane, Rik Ossenkoppele).

## Abstract

Elucidating mechanisms underlying the clinical heterogeneity observed among individuals with Alzheimer’s disease (AD) is key to facilitate personalized treatments. We categorized 679 individuals with AD into subgroups based on a relative impairment in one cognitive domain (i.e. AD-Memory, AD-Executive-Functioning, AD-Language and AD-Visuospatial-Functioning). We compared atrophy patterns derived from MRI and identified patterns that closely matched the respective cognitive profiles, i.e. medial temporal lobe atrophy in AD-Memory, fronto-parietal in AD-Executive-Functioning, asymmetric left-temporal in AD-Language, and posterior in AD-Visuospatial-Functioning. We then determined spatial correlations between subgroup-specific atrophy and a transcriptomic atlas of gene expression, which revealed both shared (e.g. mitochondrial respiration and synaptic function/plasticity) and subgroup-specific (e.g. cell-cycle for AD-Memory, protein metabolism in AD-Language, and modification of gene expression in AD-Visuospatial-Functioning) biological pathways associated with each subgroup’s atrophy patterns. We conclude that cognitive heterogeneity in AD is related to neuroanatomical differences, and specific biological pathways may be involved in their emergence.

## Introduction

The clinical phenotype of Alzheimer’s disease (AD) is typically characterized by prominent memory impairment. However, there is considerable variation in the cognitive manifestation of AD that can also present with substantial deficits in cognitive domains other than memory (Lam et al., 2013a). In roughly 5% of late-onset (LOAD) and 22-64% of early-onset (EOAD) AD patients, cognitive impairment in domains other than memory constitute the most prominent feature (Mendez, 2017). Moreover, at the ends of the clinical spectrum reside various atypical variants, which are characterized by early and predominant impairments in a single cognitive domain. Examples of well-characterized atypical variants of AD include posterior cortical atrophy (PCA) (Crutch et al., 2017) characterized by visual processing deficits such as simultanagnosia and space perception problems, and logopenic variant primary progressive aphasia (lvPPA) (Gorno-Tempini et al., 2008) where impairments in language function (e.g. single-word retrieval and sentence repetition) are prominent.

While these atypical variants represent phenotypical extremes, there is substantial interindividual cognitive variability in persons who do not meet clinical criteria for these specific variants of AD and thus remain classified under the moniker of “typical AD” (Ossenkoppele et al., 2019). A framework has been proposed to categorize people with typical AD into cognitively-defined subgroups based on their relative performance on domain-specific cognitive tests (Crane et al., 2017a). Earlier studies that used this approach have shown that subgroup membership had important clinical implications, as subgroups characterized by non-amnestic impairments showed faster progression compared to memory-predominant subgroup (Mez et al., 2013a). A deeper understanding of the underlying mechanisms that govern the differences between cognitively-defined subtypes of typical AD and their clinical implications might identify differential pathways that play a role in the pathogenesis of AD, improve the accuracy of diagnosis and prognosis, and aid in the development of personalized medicine strategies and design of clinical trials.

Cognitive phenotypes that characterize atypical AD variants (e.g. lvPPA, PCA) are associated with marked clinical differences and regional variations in neurodegeneration (Lam et al., 2013b; Ossenkoppele et al., 2015b). Moreover, differences in regional gene expression profiles associate with regional differences in anatomical (Whitaker et al., 2016) and functional (Richiardi et al., 2015) properties of the brain, as well as with selective regional vulnerability to neurodegenerative disease (Freeze et al., 2019; Grothe et al., 2018; Sepulcre et al., 2018). Based on these observations, we aimed to address three research objectives with regard to cognitively-defined subgroups within the broad spectrum of ‘typical’ AD individuals. First, to replicate previous findings with regards to faster clinical progression for non-amnestic subgroups and assess whether this also translates into higher mortality rates. Second, to examine whether cognitively-defined subgroups associate with regional variations in atrophy. Third, to explore specific genetic pathways that might explain differences in regional susceptibility to neurodegeneration, by relating subgroup-specific atrophy to brainwide gene expression profiles based on a stereotactic characterization of the transcriptional architecture of the human brain as provided by the Allen human brain atlas (Hawrylycz et al., 2015).

## Results

### Subgroup characteristics

Of the 679 amyloid-β positive individuals who were eligible for categorization into AD-subgroups (see *“Participants”* section), about one third (n=231, 34%) were classified as having no domain with relative impairments compared to the average across the four cognitive domains within each subject, meaning these individuals had similar levels of impairment across all four cognitive domains. These individuals were classified into the “AD-No Domains” (AD-ND) group, and served as a reference group. Forty-one subjects (6%) were classified as AD-Memory (AD-Mem), 117 (17%) as AD-Executive-Functioning (AD-Exec), 33 (5%) as AD-Language (AD-Lang) and 171 (25%) as AD-Visuospatial-Functioning (AD-VS). These AD-subgroups are named according to the cognitive domain that showed substantial relative impairment compared to a global average across cognitive domains within each subject (see *“Subgroup classification”* in the Methods section). In addition, 86 subjects (13%) showed substantial relative impairment on more than one domain and were classified as AD-Multiple. Because of the heterogeneous composition of the AD-Multiple group (Supplemental-Fig.-1), our main analyses are focused on the other subgroups. Table-1 displays the demographic and clinical characteristics for the whole sample and cognitively-defined subgroups, and all pairwise differences between groups are given in the legend. Mean age of the total sample was 66.2±7.4, 47% were male, MMSE was 21.2±5.1 and 69% were *APOEε4* positive. Using the AD-ND group as the reference we observed that AD-VS were younger (63.7±7.2 *vs* 66.9±7.7, p<0.01) and AD-Lang had a lower *APOEε4* prevalence (51.5 *vs* 72.7%, p<0.01). AD-Mem showed the highest *APOEε4* prevalence (90.2%), which was significantly higher than in AD-ND (72.7%, p=0.017). Furthermore, compared to AD-ND, AD-Exec had more global atrophy (0.37±0.04 *vs* 0.38±0.04, p<0.01) and AD-Mem had less atrophy (0.41±0.04 *vs* 0.38±0.04, p<0.01).

**Table-1.** Demographic and clinical characteristics of the sample

**Cognitively defined Alzheimer's disease subgroup**
|  | Total | AD-Memory | AD-Executive F. | AD-Language | AD-Visuospatial F. | AD-Multiple | AD-No Domains | p for group effect |
| --- | --- | --- | --- | --- | --- | --- | --- | --- |
| **N** | 679 | 41 (6%) | 117 (17%) | 33 (5%) | 171 (25%) | 86 (13%) | 231 (34%) |  |
| **Age at AD diagnosis (SD)** | 66.3 (7.43) | 68.3 (6.6) | 68.0 (6.6) | 64.4 (6.4) | 63.7 (7.3) | 66.8 (7.4) | 66.9 (7.7) | <0.001<sup>a</sup> |
| **Early onset, <60 years** | 282/679 (42%) | 11/41 (27%) | 40/117 (34%) | 17/33 (52%) | 95/171 (56%) | 33 (38%) | 86/231 (37%) | <0.001<sup>b</sup> |
| **Sex, male** | 318 (47%) | 17 (42%) | 60 (51%) | 18 (55%) | 79 (46%) | 48 (56%) | 96 (42%) | 0.353 |
| **MMSE (SD)** | 21.2 (5.1) | 23.68 (2.8) | 20.40 (5.5) | 17.24 (5.7) | 22.26 (4.8) | 19.97 (5.1) | 21.46 (4.8) | <0.001<sup>c</sup> |
| **Edu, median [IQR]<sup>^</sup>** | 5 [4. 6] | 5 [4. 6] | 5 [4. 6] | 5 [5. 6] | 5 [4. 6] | 5 [4. 6] | 5 [4. 6] | 0.038<sup>d</sup> |
| **APOEε4 positive** | 467 (69%) | 37 (90%) | 75 (64%) | 17 (52%) | 113 (66%) | 57 (66%) | 168 (73%) | 0.002<sup>e</sup> |
| **Total gray matter (SD)<sup>^^</sup>** | 0.38 (0.04) | 0.41 (0.04) | 0.37 (0.04) | 0.38 (0.04) | 0.39 (0.04) | 0.37 (0.04) | 0.38 (0.04) | <0.001<sup>f</sup> |

Differences were assessed using ANOVA with post-hoc independent samples t-tests and $\chi^2$ - tests with post-hoc Fishers exact tests, where appropriate. The Kruskal-Wallis tests was used for the non-normally distributed education variable. The AD-Multiple group was excluded from the comparisons. *APOE* – Apolipoprotein E
a – AD-Mem>AD-Lang and AD-VS; AD-Exec>AD-Lang and AD-VS; AD-ND>AD-VS
b – AD-VS>AD-Mem, AD-Exec and AD-ND; AD-Lang>AD-Mem
c – AD-Mem, AD-Exec, AD-VS, AD-ND>AD-Lang; AD-Mem>AD-Exec, AD-ND; AD-VS>AD-Exec
d – AD-Mem, AD-Lang and AD-VS>AD-Exec
e – AD-Mem>AD-Exec, AD-Lang and AD-ND; AD-Exec, AD-VS and AD-ND>AD-Lang; AD-VS>AD-Exec
f – AD-Mem>AD-Exec, AD-Lang, AD-VS and AD-ND; AD-VS and AD-ND>AD-Exec
<sup>^</sup> - Assessed using the qualitative Dutch Verhage scale ranging from 1 for not completing primary school to 7 for a completed university degree
<sup>^^</sup> - Ratio of gray matter volume to total intracranial volume that indicates whole brain gray matter atrophy. Lower values indicate more atrophy

### Rates of clinical disease progression and mortality

Linear-mixed effects models revealed higher baseline MMSE scores in AD-Mem (β(SE)=1.84(0.85), p=0.03) and AD-VS (1.19(0.51), p=0.02), and lower scores in AD-Lang (−4.0(0.98), p<0.01) when compared to AD-ND. Furthermore, all subgroups progressed faster over time on the MMSE than AD-ND (−0.50(0.18), p=0.01 for AD-Exec; −1.61(0.57), p<0.01 for AD-Lang; −0.56(0.13), p<0.01 for AD-VS), except for AD-Mem (−0.29(0.23), p=0.20, Fig.-1A). Cox regression analyses assessing age-adjusted mortality hazard ratios between subgroups using the AD-ND group as reference, revealed a higher mortality rate in AD-Exec (HR[95%CI]=1.94[1.38-2.73], p<0.01), AD-Lang (2.16[1.29-3.63], p=0.05) and AD-VS (1.48[1.07-2.00], p=0.02). There were no differences in mortality rates between AD-Mem and AD-ND (0.80[0.41-1.54], p=0.5; Fig.-1B).

**Fig.-1.**
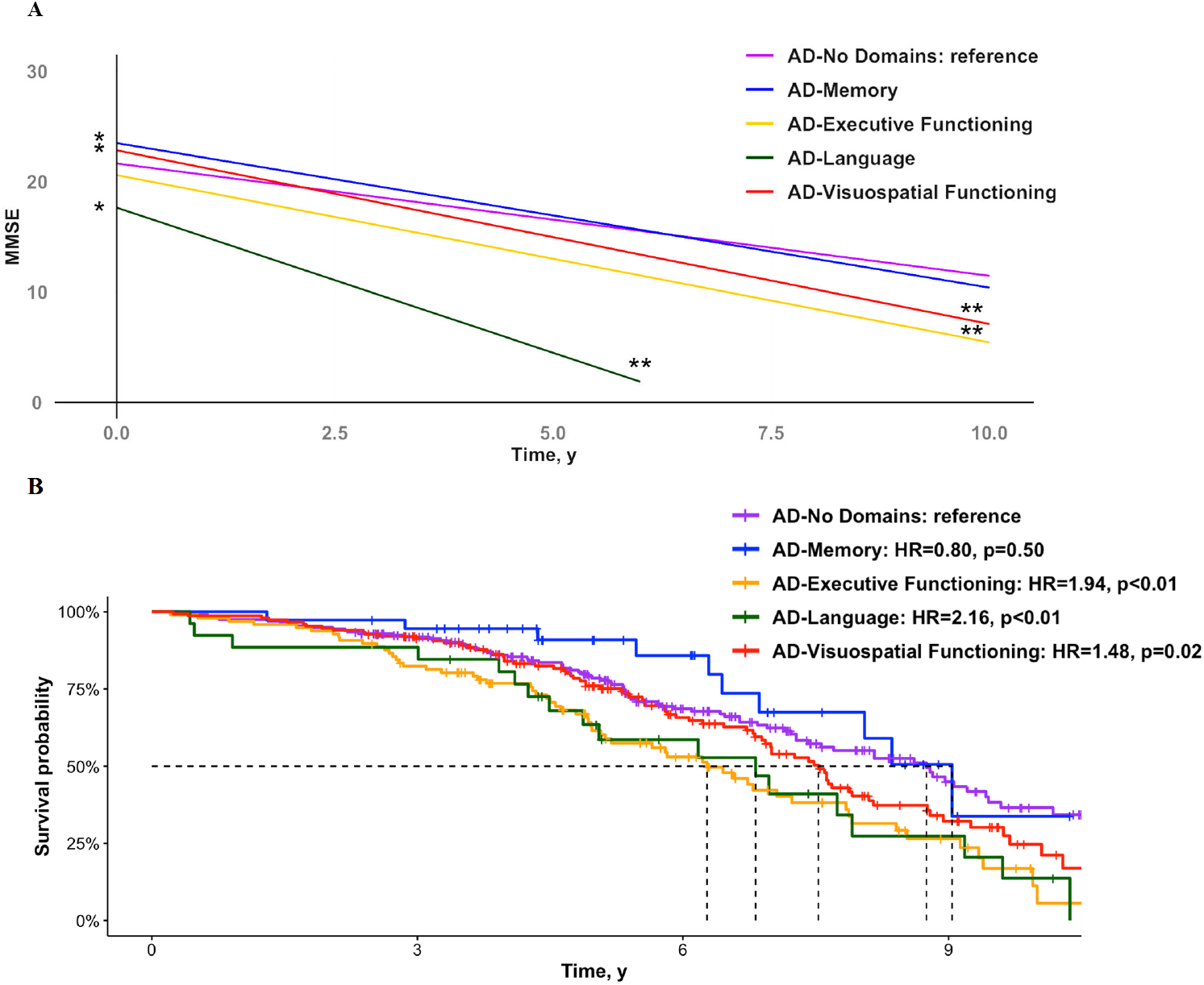
Clinical progression and mortality rates across cognitively-defined AD-subgroups. The plot in panel A displays results from linear-mixed effects models assessing the effect of subgroup and the interaction effect between subgroup*time on MMSE scores, adjusted for age, sex and education. The model includes one predictor for all subgroups with AD-ND as the reference. The Kaplan-Meier curves in panel B represents the survival probability over time for the various subgroups. Hazard ratios were calculated using a Cox proportional hazard model using AD-ND as the reference, and was adjusted for age. * - effect of subgroup on baseline MMSE, ** - interaction effect of subgroup*time on MMSE over time

### Regional patterns of atrophy

Voxel-wise contrasts between cognitively-defined AD subgroups and cognitively normal controls revealed prominent temporoparietal atrophy patterns across all subgroups (Fig.-2A), as well as subgroup-specific atrophy patterns. The latter atrophy patterns are best appreciated in the voxel-wise contrasts between AD-ND and the other AD-subgroups (Fig.-2B). Atrophy in the AD-Mem group was mainly localized in the medial temporal lobe – especially in bilateral hippocampus – while AD-Lang showed a temporal predominant atrophy pattern that was lateralized towards the left hemisphere. AD-Exec displayed a widespread pattern with greater cortical involvement compared to AD-ND. For AD-VS, the pattern of atrophy markedly occupied posterior brain areas. Atrophy patterns for the AD-ND subgroup showed a pattern characteristic for AD, encompassing the bilateral MTL, temporal and inferior parietal regions (Fig.-3A). This same pattern was seen in AD-Multiple (Supplemental-Fig.-1). Regional effects sizes (i.e. T-values) for the contrasts between AD-subgroups and cognitively normal controls are presented in the supplement (Supplemental-Fig.-2).

**Fig.-2.**
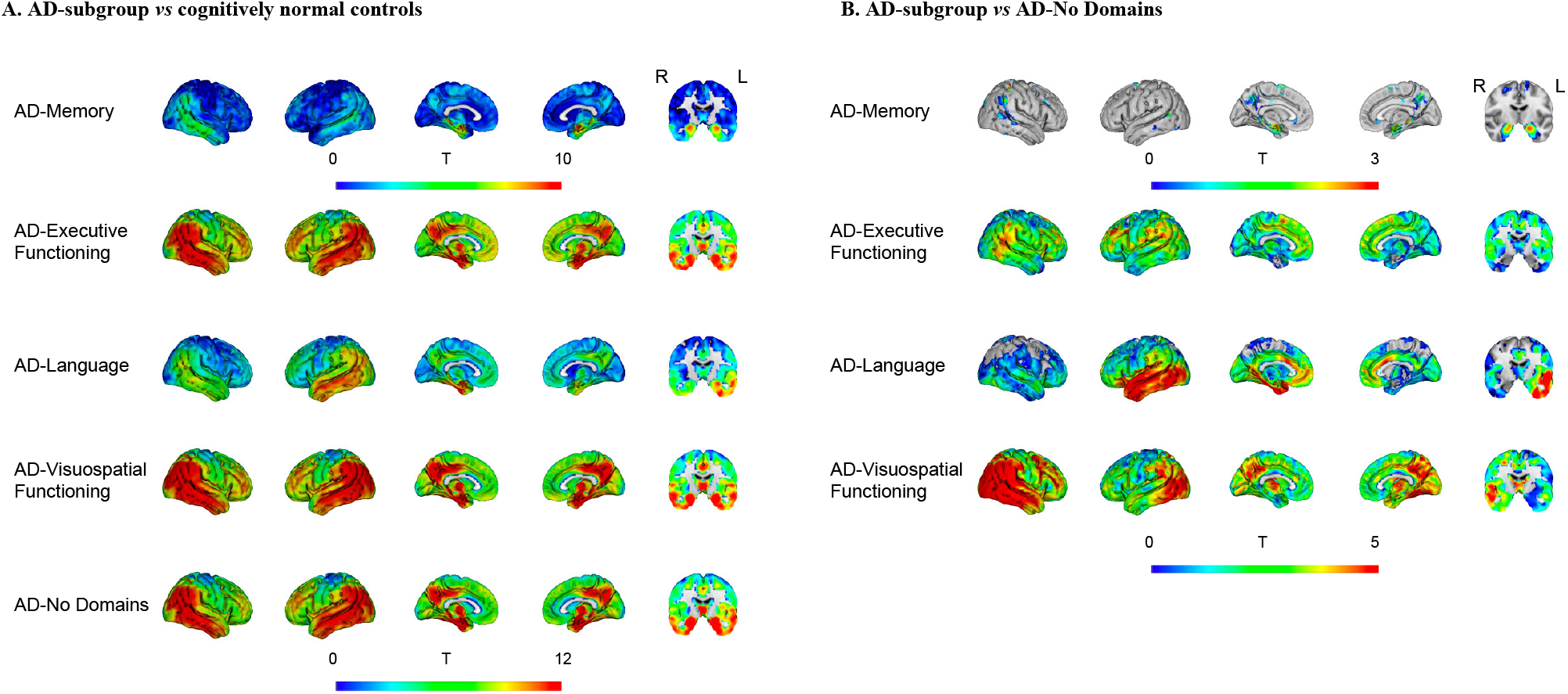
Regional atrophy across cognitively-defined AD-subgroups. Results from voxel-based morphometry analyses, displayed as T-maps, representing subgroup-differences in regional gray matter volumes, adjusted for age, sex, scanner, intracranial volume. Higher T-values indicate more atrophy. Panel A displays results from voxel-wise contrasts between subgroups and cognitively normal controls. Panel B displays results from voxel-wise contrasts between the domain-specific subgroups and the AD-ND group, which were additionally adjusted for whole brain gray matter to intracranial volume ratios (to adjust for differences in overall atrophy between subgroups). Voxel-wise contrasts showing only significant voxels are displayed in the supplement (Supplemental-Fig.-3). The T-values from the comparisons against controls in Panel A will be used as input for the neurogenetic analyses below.

### Regional patterns of atrophy in AD-subgroups vs atypical variants of AD

Given their clinical resemblances, we aimed to compare the spatial patterns of atrophy between AD-Lang and AD-VS and two positive-control reference groups, one of lvPPA participants and one of PCA participants (see *“Participants”* section). A striking similarity can be appreciated from Fig.-3A, which displays the voxel-wise contrasts *vs* cognitively normal controls (see Supplemental-Fig.-4 for a detailed depiction of atrophy patterns in the lvPPA and PCA reference groups). Spearman correlation analyses between regional gray matter atrophy for AD-Lang *vs* lvPPA (rho=0.92, p<0.01), and AD-VS *vs* PCA (rho=0.65, p<0.01) confirmed a high spatial correspondence (Fig.-3B). Furthermore, the Sørensen–Dice coefficient (DSC; see *“Statistical Analyses”* section) for overlap between the most atrophied voxels (Fig.-3C) for AD-Lang and lvPPA was 0.73, which far exceeded the DSC for overlap between lvPPA and all other groups (AD-Mem: 0.19, AD-Exec: 0.23, and AD-VS: 0.19) and also far exceeded the DSC of overlap between AD-Lang and the other subgroups (AD-Mem: 0.23, AD-Exec: 0.22, and AD-VS: 0.19). The DSC for overlap between AD-VS and PCA was 0.33, which exceeded the DSC for overlap between PCA and all other groups (AD-Mem: 0.04, AD-Exec: 0.17 and AD-Lang: 0.04; Fig.-5D) and was higher than the DSC for overlap between AD-VS and AD-Mem (0.27) and AD-Lang (0.19), though lower than DSC for overlap with AD-Exec (0.50; Fig.-5D).

**Fig.-3.**
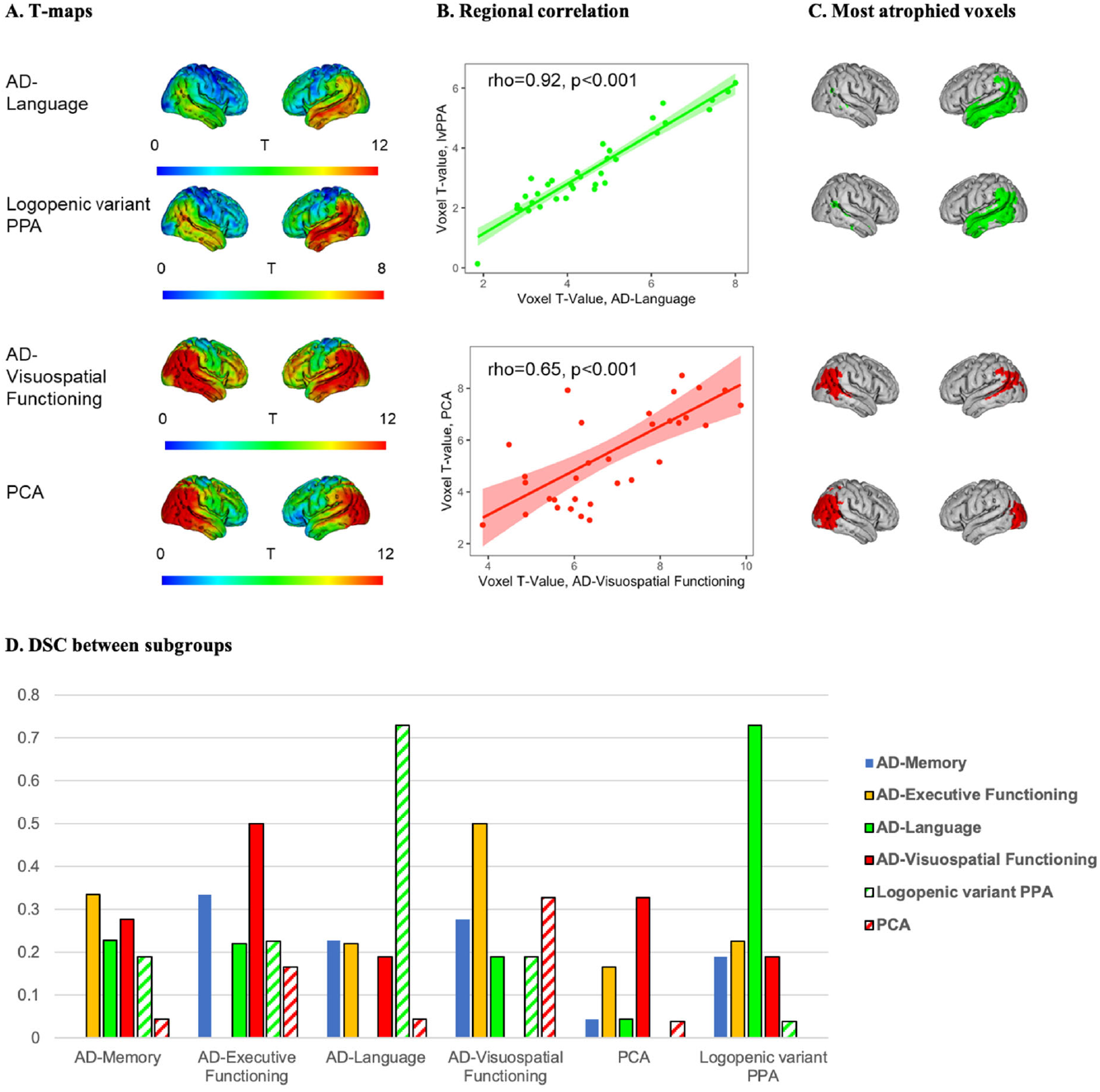
Similarity between AD-Language and lvPPA, and between AD-Visuospatial-Functioning and PCA. Panel A displays the voxel-wise contrasts for AD-Lang, lvPPA, AD-VS and PCA *vs* cognitively normal controls, adjusted for age, sex, scanner and intracranial volume. Panel B displays Spearman correlation between regional T values from the voxel-wise contrasts against cognitively normal controls for AD-Lang and lvPPA (top), and AD-VS and PCA (bottom). Panel C displays binarized maps of the most atrophied voxels for each T-map according to the threshold: (Meant + 2*SDt), with the mean as average within voxel-wise T-maps (contrasts against cognitively normal controls) and the SD the standard deviation within that image. Panel D displays the percentage overlap of the most atrophied voxels along with the Sørensen–Dice coefficient (DSC), which is used to assess similarity between groups and is calculated as: 2*(A⋂B)/(A+B). 0 indicates no overlap and 1 complete overlap.

### Neurogenetic profiles associated with subgroup-specific regional atrophy patterns

To examine whether the observed differences in regional atrophy associate with specific neurogenetic properties of the affected brain regions, we assessed the spatial correspondence of atrophy patterns with expression of candidate genes and genome-wide gene expression profiles from brains of controls with normal cognition, as provided by the Allen human brain atlas (Hawrylycz et al., 2015).

### Association between spatial pattern of atrophy and gene expression of candidate genes

For initial spatial association analyses we selected candidate genes based on their reported association with AD (Grothe et al., 2018; Jansen et al., 2019; Karch and Goate, 2015; Sepulcre et al., 2018). Results are summarized in Fig.-4. *BIN1* and *APOE* expression levels showed a consistent positive spatial correlation with regional atrophy across AD-subgroups, and *SORL1* and *ABCA7* expression levels showed a consistent negative association. The association with regional atrophy in PCA was in the opposite direction for these genes and the correlations with atrophy in AD-VS were consistently of lower magnitude compared to the other subgroups. Finally, *MAPT* gene expression was positively associated with atrophy across all AD-subgroups and *CLU* gene expression was negatively associated with atrophy across all AD-subgroups (Fig.-4).

**Fig.-4.**
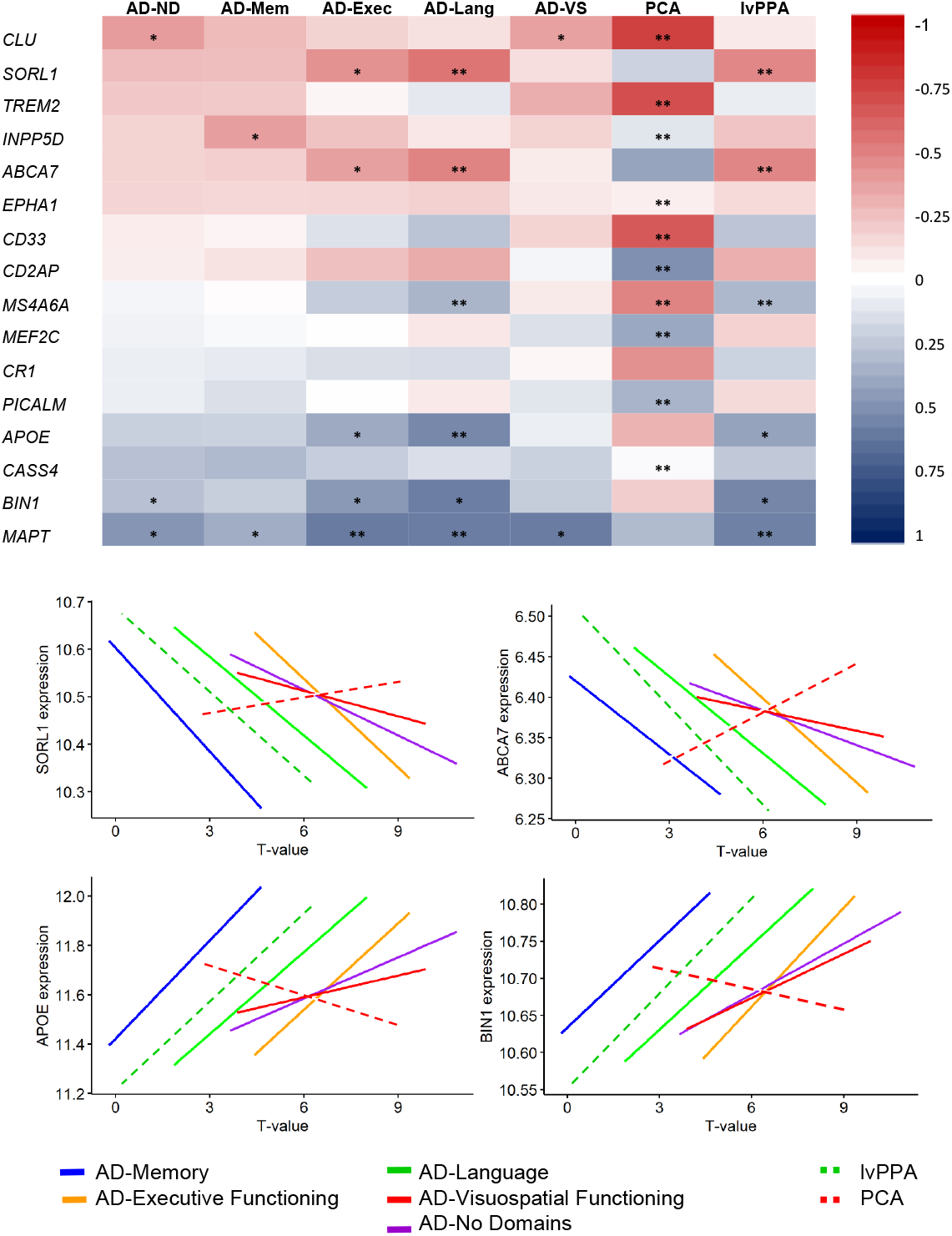
Spatial correlation between AD-subgroup specific atrophy and expression of candidate genes. The heatmap shows the association between regional atrophy and regional expression of genes previously associated with AD. Scatter plots highlight a selection of the spatial associations, with T-values of the subgroup-specific atrophy pattern (compared to cognitively normal controls) within the 34 Desikan-Killiany regions on the x-axis and gene expression obtained from the Allen human brain database within the same regions on the y-axis. The slopes show the Spearman correlations and individual data points are removed for visualization purposes. * - statistically significant Spearman’s correlation at p<0.05 (uncorrected), ** - statistically significant Spearman’s correlation at p<0.05 FDR-corrected

### Genetic pathways associated with atrophy patterns across multiple subgroups

To assess genetic pathways that might explain differences in subgroup-specific atrophy we performed exploratory gene-set enrichment analyses (GSEA, Broad Institute, Cambridge, MA, USA) using gene-expression data for all 18,686 genes provided by the Allen human brain atlas. Supplemental-Table-1 lists all enriched gene sets for the different subgroups that were identified through this approach and Fig.-5 summarizes their overlap across groups. Two positively enriched clusters associated with synaptic function and plasticity (e.g. dopamine release cycle, and long-term synaptic and neuronal plasticity) were shared between all subgroups. In AD-Mem and AD-VS, we observed a large negatively enriched cluster with gene sets implicated in mitochondrial respiration (e.g. ATP synthesis, respiratory electron transport), a cluster that was also present in AD-ND. Another large negatively enriched cluster present in multiple subgroups (AD-Mem, AD-Exec and AD-VS) comprised gene sets associated with protein metabolism (e.g. methylation, amino acetylation), which was also present in AD-ND. A smaller negatively enriched cluster associated with autophagy (e.g. mitophagy, mitochondrial depolarization) was present in AD-Mem and AD-ND, and another small cluster associated with the immune system (e.g. interleukin-7 and regulation of alpha/beta t-cell activation) was negatively enriched in AD-Mem, AD-VS and AD-ND, and positively enriched in AD-Exec and AD-Lang (Fig.-5; Supplemental-Fig.-5; Supplemental-Table-1).

### Genetic pathways uniquely associated with subgroup-specific atrophy patterns

For AD-Mem, three negatively enriched clusters were not shared with AD-ND or any of the other subgroups, indicating that these are uniquely implicated in AD-Mem. These clusters were associated with the cell cycle (e.g. DNA replication, regulation of apoptosis), ribosomal structure (e.g. ribosome assembly, small ribosomal subunit) and membrane proteins (e.g. MHC protein and cell lumen). Another small negatively enriched cluster associated with RNA metabolism (e.g. mRNA splicing, precatalytic spliceosome) was present only in AD-Mem, while this cluster was positively enriched in AD-Exec. For AD-Lang, two unique clusters were identified comprising negatively enriched gene sets associated with taste receptor activity and positively enriched gene sets associated with metabolism of proteins (e.g. genes associated with axon guidance and angiogenesis, and cytosolic ribosome), respectively. While other subgroups also showed enrichment for gene sets implicated in protein metabolism, these did not overlap with the AD-Lang specific gene sets and were also negatively rather than positively enriched. For AD-VS, a large negatively enriched cluster associated with modification of gene expression (e.g. epigenetic regulation and depurination), a smaller negatively enriched cluster associated with metabolism of carbohydrates (e.g. gluconeogenesis, lysosomal lumen), and a small positively enriched cluster of gene sets associated with keratinization were unique to this subgroup. There were no clusters for AD-ND that were unique to this group (Fig.-5; Supplemental-Fig.-5; Supplemental-Table-1).

**Fig.-5.**
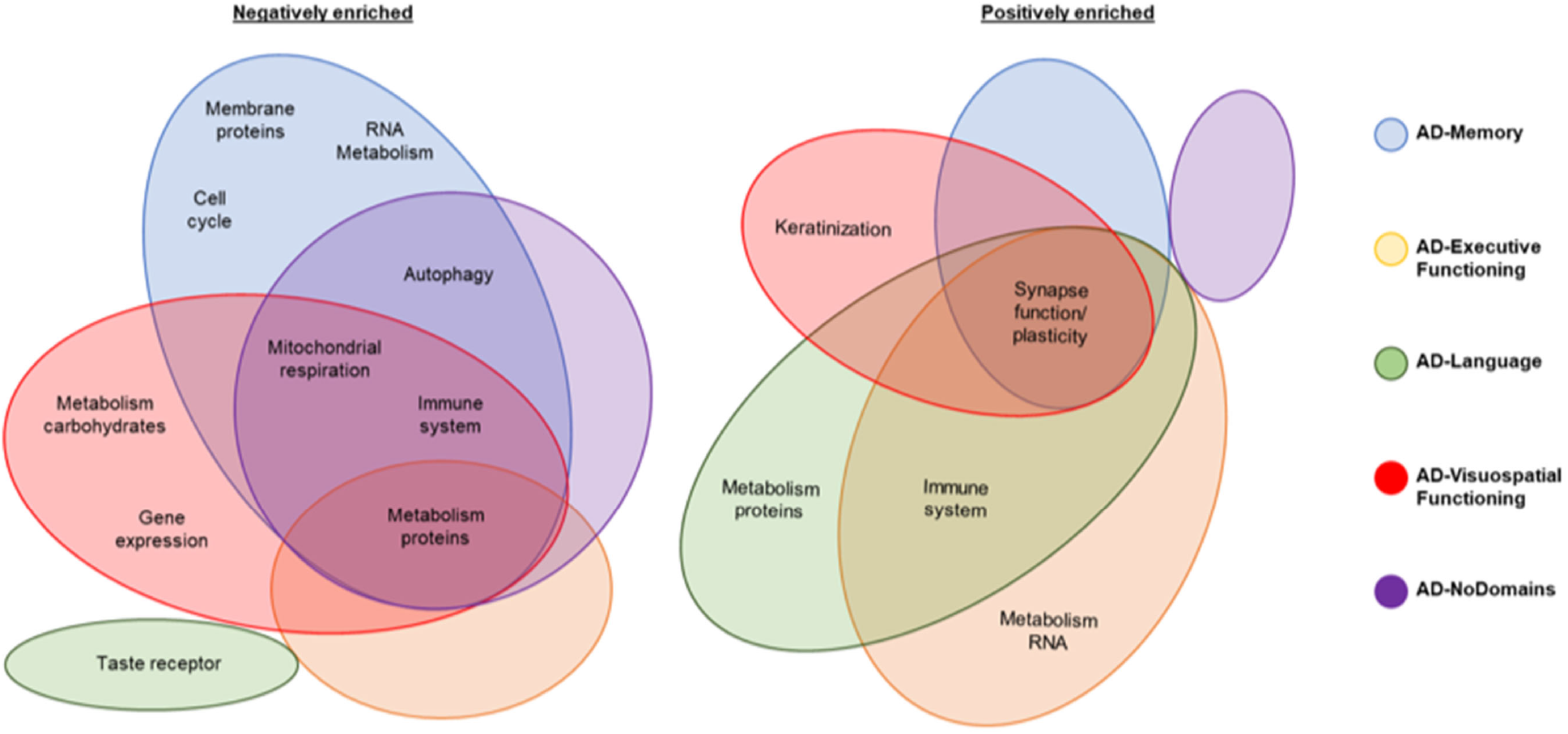
Overview of shared and unique gene set groups associated with atrophy patterns across AD-subgroups. Groups of gene sets named according to their shared biological function

## Discussion

We categorized 679 amyloid-β positive individuals clinically diagnosed with AD-type dementia into subgroups based on the distribution of impairments across four cognitive domains. We found that these cognitively-defined subgroups displayed distinct patterns of atrophy, indicating that cognitive heterogeneity is associated with differences in regional susceptibility to atrophy, even within the spectrum of typical AD. More specifically, we observed greater medial-temporal involvement in AD-Mem, more atrophy in medial-frontal/parietal regions in AD-Exec, left>right temporal atrophy in AD-Lang and greater involvement of posterior parts of the brain in AD-VS. The atrophy patterns of AD-Lang and AD-VS were highly analogous to those observed in lvPPA and PCA groups, suggesting that atypical variants and AD-subgroups are part of a clinical-radiological spectrum. To explore potential leads for neurogenetic properties that might underlie the observed clinical and neurobiological heterogeneity among subgroups, we associated regional atrophy patterns in AD subgroups with brain-wide gene expression profiles from individuals without neurological abnormalities provided by the Allen brain database of the human transcriptome. We found overlapping genetic pathways that were associated with atrophy patterns across multiple subgroups, including gene sets involved in metabolism of proteins, mitochondrial respiration, the immune system, and synaptic function/plasticity. There were also genetic pathways that were unique to specific cognitively-defined subgroups, including pathways involved in cell cycle and autophagy for the AD-Mem group, certain sets of protein metabolism in AD-Lang, and modification of gene expression in AD-VS. These findings point to distinct molecular factors involved in the pathogenesis of AD, which may aid the development of personalized medicine strategies.

### Clinical implications of cognitive heterogeneity

Compared to AD-ND, all subgroups except AD-Mem showed faster clinical disease progression and had a higher mortality rate, illustrating important clinical implications of AD subgroup membership. Previous examinations using the same cognitive-subtype classification in independent datasets demonstrated various clinical differences between subgroups, including fewer depressive symptoms in AD-Mem (Bauman et al., 2019), and - in line with our results - faster functional and cognitive decline in subgroups other than AD-Mem (Mez et al., 2013a, 2013b). Furthermore, the faster progression and higher mortality rates in subgroups other than AD-Mem is in accordance with previous examinations using other methods to classify subjects (Scheltens et al., 2018; Ten Kate et al., 2018). Taken together, these results suggest that subgroup categorization based on cognitive heterogeneity among individuals with AD holds relevance for an accurate diagnosis and prognosis, which, in turn, has important implications for patient wellbeing, as uncertainty about diagnosis and prognosis are often perceived as highly burdensome.

### Associations between genetics and clinical phenotype in Alzheimer’s disease

The mechanisms underlying clinical-biological heterogeneity in AD are still largely unknown. However, previous examinations have revealed that there are specific genetic risk factors that influence the clinical manifestation of AD. For instance, it is has repeatedly been shown that *APOEε4* carriers have more extensive medial temporal atrophy and memory deficits, while *APOEε4* negative individuals are predisposed to vulnerability of cerebral networks beyond the medial temporal lobe and consequently show mainly impairments in domains other than memory (van der Flier et al., 2011). In line with these findings, the prevalence of *APOEε4* in our AD-Mem group was a striking 90%, which is substantially higher than for all other AD-subgroups in our study and also compared to what is typically reported in AD cohort studies (66% (Mattsson et al., 2018)). It also replicates a multi-cohort study (Mukherjee et al., 2018) showing that higher proportions of people with AD-Mem had ≧1 *APOEε4* allele than the average for the study. The reason for this specific association between *APOEε4* and AD-Mem is not known but is strikingly consistent across samples, including specialty clinic settings (e.g. Amsterdam UMC Alzheimer center, the Alzheimer Disease Neuroimaging Initiative [ADNI], University of Pittsburgh Alzheimer’s Disease Research Center) and prospective cohort studies (e.g. Adult Changes in Thought [ACT] study, Religious Order Study [ROS], and the Rush Memory and Aging Project [MAP]) (Crane et al., 2017b; Mukherjee et al., 2018, 2012; Van Der Flier et al., 2014). This consistency highlights *APOEε4* positivity as an important determinant to develop a memory-predominant clinical presentation of AD.

In addition to the usual suspect in AD research, the *APOE* gene, there is a range of other genes that have been associated with AD risk (Grothe et al., 2018; Jansen et al., 2019; Sepulcre et al., 2018). In previously published genetic association studies using the same subgroup classification scheme, genetic loci previously associated with AD subgroup-specific associations showed varying odds-ratios (Crane et al., 2017a) and novel loci were specific for different subgroups (Mukherjee et al., 2018). The discovery of subgroup-specific novel genetic loci is still in its infancy and examinations of previously identified loci and genes may be instrumental in determining potential therapeutic targets and personalized medicine strategies. However, these approaches are reliant on large sample sizes and require additional fine mapping because top hits are often found outside of coding regions and are very rarely the causal SNP.

### Spatial correlation between expression of known risk genes and atrophy

To better understand how genetic risk factors may relate to differential regional susceptibility to atrophy in AD, we examined whether region-specific expression levels of genes located within known AD risk loci were associated with the differential atrophy patterns observed across AD-subgroups. We found that spatial correlations between regional atrophy and expression of *BIN1* and *APOE* (top AD risk genes) were weaker for AD-VS than for the other subgroups and even in the opposite direction for PCA. *MAPT* gene expression was positively associated with atrophy across all AD-subgroups and reference groups, indicating that *MAPT* is associated with atrophy regardless of clinical phenotype. This is in line with two previous studies demonstrating consistent regional associations of *MAPT* expression levels with the typical (averaged) atrophy pattern in AD dementia patients as well as with the cortical spreading pattern of tau pathology as determined from longitudinal tau-PET assessments in older controls (Grothe et al., 2018; Sepulcre et al., 2018).

### Brain-wide gene expression profiles associated to atrophy across subgroups

We also evaluated gene-expression driven biological pathways associated with regional atrophy patterns across subgroups. We used a brain imaging-gene expression co-localization approach that has recently shown promise in identifying potential novel candidate genes and biological pathways implicated in regional susceptibility to pathologic alterations in AD and other neurodegenerative diseases (Freeze et al., 2019; Grothe et al., 2018; Sepulcre et al., 2018). This approach utilizes the Allen Brain Database of the Human Transcriptome, a publicly available resource which provides stereotactically characterized expression profiles of cognitively normal control subjects and allows the assessment of spatial correlations between brain imaging findings and regional gene expression characteristics (Arnatkevičiūtė et al., 2019). We used this method in combination with an exploratory gene set enrichment approach (GSEA) approach, which accounts for the high-dimensionality of the transcriptome-wide gene expression data by searching for functional genetic pathways robustly associated with the respective atrophy pattern across several implicated genes. We observed that genetic pathways that are negatively enriched in regions that show atrophy (i.e. lower expression in regions with more atrophy) across multiple subgroups could be roughly classified into mitochondrial respiration, metabolism of proteins and the immune system. Mitochondrial dysfunction is thought to be (potentially causally (Swerdlow et al., 2014)) implicated in the pathogenesis of AD (Flannery and Trushina, 2019), possibly through an interaction with *APOE* (Yin et al., 2020). Our findings regarding the association between atrophy and lower expression of genes associated with mitochondrial respiration could be due to increased susceptibility in these regions to mitochondrial dysfunction, subsequent oxidative stress, and eventual cell-death and atrophy. With regard to gene sets associated with protein metabolism, ribosomal protein synthesis is lower in brain tissue that is affected by AD (Langstrom et al., 1989), which is in line with our results. This process has usually been regarded as a downstream consequence of pathology rather than an upstream process (Langstrom et al., 1989), but points to another possible avenue for intervention in AD. Interestingly, leading edge analyses of the most relevant genes driving the enrichment signal for these gene sets pointed to genes from the mitochondrial ribosomal protein (MRP*) family (see Supplemental-Fig.-6), which has previously been proposed as a target to combat mitochondrial dysfunction (Sylvester et al., 2004). Another genetic pathway that was implicated in multiple groups was immune function. We found that atrophy in AD-Mem, AD-Exec and AD-ND was associated with lower expression of genes associated with interleukin-7, a cytokine involved in T-cell development. These findings are in line with a previous study showing that genes involved in immune response are negatively enriched in regions vulnerable to AD pathology (Freer et al., 2016) and supports a key role of inflammation in AD pathogenesis (Gjoneska et al., 2015).

However, in contrast to these findings, we also observed that expression of genes associated with regulation of T-cell activation and differentiation was positively enriched in areas with atrophy in brain regions associated with AD-Lang and AD-Exec. Furthermore, we observed that genes associated with synaptic function/plasticity were also enriched in the regions with more atrophy across all AD-subgroups, which replicates a previous examination in an independent sample of AD patients (Grothe et al., 2018). In AD, the spatial pattern of neurodegeneration closely matched regional distributions of tau-pathology (Braak and Braak, 1991; Whitwell et al., 2012), and the most plastic brain regions, such as the medial temporal lobe (Gonçalves et al., 2016), also appear to be the most vulnerable to the initial deposition of tau pathology (Braak and Braak, 1991; Walhovd et al., 2016). The correspondence of regions with both heightened synaptic plasticity and atrophy therefore becomes apparent. Initially, plastic brain regions would be assumed to be the most resistant to pathology and neurodegeneration, but in the long-term this may become maladaptive as the brain ages and pathology sets in (Hillary and Grafman, 2017). The association between regional susceptibility to neurodegeneration and enrichment of genetic pathways associated with synaptic plasticity is therefore consistent with a selective vulnerability of plastic brain regions like the medial temporal lobe. While in the present study, AD-Mem showed the most pronounced medial temporal atrophy, the other subgroups were still characterized by an AD-characteristic pattern with atrophy including the medial temporal lobe in comparison with cognitively normal controls, which might explain why this cluster was observed across multiple subgroups.

### Unique gene-expression profiles underlying clinical-radiological heterogeneity between AD subgroups

In addition to overlapping genetic pathways across multiple subgroups, we also observed clusters of gene sets that were uniquely associated with a single subgroup. For AD-Mem, we found a large cluster of negatively enriched gene sets associated with the cell-cycle. Associations between the cell-cycle and AD have been demonstrated before and involves the dysfunction of neuronal cell-cycle reentry leading to the two-hit hypothesis. The first hit involves dysfunctional cycle reentry, which would normally result in apoptosis and no development of AD pathology. However, chronic oxidative stress can cause a second hit that prevents normal apoptosis and allows the build-up of AD pathology (Moh et al., 2011). More research is necessary to determine why this mechanism would be more pronounced in AD-Mem than in other subgroups. For AD-Lang, we found that gene sets associated with metabolism of proteins were positively enriched in regions that showed the most atrophy in this subgroup, while in the other subgroups gene sets associated with metabolism of proteins were negatively enriched in the most atrophied regions. Furthermore, while the gene sets were related to overlapping biological functions, the specific gene sets implicated in AD-Lang were different form the ones implicated in the other subgroups, and analysis of the leading-edge genes in the cluster for AD-Lang pointed to the ribosomal protein family (RP*) rather than the mitochondrial ribosomal protein family (MRP*) like in the other subgroups (Supplemental-Fig.-6). The differential implication of a range of gene sets related to the same biologic process is intricate, and it remains to be determined whether and how these differences might contribute to differential regional susceptibility to atrophy across subtypes. While given the current state of research it is hard to establish a causative role between higher ribosomal protein expression and AD-Lang specific atrophy (i.e. left-lateralized temporal), this finding does indicate that potential therapeutic approaches aimed at modulating expression of ribosomal proteins might not be beneficial to all AD patients and may even be detrimental in some cases (Caccamo et al., 2015). We also found a negatively enriched cluster of gene sets that was unique to AD-Lang and associated with (bitter) taste receptors. While this association was present in the lvPPA group, and loss of taste and dysregulation of taste receptor proteins has been associated with AD before (Ansoleaga et al., 2013), there is currently no clear evidence on how this might be particularly associated with the characteristic left-lateralized temporal atrophy and language impairment. The function of taste receptor proteins within the brain is still largely unknown (Ansoleaga et al., 2013) and the gustatory cortex (anterior insula and frontal operculum) does not encompass any regions that were particularly implicated in AD-Lang. In AD-VS, we found a rather large, unique cluster of negatively enriched gene sets associated with gene expression modification. Gene sets within this cluster were mainly associated with epigenetic modifications (e.g. methylation, acetylation) and enrichment of this gene set cluster was mainly driven by the histone cluster protein (HIST*) gene family (Supplemental-Fig.-6), which is associated with packaging and ordering DNA into nucleosomes (Esposito and Sherr, 2019). Previous studies have shown that gene expression modification is implicated in AD through what is called an epigenetic blockade, referring to a large-scale decline in gene expression that is affected by post-translational histone modification (Gräff et al., 2012). Studies in mouse models have shown that this epigenetic blockade might be reversible (Gräff et al., 2012), opening up potential targets for therapeutic interventions. This epigenetic modification has been previously linked to an AD phenotype in animal models (Kosik et al., 2012), but there is no previous evidence that this points to a specific relation with visuospatial impairments and posterior atrophy. However, a recent GWAS study examining genetic loci associated with regional gray matter volumes (van der Lee et al., 2019) reported a distinct locus (rs12411216) specifically associated with occipital volumes. This locus is located in an intron of *MIR92B* and *THBS3,* showing a signal peak covering more than 20 genes with promotor histone marks overlapping the variant. This is an intriguing similarity to our findings and warrants further investigation. Another unique cluster of positively enriched gene sets found in AD-VS were related to keratinization. Keratin is most notably associated with fibrous structural protein that makes up hair and nails and its function in the context of the neuronal milieu is not well known but may be associated with mechanical stability of cells (Moll et al., 2008). How this might relate to susceptibility to posterior cortical atrophy and related visuospatial impairments is unknown at this point.

Taken together, our results regarding neurogenetic factors associated with the distinct atrophy patterns across subgroups revealed several distinct clusters of biologically coherent gene sets, some of which showed unique associations with subgroup-specific atrophy patterns. Of note, several of the identified genetic pathways have been previously implicated in AD through diverse lines of genetic and molecular research. We further add to these findings by showing that expression levels of genes associated with these pathways are spatially linked to regionspecific brain atrophy, and we provide several potential molecular targets for future investigations on therapeutic interventions. By outlining that certain genetic pathways are uniquely implicated in specific cognitively-defined subgroups, we show that not all therapeutic targets might be equally beneficial for everybody with AD and highlight the need for examinations into personalized medicine strategies.

### Strengths and limitations

Strengths of the study include a relatively large sample size from a single center with consistent assessments of AD biomarker positivity and well phenotyped patients with dementia who had 3Tesla MRI available, and the integration of knowledge from multiple sources (i.e. clinical, imaging and genetics). Furthermore, in contrast to previous examinations that focused on categorization of individuals with AD based on structural properties (Ossenkoppele et al., 2019; Risacher et al., 2017; Ten Kate et al., 2018; Zhang et al., 2016), structural properties in combination with cognition (Sun et al., 2019), neuropathological features (Murray et al., 2011; Whitwell et al., 2012), or clustering analyses (Scheltens et al., 2017; Stopford et al., 2008) and factor scoring (Sevush et al., 2003) of cognitive data, we used a classification scheme that relies on the intra-individual distribution of impairments across cognitive domains. This relatively simple method relies on patientspecific profiles of impairments across cognitive domains and can be performed on an individual basis, which is in contrast to approaches such as clustering analyses that rely on large sample sizes and sufficiently distributed data. Our study also has several limitations. First, the relative group sizes of the cognitively-defined subgroups presented in this work may not be representative of group sizes in other cohorts. The source of our sample was the Amsterdam Dementia cohort, which is a cohort from a tertiary memory clinic which specializes in, and is therefore enriched for, early onset and atypical AD clinical presentations (Van Der Flier et al., 2014). While this might limit the direct generalizability of our findings to other cohorts, and replication in more clinically representative cohorts is needed, the variety in clinical profiles in the ADC cohort allowed us to obtain sufficiently large subgroup sizes to assess differences in clinical progression and to obtain robust estimates of the differential regional susceptibilities to atrophy and their related gene expression profiles. Furthermore, we have outlined results from stratified groups of EOAD and LOAD in the supplement (Supplemental-Tables 2-3; Supplemental-Fig.-7-8), and show that, although atrophy is generally more pronounced in EOAD, the spatial pattern differences across subgroups is very similar to what we see in LOAD. Another potential limitation of the study is that we did not account for comorbidities, which might be present to varying degrees across these subgroups. For instance, white matter changes are particularly related to impairments in executive processing and may thus potentially be overrepresented in the AD-Exec subgroup (Prins and Scheltens, 2015). Other limitations of the present study include the absence of information on cause of death in the mortality data, the difficulty in assessing specific characteristics of individuals in the heterogeneous AD-Multiple subgroup, and that the Allen Human Brain database only contains gene-expression values from six younger subjects with limited bi-hemispheric data (Hawrylycz et al., 2012), which prevented us from assessing neurogenetic properties related to lateralization of atrophy. This may particularly be an issue for the AD-Lang subgroup, as the atrophy pattern in this subgroup showed a marked hemispheric asymmetry.

### Rethinking the definition of typical Alzheimer’s disease

Elucidating the associations between clinical and neurobiological heterogeneity in AD is crucial in understanding pathogenesis and developing effective treatments. Complementary to previous findings (Bauman et al., 2019; Crane et al., 2017a; Mez et al., 2013a, 2013b; Mukherjee et al., 2018), we show that cognitively-defined non-amnestic AD subgroups are associated with i) a faster clinical progression, ii) higher mortality rates, and iii) subgroup-specific regional atrophy patterns, which are iv) related to different genetic pathways. These insights into the heterogeneity among AD patients are testament to the considerable variability within the spectrum of the disease, and meaningful stratification into distinct subgroups based on cognitive data might prove useful in future diagnostic and prognostic work-ups, and may even result in personalized medicine strategies. Although none of the participants in our sample fulfilled diagnostic criteria for recognized atypical variants of AD, our cognitively-defined subgroups were both clinically and radiologically reminiscent of these phenotypic extremes (Crutch et al., 2017; Gorno-Tempini et al., 2008; Madhavan et al., 2013). In line with observations in previous work (Snowden et al., 2007; Stopford et al., 2008), we would therefore propose that phenotypic presentations of AD are more accurately arrayed along a clinical-radiological spectrum. Atypical presentations are increasingly recognized and included in diagnostic criteria. However, the vast majority of AD patients do not meet the rather strict clinical criteria for an atypical variant of AD and are therefore, by default, regarded as typical AD. We would propose that future diagnostic criteria should also account for the considerable heterogeneity among these individuals (Fig.-6). This hypothetical model is a work in progress and more research is needed to map the clinical and neurobiological heterogeneity in AD in all its complexity into one model. For instance, it is necessary to properly identify and define a distinctive selective amnestic variant of AD, if it exists. Findings concerning the clinical (slower progression and less depressive symptoms (Bauman et al., 2019; Mez et al., 2013b, 2013a)), neurodegenerative (regionally-specific MTL atrophy ((Lam et al., 2013b)), and genetic characteristics (higher proportion of *APOEε4* (Crane et al., 2017a; Mukherjee et al., 2018) and partly unique atrophy-related gene expression profile) all indicate that there is a distinction between memory-predominant (amnestic) AD and the typical clinical presentation with heterogeneous impairments across domains that is most often observed in AD (such as seen in our AD-ND group). The degree to which this amnestic variant overlaps with the AD-Mem subgroup defined by our approach is currently unclear, which is why we indicated the different categories in Fig.-6 with a dashed line. Largely the same arguments hold true for the relationship between the AD-Exec subgroup of our study and the dysexecutive variant of AD (Dickerson and Wolk, 2011; Ossenkoppele et al., 2015c), for which recently provisional research criteria were proposed (Townley et al., n.d.). Furthermore, the framework we have developed emphasizes patterns of cognitive functioning at the time of AD diagnosis, and ignores behavioral and personality changes. We suspect there may be a behavioral variant where behavioral aspects are more prominent than expected for the overall level of clinical impairment. While previous research has tried to delineate such a behavioral variant of AD (Dubois et al., 2007; Ossenkoppele et al., 2015c), this is difficult to study as behavior is rarely assessed as comprehensively as cognition in the research evaluation of people with newly diagnosed AD. Furthermore, it has yet to be determined whether a possible behavioral variant can be distinguished from a dysexecutive variant (Ossenkoppele et al., 2015b; Townley et al., n.d.). We again denote these uncertainties with dashed lines in our model, and highlight this as a potential area for future investigation. Future research will need to continue outlining the clinical and neurobiological disparities within the spectrum of AD (e.g. by mapping tau (Braak and Braak, 1991; Whitwell et al., 2008) and amyloid-β (Lehmann et al., 2013) pathology) and to examine other factors that might be involved in their emergence (e.g. pre-morbid learning disabilities (Miller et al., 2018, 2013) and structural properties of the pre-morbid brain (Batouli et al., 2014)). These efforts will advance the ongoing quest to answer fundamental questions about the mechanisms that are involved in the etiology of AD and the emergence of clinical and radiological heterogeneity among individuals with AD.

**Fig.-6.**
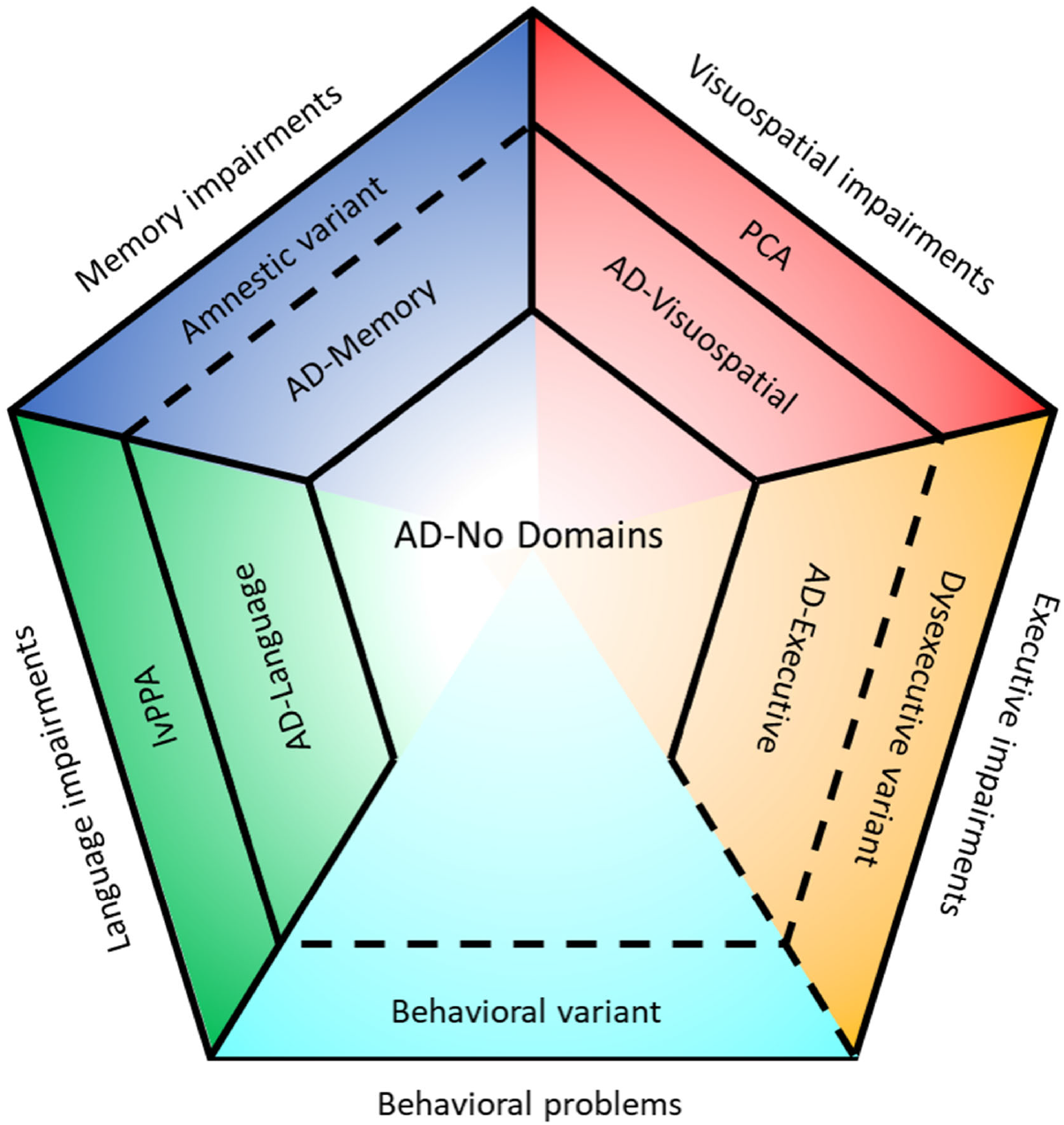
Hypothetical model of the Alzheimer’s disease clinical-neurobiological spectrum. Solid lines represent differences in categories that are either outlined in this manuscript or provided by established clinical criteria (i.e. for lvPPA and PCA). Dashed lines represent suspected differences in categories that are not yet established and are under investigation or need to be assessed in new lines of research. Note that relative sizes of the partitions are not representative for prevalence of the categories.

## Materials and Methods

### Participants

For this single-center, case-control study, we included participants from the Amsterdam Dementia Cohort (ADC; a memory clinic population at Amsterdam UMC) (Van Der Flier et al., 2014). Participants that would be categorized into AD-subgroups (see *“Subgroup classification”* section) were selected based on the following criteria: 1) clinical diagnosis of AD dementia (either EOAD or LOAD) (McKhann et al., 2011) at time of dementia screening, 2) molecular biomarker profile indicative of AD neuropathology (i.e. annual upward drift corrected CSF amyloid-β42<813pg/mL (Tijms et al., 2018); and/or a positive Aβ [^18^F]flutemetamol, [^18^F]florbetaben, [^18^F]florbetapir, or [^11^C]Pittsburgh compound-B] PET scan determined by visual assessment (Ossenkoppele et al., 2015a)), 3) availability of a 3Tesla MRI scan, and 4) availability of neuropsychological data to compute all cognitive domains assessed (see *“Cognition”* and *“Subgroup classification”* sections). Exclusion criteria for participants that were categorized into AD-subgroups were: 1) meeting core criteria for an atypical variant of AD, i.e. PCA or lvPPA (Crutch et al., 2017; Gorno-Tempini et al., 2008), 2) psychiatric or neurological disorders (other than AD), and 3) known genetic mutations associated with familial AD. A total of 679 participants were included based on these criteria. We selected a control group of cognitively normal subjects who attended our memory clinic but were found to have negative amyloid-β biomarkers and no objectifiable cognitive impairments (n=127, age 58±9, 42% male, MMSE 29±1). To assess how the cognitively-defined AD subgroups relate to established atypical variants of AD, we additionally selected “positive control” samples of individuals with lvPPA (n=20, age 66.9±5.2, 60% male, MMSE 23.1±4.1) and PCA (n=69, age 62.0±6.1, 41% male, MMSE 20.2±4.6) from our previous studies (Bergeron et al., 2018; Groot et al., 2020), which were diagnosed according to published clinical criteria (Crutch et al., 2017; Gorno-Tempini et al., 2008). There are currently no consensus diagnostic criteria for selective amnestic or behavioral/dysexecutive variants of AD. Written informed consent was obtained from all participants, and the local medical ethics review committee of the Amsterdam UMC approved the study.

### Cognition

Neuropsychological test scores were categorized into four cognitive domains; memory, executive-functioning, language and visuospatial-functioning. Distribution of tests for each domain is provided in the supplement (Supplemental-Table-4). Confirmatory factor analyses using Mplus software (version 7.419) (Muthén and Muthén, 1998) were implemented to generate composite scores for these four cognitive domains. Domain specific composite scores were then co-calibrated to a metric based on the distribution of scores among 4,050 people with incident AD from the Adult Changes in Thought (ACT) study, the Alzheimer’s Disease Neuroimaging Initiative (ADNI), the Rush Memory and Aging Project (MAP) and Religious Orders Study (ROS), and the University of Pittsburgh Alzheimer Disease Research Center (PITT). These four LOAD (age >60 years) studies formed our legacy cohorts. Brief descriptions of each of the cohorts and detailed methods on how co-calibration of scores was achieved can be found in the supplement (Supplemental-Text 1) and our previous publications (Crane et al., 2017a; Mukherjee et al., 2018). Briefly, we identified “anchor” items with the same stimulus, and which were scored identically, in the ADC and the legacy cohorts. These items were used to anchor the domain specific composites across cohorts. We then used bifactor models in Mplus to co-calibrate scores in the ADC LOAD (age >60 years) cohort to those in the legacy cohorts, for each of cognitive domains. We used the ACT sample of people with incident Alzheimer’s disease as our reference population for the purpose of scaling domain scores. We z-transformed scores from other studies to ACT-defined metrics for each domain. The scores of the ADC EOAD (<60 years) subjects were then co-calibrated using the item parameters from the ADC LOAD model. The ACT-normalized cognitive scores of the sample from the ADC cohort are presented in Supplemental-Table-5.

### Subgroup classification

As previously published (Crane et al., 2017a), subgroup classification relies on scores across all four domains assessed (i.e. memory, executive-functioning, language and visuospatial-functioning). Classification is achieved by first averaging, for each individual, the scores across the four cognitive domains and then determining the difference for every domain score from that average. We used a difference of 0.80 units to identify domains with scores substantially lower than a person’s average score. This threshold was previously empirically determined after assessing a range of candidate thresholds (see (Crane et al., 2017a) for further details). We then considered the number of domains with scores substantially lower than the person’s average score to assign people to groups (see Supplemental-Fig.-9 for a visual representation of categorization). This classification yields 6 groups. Consistent with our previous publications on these subgroups (Crane et al., 2017b; Mukherjee et al., 2018), these were named according to which domain showed substantial relative impairment: AD-Mem, AD-Exec, AD-Lang, AD-VS, AD-Multiple (more than one domain relatively impaired) and AD-ND (none of the domains relatively impaired).

### Clinical disease progression and mortality rates

Clinical disease progression was assessed by the Mini-Mental State Examination (MMSE), which is a measure of global cognition often used to track clinical disease progression. Median (range) number of MMSE assessments across subjects was 2 (1-9) and mean (SD) follow-up time was 1.7 (1.8) years. Furthermore, in the majority of our patients (578/679; 85%), data on mortality were available, which we obtained from the national municipal population register. Of the 679 individuals in our baseline sample, 286 individuals (49.5% of the sample with mortality data) were deceased at a mean follow-up of 5.7 (2.5) years.

### Neuroimaging

MRI scans were performed according to standardized acquisition protocols including a 3D T1-weighted structural MRI sequence with nearly isotropic (±1mm) voxel size acquired on three different 3Tesla MRI scanners (Supplemental-Table-6). We adjusted for scanner type in the statistical models (see “Statistical analysis” section). A standard SPM12 (Wellcome Trust Centre for Neuroimaging, UCL, London, UK) processing pipeline was used to segment and normalize MRI images to MNI space (Groot et al., 2018). The resulting gray matter density images were used as input for a voxel-based morphometry analyses assessing the spatial distribution of atrophy within each subgroup (see *“Statistical analysis”* section).

### Regional gene expression profiles

Brain-wide gene expression profiles were obtained from microarray data from the Allen Database of the human brain transcriptome, which is publicly available from the Allen brain institute (Seattle, WA, USA; http://human.brain-map.org). This dataset contains regional gene expression data from around 61.000 microarray probes collected from ~3700 tissue samples, which were obtained from six control subjects who had died without any evidence of neurologic disease (aged 24-57). Anatomical information for each of the probes is provided and can be used to determine the location within stereotactic standard space (MNI). Microarray data from the ~3700 tissue samples, with their corresponding anatomical locations, have been previously used to interpolate brain-wide gene-specific expression scores at the voxel level based on relative distances between samples using spatial variogram modelling (see (Gryglewski et al., 2018) for details). These brain-wide gene expression maps have been made publicly available at www.meduniwien.ac.at/neuroimaging/mRNA.html, and were parcellated for our study into 34 cortical regions-of-interest (ROI) as defined in the Desikan-Killiany neuroanatomical atlas (French and Paus, 2015). This resulted in a final dataset that includes gene expression data on 18,686 protein-coding human genes within each of the 34 ROIs covering the entire cerebral cortex. While initial analyses from the Allen human brain project focused on microarray data from both hemispheres, this approach was abandoned after two subjects were deemed to have no inter-hemispheric differences in gene expression (Hawrylycz et al., 2012). Gene expression analyses in the other four donors was limited to the left-hemisphere only. Given the paucity of data from both hemispheres, and in line with previous examinations (Grothe et al., 2018; Sepulcre et al., 2018), we restricted all our analyses assessing associations between atrophy and gene-expression to the left hemisphere. In an initial candidate gene analysis, we assessed the spatial correlation between atrophy in each of the subgroups (see *“Statistical analysis”* section) and expression of genes that have previously been associated with AD risk (Grothe et al., 2018; Jansen et al., 2019; Karch and Goate, 2015; Sepulcre et al., 2018) (see Fig.-4 for an overview of these genes). Then, the relationship between regional atrophy patterns and brain-wide gene expression profiles (including all 18,686 genes) was assessed using spatial association analyses and subsequent explorative gene set enrichment analysis (GSEA) of the associated genome-wide expression profiles (Grothe et al., 2018; Subramanian et al., 2005). Identified gene sets were further grouped into clusters of related gene sets and organized into a network structure (Merico et al., 2011) (see the *“Statistical analysis”* section for further details). We then examined and described groups of gene sets (clusters) shared with AD-ND or the other subgroups, as well as those that were uniquely enriched in one subgroup. A cluster was deemed uniquely implicated in one subgroup if the majority of the gene sets comprising that cluster were not shared with AD-ND or the other subgroups.

### Statistical analysis

All statistical analyses were performed in SPM12 or R version 3.5.2. Differences in demographic and clinical characteristics between the subgroups were assessed using independent-samples t-tests (continuous variables), χ^2^-tests (categorical variables) and Kruskal-Wallis tests (for the non-normally distributed education variable).

### Clinical disease progression and mortality

Linear-mixed effects models were assessed to examine differences between subgroups in MMSE scores at baseline and their change over time. We ran a model using one predictor for all subgroups, with AD-ND as the reference and adjusted for age, sex and education. To assess the difference in rates of mortality across cognitively-defined subgroups, age-adjusted Cox regression analyses were performed. Again, we ran one model comparing all subgroups to the AD-ND group as the reference. Statistical significance for linear mixed and Cox proportional hazard models was set at p<0.05.

### Spatial patterns of atrophy

To assess the brain-wide spatial pattern of atrophy for each of the groups, SPM12 was used to perform voxel-wise gray matter density contrasts between subgroups and amyloid-β negative, cognitively normal controls, as well as between the subgroups and the AD-ND subgroup. These analyses were adjusted for age, sex, intracranial volume and scanner type. We aimed to consider the possibility that one group may have presented later in the disease course on average than another group, leading to overall greater atrophy due to disease stage at time of diagnosis. Therefore, we included a term for overall atrophy (gray matter to intracranial volume ratio) in the contrast models for comparisons against the AD-ND group. All contrasts yield statistical parametric T-maps that represent the voxel-wise difference in gray matter volume between groups. The T-maps for the contrasts against cognitively normal controls were then used to assess spatial similarities between atrophy patterns among the cognitively-defined subgroups and the lvPPA and PCA positive-control reference groups, using Spearman correlations between regional atrophy in 34 regions of the Desikan-Killiany atlas. Additionally, we determined the percentage overlap between the most atrophied voxels from the T-maps (Meant + 2*SD_t_) and calculated the Sørensen–Dice coefficient (DSC) as: 2*(A⋂B)/(A+B), with 0 signifying no overlap and 1 complete overlap.

### Associations between atrophy and regional gene-expression profiles

For spatial association analyses with regional gene expression profiles, atrophy scores were extracted from the same 34 left-hemispheric cortical ROIs that were used for measuring the regional gene expression data (See *“Regional gene-expression profiles’”*).

**Fig.-7.**
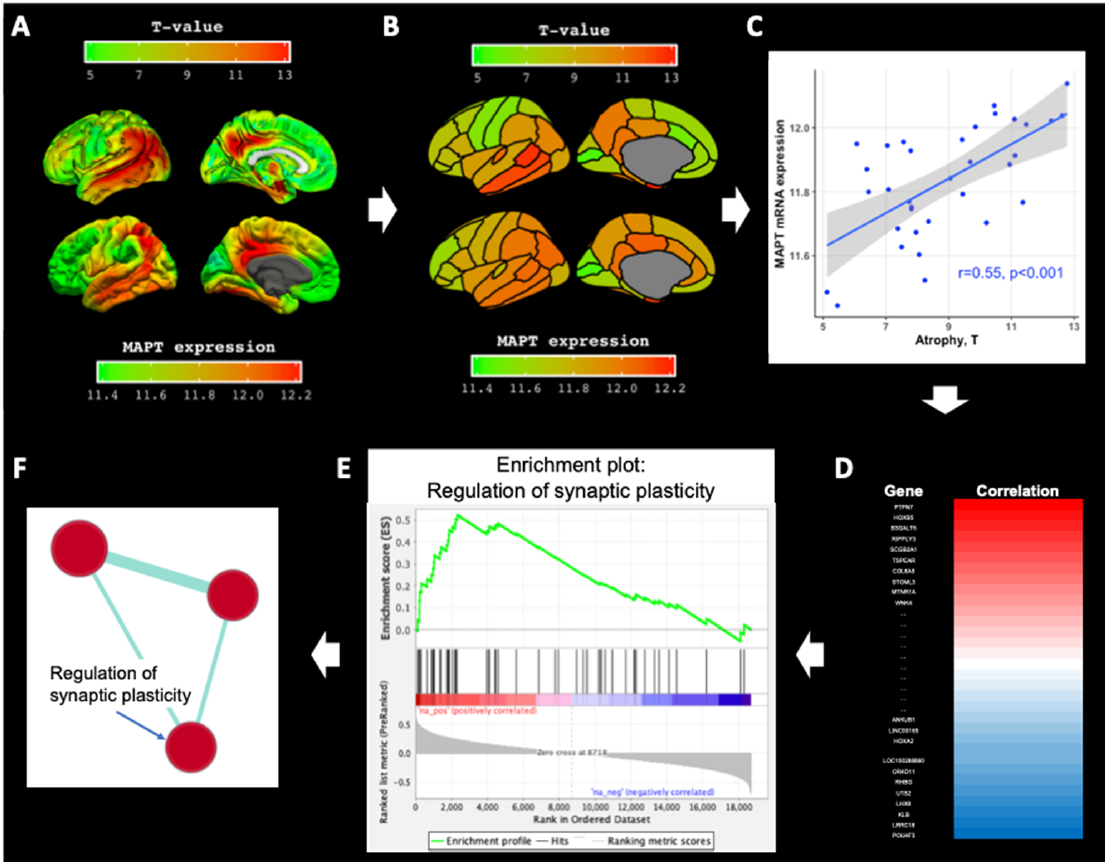
Gene-set enrichment analyses and grouping of gene sets. The figure displays how gene set enrichment analyses (GSEA) and grouping of gene sets were performed using atrophy in the whole sample and the MAPT expression profile as an example. Panel A displays voxel-wise differences in gray matter volumes between AD subjects and cognitively normal controls (top row; T-maps, adjusted for covariates), and brain-wide MAPT expression values obtained from the Allen database (bottom row). Panel B displays the parcellation of the T-map and gene expression profile into regions-of-interest (ROI) from the Desikan-Killiany atlas. T-values and gene expression values obtained within each ROI are then correlated across ROIs using Spearman’s correlation analyses (Panel C). The steps in panel A through C are repeated for all 18686 genes, yielding a correlation spectrum that is ranked from positive to negative (Panel D). GSEA software then uses *a priori* defined gene sets from curated databases and determines for each gene set where on the spectrum each gene within this set falls. This produces an enrichment score that indicates whether a gene set, as a whole, preferentially falls towards one end of the spectrum. Gene sets that cluster towards the positive end show higher than expected neurotypical expression levels in atrophic brain regions (‘positively enriched’), whereas those clustering towards the negative end show lower expression in atrophic brain regions (‘negatively enriched’). The example in Panel E shows positive enrichment for the gene set ‘synaptic plasticity’. Panel F displays results from the gene set grouping analysis in Cytoscape where significantly enriched gene sets are plotted as interrelated nodes connected by edges denoting their overlapping genes

To that end, we took the T-maps of the voxel-wise gray matter contrasts versus the amyloid-β negative cognitively normal control group (Fig.-7A) and extracted regional T-values using the Marsbar ROI toolbox implemented in SPM12 (Fig.-7B). For each gene, the 34 regional gene expression values were then related to the 34 regional T-values (atrophy) using Spearman’s correlations (Fig.-7C) (Krienen et al., 2016). This results in 18,686 correlation coefficients (one for each gene), indicating the relationship between brain-wide gene expression and the spatial pattern of atrophy. For the candidate genes analysis, we extracted the spatial (Spearman) correlation, with their associated p-values, between the candidate genes and atrophy in each of the subgroups. Then, for explorative analyses the correlation coefficients of all 18,686 genes were rank ordered according to the strength of the association (Fig.-7D). To reduce the wealth of gene-specific correlations into more comprehensible data we used gene set enrichment analysis (GSEA). GSEA is a statistical approach developed specifically to condense data from large microarrays on single genes into more comprehensible information on functional gene sets. GSEA uses the complete spectrum of information provided by the rank ordered gene-specific atrophy correlations and determines whether known gene sets, grouping genes with related biologic functions, are negatively or positively enriched for a specific atrophy pattern as a whole. In the present study, we explored 6,032 different gene sets obtained from the curated Reactome (http://reactome.org/; dataset: c2.cp.reactome.version:7.0 from the Molecular Signatures Database (MSigDB)), and gene ontology (GO; http://www.geneontology.org/; dataset: c5.all version 7.0) databases. These gene sets are defined by genes that are commonly associated with a specific biological process or GO annotation, respectively. Positive or negative enrichment of the gene sets was determined by the “Run GSEA pre-ranked” tool on default parameters and implemented in GSEA version 4.0.3. software package (available from: http://software.broadinstitute.org/gsea/index.jsp) by assessing whether genes within a set are non-normally distributed (skewed) towards one edge of the rank-ordered correlation spectrum (Fig.-7E). The non-normality of the distribution is represented by the normalized enrichment score, which also accounts for different sizes of the tested gene sets, and the statistical significance (p) of this score was adjusted using the false discovery rate (FDR). The threshold for statistical significance was set to PFDR<0.10 for comparisons within each of the subgroups (Subramanian et al., 2005). Using this approach, information regarding the spatial correlation between atrophy pattern and regional gene expression is summarized into gene sets. As a further data reduction step to improve interpretability of results, we used the Cytoscape add-in “Enrichment Map” (Merico et al., 2011) to group related gene sets into clusters of similar biological functions. Grouping is achieved by assessing similarity between gene sets (genes that are present in multiple gene sets) and results are visualized in a network structure where related gene sets (nodes) are connected by edges that represent the degree of overlapping genes, thus forming clusters (see Fig.-7F and figure legend). Additionally, we ran “Leading-edge” analyses in the GSEA 4.0.3. software package to identify the most relevant individual genes driving the enrichment signal for the different gene sets in a given cluster (“leading-edge genes”; see supplement).

## Acknowledgments

Research of Alzheimer center Amsterdam is part of the neurodegeneration research program of Amsterdam Neuroscience. Alzheimer Center Amsterdam is supported by Stichting Alzheimer Nederland and Stichting VUmc fonds. Wiesje van der Flier holds the Pasman chair. The clinical database structure was developed with funding from Stichting Dioraphte. The authors would further like to thank Murray Reed (funded by Austrian Academy of Sciences, DOC 928), Matej Murgaš (funded by FWF Austrian Science Fund DOC 33-B27) and Gregory Miles James.

