## Supplemental Material for "Differential Brain Atrophy Patterns and Neurogenetic Profiles in Cognitively-Defined Alzheimer’s Disease Subgroups"

### Supplementary Materials

| AD-Memory | Size | FDR(p) | ES | Cluster |
|---|---|---|---|---|
| **GO OXIDOREDUCTASE ACTIVITY ACTING ON NAD P H QUINONE OR SIMILAR COMPOUND AS ACCEPTOR** | 51 | 0.0379 | -0.4927 | Mitochondrial respiration |
| **REACTOME THE CITRIC ACID TCA CYCLE AND RESPIRATORY ELECTRON TRANSPORT** | 158 | 0.0029 | -0.4557 | Mitochondrial respiration |
| **REACTOME MITOCHONDRIAL CALCIUM ION TRANSPORT** | 23 | 0.0665 | -0.5656 | Mitochondrial respiration |
| **GO RESPIRATORY CHAIN COMPLEX** | 72 | 0.0043 | -0.5134 | Mitochondrial respiration |
| **REACTOME METABOLISM OF PORPHYRINS** | 15 | 0.0466 | -0.641 | Mitochondrial respiration |
| GO ORGANELLAR RIBOSOME | 87 | 0.0013 | -0.5283 | Mitochondrial respiration |
| **GO CELLULAR RESPIRATION** | 174 | 0.0227 | -0.4153 | Mitochondrial respiration |
| REACTOME RESPIRATORY ELECTRON TRANSPORT | 89 | 0.0025 | -0.5173 | Mitochondrial respiration |
| **GO AEROBIC RESPIRATION** | 80 | 0.0451 | -0.4467 | Mitochondrial respiration |
| GO MITOCHONDRIAL RESPIRATORY CHAIN COMPLEX I | 45 | 0.0027 | -0.5742 | Mitochondrial respiration |
| GO MITOCHONDRIAL ELECTRON TRANSPORT NADH TO UBIQUINONE | 45 | 0.0027 | -0.5746 | Mitochondrial respiration |
| **GO ENERGY COUPLED PROTON TRANSPORT DOWN ELECTROCHEMICAL GRADIENT** | 22 | 0.0876 | -0.5541 | Mitochondrial respiration |
| **GO NADH DEHYDROGENASE ACTIVITY** | 39 | 0.0052 | -0.5722 | Mitochondrial respiration |
| **REACTOME PROCESSING OF SMDT1** | 16 | 0.0578 | -0.632 | Mitochondrial respiration |
| GO NADH DEHYDROGENASE COMPLEX ASSEMBLY | 58 | 0.0025 | -0.5427 | Mitochondrial respiration |
| GO RESPIRATORY ELECTRON TRANSPORT CHAIN | 101 | 0.0045 | -0.4892 | Mitochondrial respiration |
| **GO RESPIRASOME** | 85 | 0.0021 | -0.513 | Mitochondrial respiration |

| | | | | |
|---|---|---|---|---|
| REACTOME RESPIRATORY ELECTRON TRANSPORT ATP SYNTHESIS BY CHEMIOSMOTIC COUPLING AND HEAT PRODUCTION BY UNCOUPLING PROTEINS | 109 | 0.002 | -0.5037 | Mitochondrial respiration |
| **REACTOME DEFECTS IN VITAMIN AND COFACTOR METABOLISM** | 21 | 0.0402 | -0.5966 | Mitochondrial respiration |
| REACTOME COMPLEX I BIOGENESIS | 49 | 0.003 | -0.5571 | Mitochondrial respiration |
| GO MITOCHONDRIAL MATRIX | 457 | 0.0029 | -0.4126 | Mitochondrial respiration |
| GO MITOCHONDRIAL PROTEIN COMPLEX | 250 | 0.0014 | -0.4709 | Mitochondrial respiration |
| **GO CYTOCHROME COMPLEX ASSEMBLY** | 33 | 0.0657 | -0.5101 | Mitochondrial respiration |
| GO ATP SYNTHESIS COUPLED ELECTRON TRANSPORT | 82 | 0.0019 | -0.5472 | Mitochondrial respiration |
| GO ORGANELLE ENVELOPE LUMEN | 85 | 0.0026 | -0.5055 | Mitochondrial respiration |
| GO NEGATIVE REGULATION OF CD4 POSITIVE ALPHA BETA T CELL DIFFERENTIATION | 18 | 0.0724 | -0.5952 | Mitochondrial respiration |
| **GO INNER MITOCHONDRIAL MEMBRANE PROTEIN COMPLEX** | 122 | 0.0109 | -0.449 | Mitochondrial respiration |
| GO ORGANELLE INNER MEMBRANE | 499 | 0.0045 | -0.4025 | Mitochondrial respiration |
| **GO MITOCHONDRIAL MEMBRANE PART** | 213 | 0.0461 | -0.3843 | Mitochondrial respiration |
| **GO ENERGY DERIVATION BY OXIDATION OF ORGANIC COMPOUNDS** | 260 | 0.0973 | -0.3512 | Mitochondrial respiration |
| GO OXIDATIVE PHOSPHORYLATION | 122 | 0.0019 | -0.5118 | Mitochondrial respiration |
| **REACTOME MITOCHONDRIAL PROTEIN IMPORT** | 65 | 0.0383 | -0.4711 | Mitochondrial respiration |
| **GO CYTOCHROME COMPLEX** | 25 | 0.0218 | -0.5998 | Mitochondrial respiration |
| **GO OXIDOREDUCTASE COMPLEX** | 99 | 0.0924 | -0.4019 | Mitochondrial respiration |
| **GO MITOCHONDRIAL RESPIRATORY CHAIN COMPLEX ASSEMBLY** | 89 | 0.0099 | -0.4736 | Mitochondrial respiration |
| **GO AEROBIC ELECTRON TRANSPORT CHAIN** | 17 | 0.0409 | -0.6337 | Mitochondrial respiration |
| **GO ELECTRON TRANSPORT CHAIN** | 166 | 0.0662 | -0.3877 | Mitochondrial respiration |

| | | | | |
|---|---|---|---|---|
| <u>**GO OXIDOREDUCTASE ACTIVITY ACTING ON A HEME GROUP OF DONORS**</u> | 24 | 0.0878 | -0.5554 | Mitochondrial respiration |
| GO MITOCHONDRIAL TRANSLATION | 135 | 0.0025 | -0.4851 | Mitochondrial respiration |
| GO MITOCHONDRIAL GENE EXPRESSION | 160 | 0.0017 | -0.483 | Mitochondrial respiration |
| <u>**GO RRNA BINDING**</u> | 62 | 0.0565 | -0.4597 | Metabolism of proteins |
| **GO STRUCTURAL CONSTITUENT OF RIBOSOME** | 154 | 0.0623 | -0.3956 | Metabolism of proteins |
| <u>**GO RIBOSOMAL LARGE SUBUNIT ASSEMBLY**</u> | 28 | 0.0435 | -0.5579 | Metabolism of proteins |
| <u>**GO RIBOSOME ASSEMBLY**</u> | 59 | 0.0106 | -0.5178 | Metabolism of proteins |
| **GO LARGE RIBOSOMAL SUBUNIT** | 116 | 0.0559 | -0.4094 | Metabolism of proteins |
| **GO RIBOSOMAL SUBUNIT** | 185 | 0.0375 | -0.4035 | Metabolism of proteins |
| <u>**REACTOME METABOLISM OF COFACTORS**</u> | 19 | 0.0282 | -0.6329 | Metabolism of proteins |
| <u>**GO RIBOSOME**</u> | 223 | 0.0471 | -0.3824 | Metabolism of proteins |
| <u>**GO SMALL RIBOSOMAL SUBUNIT**</u> | 71 | 0.0747 | -0.437 | Metabolism of proteins |
| **REACTOME TRANSLATION** | 290 | 0.0294 | -0.3872 | Metabolism of proteins |
| GO RNA METHYLTRANSFERASE ACTIVITY | 64 | 0.0373 | -0.4715 | Metabolism of proteins |
| GO MATURATION OF SSU RRNA | 44 | 0.0096 | -0.5373 | Metabolism of proteins |
| <u>**REACTOME TRNA PROCESSING**</u> | 107 | 0.0895 | -0.3955 | Metabolism of proteins |
| GO TRNA METHYLATION | 39 | 0.0228 | -0.5398 | Metabolism of proteins |
| REACTOME POSITIVE EPIGENETIC REGULATION OF RRNA EXPRESSION | 98 | 0.0452 | -0.4312 | Metabolism of proteins |
| GO CATALYTIC ACTIVITY ACTING ON A TRNA | 121 | 0.0279 | -0.4348 | Metabolism of proteins |
| GO INTERLEUKIN 7 MEDIATED SIGNALING PATHWAY | 27 | 0.0079 | -0.6112 | Metabolism of proteins |
| <u>**GO NCRNA PROCESSING**</u> | 369 | 0.0467 | -0.3628 | Metabolism of proteins |

| | | | | |
|---|---|---|---|---|
| **GO S ADENOSYLMETHIONINE DEPENDENT METHYLTRANSFERASE ACTIVITY** | 152 | 0.0961 | -0.3732 | Metabolism of proteins |
| GO TRANSCRIPTION BY RNA POLYMERASE I | 62 | 0.0563 | -0.4563 | Metabolism of proteins |
| REACTOME MRNA SPLICING MINOR PATHWAY | 52 | 0.0836 | -0.4629 | Metabolism of proteins |
| GO SPLICEOSOMAL COMPLEX | 175 | 0.0749 | -0.3827 | Metabolism of proteins |
| GO 3 5 RNA HELICASE ACTIVITY | 19 | 0.0454 | -0.5966 | Metabolism of proteins |
| GO AMINO ACID ACTIVATION | 49 | 0.0181 | -0.5224 | Metabolism of proteins |
| REACTOME RNA POLYMERASE I TRANSCRIPTION INITIATION | 46 | 0.0834 | -0.4731 | Metabolism of proteins |
| **REACTOME TRNA AMINOACYLATION** | 42 | 0.0226 | -0.5275 | Metabolism of proteins |
| **GO TRNA METHYLTRANSFERASE ACTIVITY** | 34 | 0.0378 | -0.539 | Metabolism of proteins |
| **GO TRNA PROCESSING** | 129 | 0.0296 | -0.4261 | Metabolism of proteins |
| **GO RRNA METABOLIC PROCESS** | 204 | 0.0455 | -0.3869 | Metabolism of proteins |
| GO PRECATALYTIC SPLICEOSOME | 44 | 0.0286 | -0.5161 | Metabolism of proteins |
| GO RNA METHYLATION | 79 | 0.038 | -0.4503 | Metabolism of proteins |
| REACTOME CYTOSOLIC TRNA AMINOACYLATION | 24 | 0.0274 | -0.601 | Metabolism of proteins |
| REACTOME ERCC6 CSB AND EHMT2 G9A POSITIVELY REGULATE RRNA EXPRESSION | 68 | 0.0657 | -0.4425 | Metabolism of proteins |
| REACTOME INTERLEUKIN 7 SIGNALING | 30 | 0.0046 | -0.6135 | Metabolism of proteins |
| GO CHROMATIN SILENCING AT RDNA | 38 | 0.0456 | -0.5097 | Metabolism of proteins |
| REACTOME RNA POLYMERASE I TRANSCRIPTION | 102 | 0.049 | -0.4279 | Metabolism of proteins |
| **GO RIBOSOME BIOGENESIS** | 278 | 0.0364 | -0.3857 | Metabolism of proteins |
| GO TRNA METABOLIC PROCESS | 177 | 0.0109 | -0.4309 | Metabolism of proteins |
| GO RESPONSE TO INTERLEUKIN 7 | 36 | 0.005 | -0.5854 | Metabolism of proteins |

| | | | | |
|---|---|---|---|---|
| **GO CATALYTIC ACTIVITY ACTING ON RNA** | 335 | 0.0234 | -0.3878 | Metabolism of proteins |
| GO MATURATION OF SSU RRNA FROM TRICISTRONIC RRNA TRANSCRIPT SSU RRNA 5 8S RRNA LSU RRNA | 34 | 0.0461 | -0.5271 | Metabolism of proteins |
| GO PRERIBOSOME | 71 | 0.0223 | -0.4726 | Metabolism of proteins |
| **GO NCRNA METABOLIC PROCESS** | 445 | 0.0775 | -0.3433 | Metabolism of proteins |
| **GO PRERIBOSOME LARGE SUBUNIT PRECURSOR** | 20 | 0.036 | -0.621 | Metabolism of proteins |
| **GO RIBOSOMAL SMALL SUBUNIT BIOGENESIS** | 63 | 0.0302 | -0.4739 | Metabolism of proteins |
| **GO TRANSFERASE ACTIVITY TRANSFERRING ONE CARBON GROUPS** | 218 | 0.0487 | -0.3825 | Metabolism of proteins |
| **GO LIGASE ACTIVITY FORMING CARBON OXYGEN BONDS** | 41 | 0.0586 | -0.4992 | Metabolism of proteins |
| GO PRECATALYTIC SPLICEOSOME | 44 | 0.0286 | -0.5161 | Metabolism of RNA splicing |
| GO 3 5 RNA HELICASE ACTIVITY | 19 | 0.0454 | -0.5966 | Metabolism of RNA splicing |
| REACTOME MRNA SPLICING MINOR PATHWAY | 52 | 0.0836 | -0.4629 | Metabolism of RNA splicing |
| GO SPLICEOSOMAL COMPLEX | 175 | 0.0749 | -0.3827 | Metabolism of RNA splicing |
| REACTOME INTERLEUKIN 7 SIGNALING | 30 | 0.0046 | -0.6135 | Immune system |
| GO TRANSCRIPTION BY RNA POLYMERASE I | 62 | 0.0563 | -0.4563 | Immune system |
| REACTOME RNA POLYMERASE I TRANSCRIPTION | 102 | 0.049 | -0.4279 | Immune system |
| REACTOME ERCC6 CSB AND EHMT2 G9A POSITIVELY REGULATE RRNA EXPRESSION | 68 | 0.0657 | -0.4425 | Immune system |
| GO RESPONSE TO INTERLEUKIN 7 | 36 | 0.005 | -0.5854 | Immune system |
| REACTOME POSITIVE EPIGENETIC REGULATION OF RRNA EXPRESSION | 98 | 0.0452 | -0.4312 | Immune system |
| GO CHROMATIN SILENCING AT RDNA | 38 | 0.0456 | -0.5097 | Immune system |
| GO INTERLEUKIN 7 MEDIATED SIGNALING PATHWAY | 27 | 0.0079 | -0.6112 | Immune system |

| | | | | |
|---|---|---|---|---|
| REACTOME RNA POLYMERASE I TRANSCRIPTION INITIATION | 46 | 0.0834 | -0.4731 | Immune system |
| **REACTOME DNA REPLICATION PRE INITIATION** | 79 | 0.0875 | -0.4259 | Cell cycle |
| **REACTOME STABILIZATION OF P53** | 53 | 0.0399 | -0.4858 | Cell cycle |
| **REACTOME SWITCHING OF ORIGINS TO A POST REPLICATIVE STATE** | 86 | 0.0813 | -0.4242 | Cell cycle |
| **REACTOME VIF MEDIATED DEGRADATION OF APOBEC3G** | 51 | 0.0556 | -0.4736 | Cell cycle |
| **REACTOME CROSS PRESENTATION OF SOLUBLE EXOGENOUS ANTIGENS ENDOSOMES** | 47 | 0.0657 | -0.4708 | Cell cycle |
| **REACTOME ASSEMBLY OF THE PRE REPLICATIVE COMPLEX** | 64 | 0.0493 | -0.4617 | Cell cycle |
| **REACTOME S PHASE** | 155 | 0.0878 | -0.3819 | Cell cycle |
| **REACTOME DEGRADATION OF AXIN** | 52 | 0.0393 | -0.487 | Cell cycle |
| **REACTOME NEGATIVE REGULATION OF NOTCH4 SIGNALING** | 52 | 0.0662 | -0.4681 | Cell cycle |
| **REACTOME DEGRADATION OF DVL** | 54 | 0.0747 | -0.4622 | Cell cycle |
| **REACTOME G1 S DNA DAMAGE CHECKPOINTS** | 64 | 0.0563 | -0.4506 | Cell cycle |
| **REACTOME REGULATION OF RUNX2 EXPRESSION AND ACTIVITY** | 69 | 0.0455 | -0.4466 | Cell cycle |
| **REACTOME CDK MEDIATED PHOSPHORYLATION AND REMOVAL OF CDC6** | 69 | 0.0724 | -0.442 | Cell cycle |
| **REACTOME PCP CE PATHWAY** | 88 | 0.0836 | -0.4164 | Cell cycle |
| **GO THREONINE TYPE PEPTIDASE ACTIVITY** | 20 | 0.0906 | -0.5664 | Cell cycle |
| **REACTOME DNA REPLICATION** | 121 | 0.0855 | -0.3956 | Cell cycle |
| **REACTOME REGULATION OF APOPTOSIS** | 50 | 0.0944 | -0.4543 | Cell cycle |
| **REACTOME AUF1 HNRNP D0 BINDS AND DESTABILIZES MRNA** | 52 | 0.0653 | -0.4643 | Cell cycle |
| **REACTOME REGULATION OF RUNX3 EXPRESSION AND ACTIVITY** | 53 | 0.0968 | -0.4442 | Cell cycle |

| | | | | |
|---|---|---|---|---|
| **GO PROTEASOME CORE COMPLEX** | 19 | 0.0666 | -0.5999 | Cell cycle |
| **REACTOME FBXL7 DOWN REGULATES AURKA DURING MITOTIC ENTRY AND IN EARLY MITOSIS** | 52 | 0.0863 | -0.4489 | Cell cycle |
| GO MITOPHAGY | 21 | 0.0235 | -0.6136 | Autophagy |
| GO RESPONSE TO MITOCHONDRIAL DEPOLARISATION | 20 | 0.0231 | -0.6392 | Autophagy |
| **GO REGULATION OF MITOPHAGY** | 16 | 0.0837 | -0.6123 | Autophagy |
| **GO MHC PROTEIN COMPLEX** | 24 | 0.0891 | -0.5459 | Membrane proteins |
| **GO LUMENAL SIDE OF MEMBRANE** | 32 | 0.045 | -0.5288 | Membrane proteins |
| **GO CELL REDOX HOMEOSTASIS** | 73 | 0.0977 | -0.4196 | - |
| **GO AMINOPEPTIDASE ACTIVITY** | 45 | 0.0379 | -0.5039 | - |
| **GO PERINUCLEAR ENDOPLASMIC RETICULUM** | 22 | 0.0964 | -0.5492 | - |
| **GO PROTEIN ADP RIBOSYLATION** | 29 | 0.0967 | -0.5094 | - |
| **REACTOME PURINE CATABOLISM** | 16 | 0.0288 | -0.6622 | - |
| **GO PINOCYTOSIS** | 21 | 0.0774 | -0.5703 | - |
| GO MANNOSYLTRANSFERASE ACTIVITY | 26 | 0.0229 | -0.5872 | - |
| **REACTOME NEUREXINS AND NEUROLIGINS** | 57 | 0.0297 | 0.5303 | Synapse function |
| **REACTOME DOPAMINE NEUROTRANSMITTER RELEASE CYCLE** | 23 | 0.0369 | 0.6104 | Synapse function |
| GO VOCALIZATION BEHAVIOR | 17 | 0.0336 | 0.68 | Synapse function |
| **REACTOME PROTEIN PROTEIN INTERACTIONS AT SYNAPSES** | 88 | 0.019 | 0.4752 | Synapse function |
| **GO REGULATION OF LONG TERM NEURONAL SYNAPTIC PLASTICITY** | 26 | 0.02 | 0.6364 | Synaptic plasticity |
| GO REGULATION OF NEURONAL SYNAPTIC PLASTICITY | 50 | 0.0239 | 0.5476 | Synaptic plasticity |

| | | | | |
|---|---|---|---|---|
| **GO NEGATIVE REGULATION OF SYNAPSE ORGANIZATION** | 22 | 0.0988 | 0.5916 | - |
| **GO ENTRY OF BACTERIUM INTO HOST CELL** | 15 | 0.0667 | 0.6801 | - |
| **GO RENAL VESICLE DEVELOPMENT** | 18 | 0.0928 | 0.6129 | - |
| **GO OXIDOREDUCTASE ACTIVITY ACTING ON PAIRED DONORS WITH INCORPORATION OR REDUCTION OF MOLECULAR OXYGEN REDUCED FLAVIN OR FLAVOPROTEIN AS ONE DONOR AND INCORPORATION OF ONE ATOM OF OXYGEN** | 29 | 0.0553 | 0.5538 | - |
| **AD-Executive Functioning** | **Size** | **FDR(p)** | **ES** | **Cluster** |

| | | | | |
|---|---|---|---|---|
| <u>GO NUCLEOID</u> | 43 | 0.0849 | -0.4965 | Metabolism of proteins |
| GO MITOCHONDRIAL TRANSLATION | 135 | 0.0637 | -0.4199 | Metabolism of proteins |
| GO MITOCHONDRIAL GENE EXPRESSION | 160 | 0.0242 | -0.442 | Metabolism of proteins |
| GO TRANSLATIONAL TERMINATION | 104 | 0.0722 | -0.4335 | Metabolism of proteins |
| REACTOME CYTOSOLIC TRNA AMINOACYLATION | 24 | 0.0728 | -0.5716 | Metabolism of proteins |
| GO ORGANELLAR LARGE RIBOSOMAL SUBUNIT | 57 | 0.0785 | -0.4755 | Metabolism of proteins |
| GO ORGANELLAR RIBOSOME | 87 | 0.0818 | -0.4435 | Metabolism of proteins |
| GO MITOCHONDRIAL TRANSLATIONAL TERMINATION | 89 | 0.042 | -0.4614 | Metabolism of proteins |
| **<u>GO MITOCHONDRIAL TRANSCRIPTION</u>** | 15 | 0.0907 | -0.6455 | Metabolism of proteins |
| REACTOME MITOCHONDRIAL TRANSLATION | 94 | 0.0297 | -0.4619 | Metabolism of proteins |
| GO AMINO ACID ACTIVATION | 49 | 0.075 | -0.4885 | Metabolism of proteins |
| **GO POSITIVE REGULATION OF MESENCHYMAL CELL PROLIFERATION** | 25 | 0.0932 | -0.5609 | - |
| **GO BITTER TASTE RECEPTOR ACTIVITY** | 19 | 0.0854 | -0.6462 | - |
| **<u>GO POLY PURINE TRACT BINDING</u>** | 30 | 0.0849 | -0.5406 | - |
| **GO CATECHOLAMINE UPTAKE** | 15 | 0.073 | 0.6516 | Synapse function / Synaptic plasticity |
| **GO POSTSYNAPTIC SPECIALIZATION ORGANIZATION** | 34 | 0.0655 | 0.5402 | Synapse function / Synaptic plasticity |
| **GO NEURON TO NEURON SYNAPSE** | 338 | 0.0645 | 0.3535 | Synapse function / Synaptic plasticity |

| | | | | |
|---|---|---|---|---|
| **REACTOME NEUREXINS AND NEUROLIGINS** | 57 | 0.0625 | 0.4774 | Synapse function / Synaptic plasticity |
| **GO IONOTROPIC GLUTAMATE RECEPTOR BINDING** | 32 | 0.068 | 0.5401 | Synapse function / Synaptic plasticity |
| GO REGULATION OF NEURONAL SYNAPTIC PLASTICITY | 50 | 0.0269 | 0.5147 | Synapse function / Synaptic plasticity |
| GO POSITIVE REGULATION OF NEUROTRANSMITTER TRANSPORT | 38 | 0.0784 | 0.49 | Synapse function / Synaptic plasticity |
| **GO NEGATIVE REGULATION OF SYNAPSE ORGANIZATION** | 22 | 0.0336 | 0.6412 | Synapse function / Synaptic plasticity |
| GO VOCALIZATION BEHAVIOR | 17 | 0.0301 | 0.7064 | Synapse function / Synaptic plasticity |
| **REACTOME PROTEIN PROTEIN INTERACTIONS AT SYNAPSES** | 88 | 0.0924 | 0.4114 | Synapse function / Synaptic plasticity |
| **GO INTRASPECIES INTERACTION BETWEEN ORGANISMS** | 53 | 0.087 | 0.4541 | Synapse function / Synaptic plasticity |
| **GO POSTSYNAPTIC SPECIALIZATION ASSEMBLY** | 22 | 0.0902 | 0.5628 | Synapse function / Synaptic plasticity |
| **GO REGULATION OF LONG TERM NEURONAL SYNAPTIC PLASTICITY** | 26 | 0.0681 | 0.5525 | Synapse function / Synaptic plasticity |
| **REACTOME REGULATION OF ACTIN DYNAMICS FOR PHAGOCYTIC CUP FORMATION** | 60 | 0.0287 | 0.4891 | Immune system |
| **GO FC RECEPTOR MEDIATED STIMULATORY SIGNALING PATHWAY** | 82 | 0.0804 | 0.419 | Immune system |
| **GO ARP2 3 COMPLEX MEDIATED ACTIN NUCLEATION** | 33 | 0.0782 | 0.5165 | Immune system |
| **REACTOME RHO GTPASES ACTIVATE WASPS AND WAVES** | 33 | 0.0268 | 0.5702 | Immune system |

| | | | | |
|---|---|---|---|---|
| **REACTOME NONSENSE MEDIATED DECAY NMD** | 113 | 0.0696 | 0.4069 | Metabolism of RNA |
| **REACTOME NONSENSE MEDIATED DECAY NMD INDEPENDENT OF THE EXON JUNCTION COMPLEX EJC** | 93 | 0.0351 | 0.4565 | Metabolism of RNA |
| **GO CYTOSOLIC LARGE RIBOSOMAL SUBUNIT** | 58 | 0.0807 | 0.4621 | Metabolism of RNA |
| **GO REGULATION OF EXCRETION** | 22 | 0.0799 | 0.5851 | - |
| **GO APOLIPOPROTEIN BINDING** | 17 | 0.0686 | 0.6219 | - |
| **GO OXIDOREDUCTASE ACTIVITY ACTING ON PAIRED DONORS WITH INCORPORATION OR REDUCTION OF MOLECULAR OXYGEN REDUCED FLAVIN OR FLAVOPROTEIN AS ONE DONOR AND INCORPORATION OF ONE ATOM OF OXYGEN** | 29 | 0.0721 | 0.5293 | - |
| **AD-Language** | **Size** | **FDR(p)** | **ES** | **Cluster** |

| | | | | |
|---|---|---|---|---|
| **GO TASTE RECEPTOR ACTIVITY** | 23 | 0.0711 | -0.6213 | Taste receptor |
| **GO BITTER TASTE RECEPTOR ACTIVITY** | 19 | 0.0126 | -0.7051 | Taste receptor |
| **GO CYTOSOLIC RIBOSOME** | 107 | 0.0002 | 0.5031 | Metabolism of proteins |
| **REACTOME ACTIVATION OF THE MRNA UPON BINDING OF THE CAP BINDING COMPLEX AND EIFS AND SUBSEQUENT BINDING TO 43S** | 59 | 0.0014 | 0.5279 | Metabolism of proteins |
| **GO ESTABLISHMENT OF PROTEIN LOCALIZATION TO ENDOPLASMIC RETICULUM** | 110 | 0.0014 | 0.4641 | Metabolism of proteins |
| **REACTOME INFLUENZA INFECTION** | 151 | 0.0013 | 0.4358 | Metabolism of proteins |
| **GO COTRANSLATIONAL PROTEIN TARGETING TO MEMBRANE** | 97 | 0.0005 | 0.5034 | Metabolism of proteins |
| **REACTOME REGULATION OF EXPRESSION OF SLITS AND ROBOS** | 165 | 0.0006 | 0.4562 | Metabolism of proteins |
| **REACTOME NONSENSE MEDIATED DECAY NMD INDEPENDENT OF THE EXON JUNCTION COMPLEX EJC** | 93 | 0 | 0.5732 | Metabolism of proteins |
| **GO CYTOSOLIC LARGE RIBOSOMAL SUBUNIT** | 58 | 0.0009 | 0.5442 | Metabolism of proteins |
| **REACTOME NONSENSE MEDIATED DECAY NMD** | 113 | 0 | 0.5289 | Metabolism of proteins |
| **REACTOME SRP DEPENDENT COTRANSLATIONAL PROTEIN TARGETING TO MEMBRANE** | 110 | 0 | 0.5051 | Metabolism of proteins |
| **GO NUCLEAR TRANSCRIBED MRNA CATABOLIC PROCESS NONSENSE MEDIATED DECAY** | 118 | 0.0011 | 0.4708 | Metabolism of proteins |
| **REACTOME SIGNALING BY ROBO RECEPTORS** | 210 | 0.0064 | 0.3938 | Metabolism of proteins |
| **REACTOME EUKARYOTIC TRANSLATION INITIATION** | 117 | 0 | 0.5199 | Metabolism of proteins |
| **GO CYTOSOLIC PART** | 237 | 0.0985 | 0.3368 | Metabolism of proteins |
| **GO PROTEIN TARGETING TO MEMBRANE** | 188 | 0.0062 | 0.4048 | Metabolism of proteins |
| **GO PROTEIN LOCALIZATION TO ENDOPLASMIC RETICULUM** | 134 | 0.0228 | 0.4058 | Metabolism of proteins |

| | | | | |
|---|---|---|---|---|
| **REACTOME SELENOAMINO ACID METABOLISM** | 114 | 0 | 0.5098 | Metabolism of proteins |
| **GO NEURON CELL CELL ADHESION** | 16 | 0.0592 | 0.6181 | Synapse function / Synaptic plasticity |
| GO VOCALIZATION BEHAVIOR | 17 | 0.0233 | 0.6362 | Synapse function / Synaptic plasticity |
| **GO POSTSYNAPTIC SPECIALIZATION ORGANIZATION** | 34 | 0.0287 | 0.5368 | Synapse function / Synaptic plasticity |
| **GO NEGATIVE REGULATION OF SYNAPSE ORGANIZATION** | 22 | 0.0013 | 0.6517 | Synapse function / Synaptic plasticity |
| **GO IONOTROPIC GLUTAMATE RECEPTOR BINDING** | 32 | 0.0444 | 0.5269 | Synapse function / Synaptic plasticity |
| GO REGULATION OF NEURONAL SYNAPTIC PLASTICITY | 50 | 0.0714 | 0.4652 | Synapse function / Synaptic plasticity |
| **GO POSTSYNAPTIC SPECIALIZATION ASSEMBLY** | 22 | 0.0591 | 0.5692 | Synapse function / Synaptic plasticity |
| **GO NEURON TO NEURON SYNAPSE** | 338 | 0.0654 | 0.3307 | Synapse function / Synaptic plasticity |
| **GO ARP2 3 COMPLEX MEDIATED ACTIN NUCLEATION** | 33 | 0.0455 | 0.519 | Immune system |
| **REACTOME REGULATION OF ACTIN DYNAMICS FOR PHAGOCYTIC CUP FORMATION** | 60 | 0.0231 | 0.468 | Immune system |
| **GO FC RECEPTOR MEDIATED STIMULATORY SIGNALING PATHWAY** | 82 | 0.0882 | 0.4062 | Immune system |
| **REACTOME RHO GTPASES ACTIVATE WASPS AND WAVES** | 33 | 0.0214 | 0.5379 | Immune system |

| | | | | |
|---|---|---|---|---|
| **GO CATECHOLAMINE UPTAKE** | 15 | 0.0743 | 0.6269 | - |
| **GO APOLIPOPROTEIN BINDING** | 17 | 0.0608 | 0.6214 | - |
| **GO REGULATION OF EXCRETION** | 22 | 0.0277 | 0.5838 | - |
| **REACTOME INSERTION OF TAIL ANCHORED PROTEINS INTO THE ENDOPLASMIC RETICULUM MEMBRANE** | 22 | 0.0727 | 0.5632 | - |
| **AD-Visuospatial Functioning** | **Size** | **FDR(p)** | **ES** | **Cluster** |
| **REACTOME POSITIVE EPIGENETIC REGULATION OF RRNA EXPRESSION** | 98 | 0.0387 | -0.4523 | Gene expression |

| | | | | |
|---|---|---|---|---|
| **REACTOME DEPURINATION** | 50 | 0.041 | -0.5027 | Gene expression |
| **REACTOME PRE NOTCH EXPRESSION AND PROCESSING** | 100 | 0.0696 | -0.4188 | Gene expression |
| **REACTOME HDACS DEACETYLATE HISTONES** | 87 | 0.0101 | -0.4943 | Gene expression |
| GO RESPONSE TO INTERLEUKIN 7 | 36 | 0.0414 | -0.5299 | Gene expression |
| **REACTOME FORMATION OF THE BETA CATENIN:TCF TRANSACTIVATING COMPLEX** | 84 | 0.0669 | -0.4291 | Gene expression |
| **REACTOME SIRT1 NEGATIVELY REGULATES RRNA EXPRESSION** | 60 | 0.0157 | -0.5108 | Gene expression |
| **REACTOME DNA METHYLATION** | 56 | 0.0103 | -0.5376 | Gene expression |
| **REACTOME NUCLEOSOME ASSEMBLY** | 64 | 0.0869 | -0.4442 | Gene expression |
| **REACTOME NEGATIVE EPIGENETIC REGULATION OF RRNA EXPRESSION** | 100 | 0.0404 | -0.4424 | Gene expression |
| **REACTOME SENESCENCE ASSOCIATED SECRETORY PHENOTYPE SASP** | 101 | 0.055 | -0.4224 | Gene expression |
| **REACTOME ACTIVATED PKN1 STIMULATES TRANSCRIPTION OF AR ANDROGEN RECEPTOR REGULATED GENES KLK2 AND KLK3** | 59 | 0.0173 | -0.501 | Gene expression |
| **REACTOME PRC2 METHYLATES HISTONES AND DNA** | 65 | 0.0149 | -0.5004 | Gene expression |
| **REACTOME ERCC6 CSB AND EHMT2 G9A POSITIVELY REGULATE RRNA EXPRESSION** | 68 | 0.0025 | -0.535 | Gene expression |
| **REACTOME MEIOTIC RECOMBINATION** | 76 | 0.0429 | -0.4578 | Gene expression |
| **GO INTERLEUKIN 7 MEDIATED SIGNALING PATHWAY** | 27 | 0.0995 | -0.5297 | Gene expression |
| **REACTOME RNA POLYMERASE I TRANSCRIPTION** | 102 | 0.0136 | -0.4632 | Gene expression |
| **REACTOME BASE EXCISION REPAIR AP SITE FORMATION** | 57 | 0.0618 | -0.4618 | Gene expression |
| **REACTOME EPIGENETIC REGULATION OF GENE EXPRESSION** | 138 | 0.0433 | -0.4116 | Gene expression |
| **REACTOME RNA POLYMERASE I PROMOTER ESCAPE** | 82 | 0.0417 | -0.4552 | Gene expression |

| | | | | |
|---|---|---|---|---|
| **<u>REACTOME RUNX1 REGULATES TRANSCRIPTION OF GENES INVOLVED IN DIFFERENTIATION OF HSCS</u>** | 119 | 0.0475 | -0.4205 | Gene expression |
| **<u>REACTOME TRANSCRIPTIONAL REGULATION OF GRANULOPOIESIS</u>** | 81 | 0.0544 | -0.4423 | Gene expression |
| **<u>REACTOME CONDENSATION OF PROPHASE CHROMOSOMES</u>** | 66 | 0.0731 | -0.4429 | Gene expression |
| GO RESPIRATORY ELECTRON TRANSPORT CHAIN | 101 | 0.1 | -0.4031 | Mitochondrial respiration |
| **GO NADH DEHYDROGENASE ACTIVITY** | 39 | 0.0475 | -0.5198 | Mitochondrial respiration |
| GO MITOCHONDRIAL ELECTRON TRANSPORT NADH TO UBIQUINONE | 45 | 0.0153 | -0.5374 | Mitochondrial respiration |
| GO OXIDATIVE PHOSPHORYLATION | 122 | 0.043 | -0.428 | Mitochondrial respiration |
| REACTOME RESPIRATORY ELECTRON TRANSPORT ATP SYNTHESIS BY CHEMIOSMOTIC COUPLING AND HEAT PRODUCTION BY UNCOUPLING PROTEINS | 109 | 0.0929 | -0.3961 | Mitochondrial respiration |
| REACTOME COMPLEX I BIOGENESIS | 49 | 0.051 | -0.4851 | Mitochondrial respiration |
| REACTOME RESPIRATORY ELECTRON TRANSPORT | 89 | 0.0926 | -0.4175 | Mitochondrial respiration |
| GO ATP SYNTHESIS COUPLED ELECTRON TRANSPORT | 82 | 0.044 | -0.4545 | Mitochondrial respiration |
| GO MITOCHONDRIAL RESPIRATORY CHAIN COMPLEX I | 45 | 0.0428 | -0.5087 | Mitochondrial respiration |
| **GO RIBOSOMAL SUBUNIT** | 185 | 0.0543 | -0.3887 | Metabolism of proteins |
| **<u>GO FATTY ACID CATABOLIC PROCESS</u>** | 97 | 0.052 | -0.4268 | Metabolism of proteins |
| GO PRERIBOSOME | 71 | 0.0603 | -0.4505 | Metabolism of proteins |
| **GO STRUCTURAL CONSTITUENT OF RIBOSOME** | 154 | 0.0957 | -0.384 | Metabolism of proteins |
| GO CATALYTIC ACTIVITY ACTING ON A TRNA | 121 | 0.056 | -0.4098 | Metabolism of proteins |
| GO MATURATION OF SSU RRNA FROM TRICISTRONIC RRNA TRANSCRIPT SSU RRNA 5 8S RRNA LSU RRNA | 34 | 0.0934 | -0.5066 | Metabolism of proteins |

| | | | | |
|---|---|---|---|---|
| **GO PRERIBOSOME LARGE SUBUNIT PRECURSOR** | 20 | 0.0985 | -0.5724 | Metabolism of proteins |
| REACTOME MITOCHONDRIAL TRANSLATION | 94 | 0 | -0.5373 | Metabolism of proteins |
| REACTOME MITOCHONDRIAL FATTY ACID BETA OXIDATION | 35 | 0.0149 | -0.5741 | Metabolism of proteins |
| GO MITOCHONDRIAL MATRIX | 457 | 0.0093 | -0.3947 | Metabolism of proteins |
| GO ORGANELLE INNER MEMBRANE | 499 | 0.0937 | -0.3319 | Metabolism of proteins |
| **GO RIBOSOMAL SMALL SUBUNIT BIOGENESIS** | 63 | 0.0727 | -0.4491 | Metabolism of proteins |
| **GO ORGANELLAR SMALL RIBOSOMAL SUBUNIT** | 28 | 0.0424 | -0.5603 | Metabolism of proteins |
| **REACTOME TRANSLATION** | 290 | 0.0622 | -0.3634 | Metabolism of proteins |
| GO AMINO ACID ACTIVATION | 49 | 0.0727 | -0.473 | Metabolism of proteins |
| **GO LARGE RIBOSOMAL SUBUNIT** | 116 | 0.0933 | -0.394 | Metabolism of proteins |
| GO MATURATION OF SSU RRNA | 44 | 0.0862 | -0.4796 | Metabolism of proteins |
| GO TRANSLATIONAL TERMINATION | 104 | 0.0094 | -0.4836 | Metabolism of proteins |
| GO TRNA METHYLATION | 39 | 0.0933 | -0.4876 | Metabolism of proteins |
| GO MITOCHONDRIAL PROTEIN COMPLEX | 250 | 0.0113 | -0.4194 | Metabolism of proteins |
| GO MITOCHONDRIAL TRANSLATION | 135 | 0.011 | -0.4587 | Metabolism of proteins |
| GO TRANSLATIONAL ELONGATION | 132 | 0.0359 | -0.4303 | Metabolism of proteins |
| GO ORGANELLAR LARGE RIBOSOMAL SUBUNIT | 57 | 0.0008 | -0.5635 | Metabolism of proteins |
| GO MITOCHONDRIAL TRANSLATIONAL TERMINATION | 89 | 0 | -0.5329 | Metabolism of proteins |
| GO TRNA METHYLTRANSFERASE ACTIVITY | 34 | 0.099 | -0.49 | Metabolism of proteins |
| **GO RIBOSOME BIOGENESIS** | 278 | 0.0995 | -0.3507 | Metabolism of proteins |
| GO MITOCHONDRIAL GENE EXPRESSION | 160 | 0.0093 | -0.4432 | Metabolism of proteins |

| | | | | |
|---|---|---|---|---|
| GO ORGANELLAR RIBOSOME | 87 | 0 | -0.5307 | Metabolism of proteins |
| **REACTOME DEFECTS IN VITAMIN AND COFACTOR METABOLISM** | 21 | 0.0939 | -0.5719 | Metabolism of proteins |
| **GO MONOCARBOXYLIC ACID CATABOLIC PROCESS** | 113 | 0.0731 | -0.4082 | Metabolism of proteins |
| GO RNA METHYLTRANSFERASE ACTIVITY | 64 | 0.0813 | -0.4483 | Metabolism of proteins |
| **GO LIGASE ACTIVITY FORMING CARBON OXYGEN BONDS** | 41 | 0.0855 | -0.4836 | Metabolism of proteins |
| GO CATALYTIC ACTIVITY ACTING ON A TRNA | 121 | 0.056 | -0.4098 | Metabolism of proteins |
| GO RNA METHYLTRANSFERASE ACTIVITY | 64 | 0.0813 | -0.4483 | Metabolism of proteins |
| GO TRNA METHYLTRANSFERASE ACTIVITY | 34 | 0.099 | -0.49 | Metabolism of proteins |
| GO TRNA METHYLATION | 39 | 0.0933 | -0.4876 | Metabolism of proteins |
| **REACTOME GLUCONEOGENESIS** | 31 | 0.0973 | -0.5056 | Metabolism carbohydrates |
| **REACTOME METABOLISM OF CARBOHYDRATES** | 280 | 0.0864 | -0.3558 | Metabolism carbohydrates |
| **GO LYSOSOMAL LUMEN** | 85 | 0.0601 | -0.4343 | Metabolism carbohydrates |
| **GO NEGATIVE REGULATION OF ALPHA BETA T CELL ACTIVATION** | 33 | 0.0371 | -0.5588 | Immune system |
| **GO NEGATIVE REGULATION OF ALPHA BETA T CELL DIFFERENTIATION** | 21 | 0.0387 | -0.6215 | Immune system |
| **GO NEGATIVE REGULATION OF HEMOPOIESIS** | 130 | 0.0941 | -0.3924 | Immune system |
| **GO NEGATIVE REGULATION OF T CELL DIFFERENTIATION** | 33 | 0.0442 | -0.5491 | Immune system |
| **GO TRANSFERASE ACTIVITY TRANSFERRING ALKYL OR ARYL OTHER THAN METHYL GROUPS** | 55 | 0.0827 | -0.4588 | ? |
| **GO GLUTATHIONE DERIVATIVE BIOSYNTHETIC PROCESS** | 21 | 0.0882 | -0.5747 | ? |

| | | | | |
|---|---|---|---|---|
| **GO LUMENAL SIDE OF MEMBRANE** | 32 | 0.0945 | -0.515 | - |
| **GO POSITIVE REGULATION OF CALCIUM ION IMPORT** | 16 | 0.0873 | -0.6191 | - |
| **GO POSITIVE REGULATION OF MESENCHYMAL CELL PROLIFERATION** | 25 | 0.0561 | -0.5561 | - |
| **REACTOME MRNA SPLICING MINOR PATHWAY** | 52 | 0.0993 | -0.4555 | - |
| **REACTOME ASSOCIATION OF TRIC CCT WITH TARGET PROTEINS DURING BIOSYNTHESIS** | 39 | 0.0958 | -0.4921 | - |
| **REACTOME BIOSYNTHESIS OF THE N GLYCAN PRECURSOR DOLICHOL LIPID LINKED OLIGOSACCHARIDE LLO AND TRANSFER TO A NASCENT PROTEIN** | 78 | 0.0889 | -0.4279 | - |
| **GO REGULATION OF LONG TERM NEURONAL SYNAPTIC PLASTICITY** | 26 | 0.0193 | 0.6231 | Synapse function / Synaptic plasticity |
| **GO VISUAL BEHAVIOR** | 51 | 0.0783 | 0.4882 | Synapse function / Synaptic plasticity |
| **GO NEURON TO NEURON SYNAPSE** | 338 | 0.0976 | 0.3535 | Synapse function / Synaptic plasticity |
| **REACTOME NEUREXINS AND NEUROLIGINS** | 57 | 0.0939 | 0.4725 | Synapse function / Synaptic plasticity |
| GO REGULATION OF NEURONAL SYNAPTIC PLASTICITY | 50 | 0.0157 | 0.5453 | Synapse function / Synaptic plasticity |
| **GO REGULATION OF DENDRITIC SPINE MORPHOGENESIS** | 39 | 0.0364 | 0.5544 | Synapse function / Synaptic plasticity |
| GO POSITIVE REGULATION OF NEUROTRANSMITTER TRANSPORT | 38 | 0.0969 | 0.519 | Synapse function / Synaptic plasticity |
| GO VOCALIZATION BEHAVIOR | 17 | 0.0113 | 0.73 | Synapse function / Synaptic plasticity |

| | | | | |
|---|---|---|---|---|
| **GO INTRASPECIES INTERACTION BETWEEN ORGANISMS** | 53 | 0.0838 | 0.4831 | Synapse function / Synaptic plasticity |
| **GO LEARNING** | 140 | 0.0858 | 0.4075 | Synapse function / Synaptic plasticity |
| **REACTOME KERATINIZATION** | 155 | 0.0361 | 0.4229 | Keratinization |
| **GO KERATINIZATION** | 162 | 0.0405 | 0.4164 | Keratinization |
| **GO KERATIN FILAMENT** | 74 | 0.0569 | 0.4631 | Keratinization |
| **GO POSITIVE REGULATION OF CIRCADIAN RHYTHM** | 16 | 0.0882 | 0.6417 | - |
| **GO ENTRY OF BACTERIUM INTO HOST CELL** | 15 | 0.0165 | 0.7425 | - |

**Supplemental Table 1. Enriched gene sets across cognitively-defined AD-subgroups**

Table displays significantly enriched gene sets at p<0.10, false-discovery-rate corrected. Gene sets in bold were not enriched in the AD-No Domains group and underlined gene sets were not enriched in any of the other subgroups, indicating that these gene sets are unique to one subgroup. For AD-Language and AD-Visuospatial Functioning we marked the gene sets in yellow that were also enriched in lvPPA and PCA, respectively. Clusters were named according to shared biological functions across the gene sets in the group. FDR – false-discovery rate, ES – effect size.

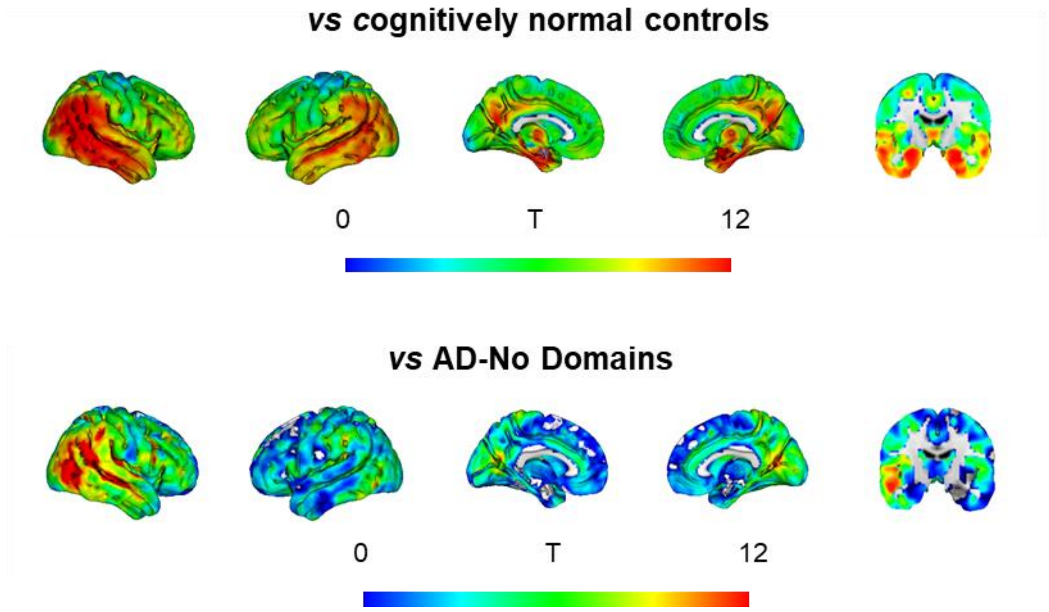

**Supplemental Figure 1. Spatial pattern of atrophy in AD-Multiple**

AD-Multiple comprised the following combinations: 49 Executive-Functioning/Visuospatial-Functioning, 15 Executive-Functioning /Language, 11 Memory/Visuospatial-Functioning, 8 Language/Visuospatial-Functioning, 2 Memory/Executive-Functioning, and 1 Executive-Functioning /Language/Visuospatial-Functioning.

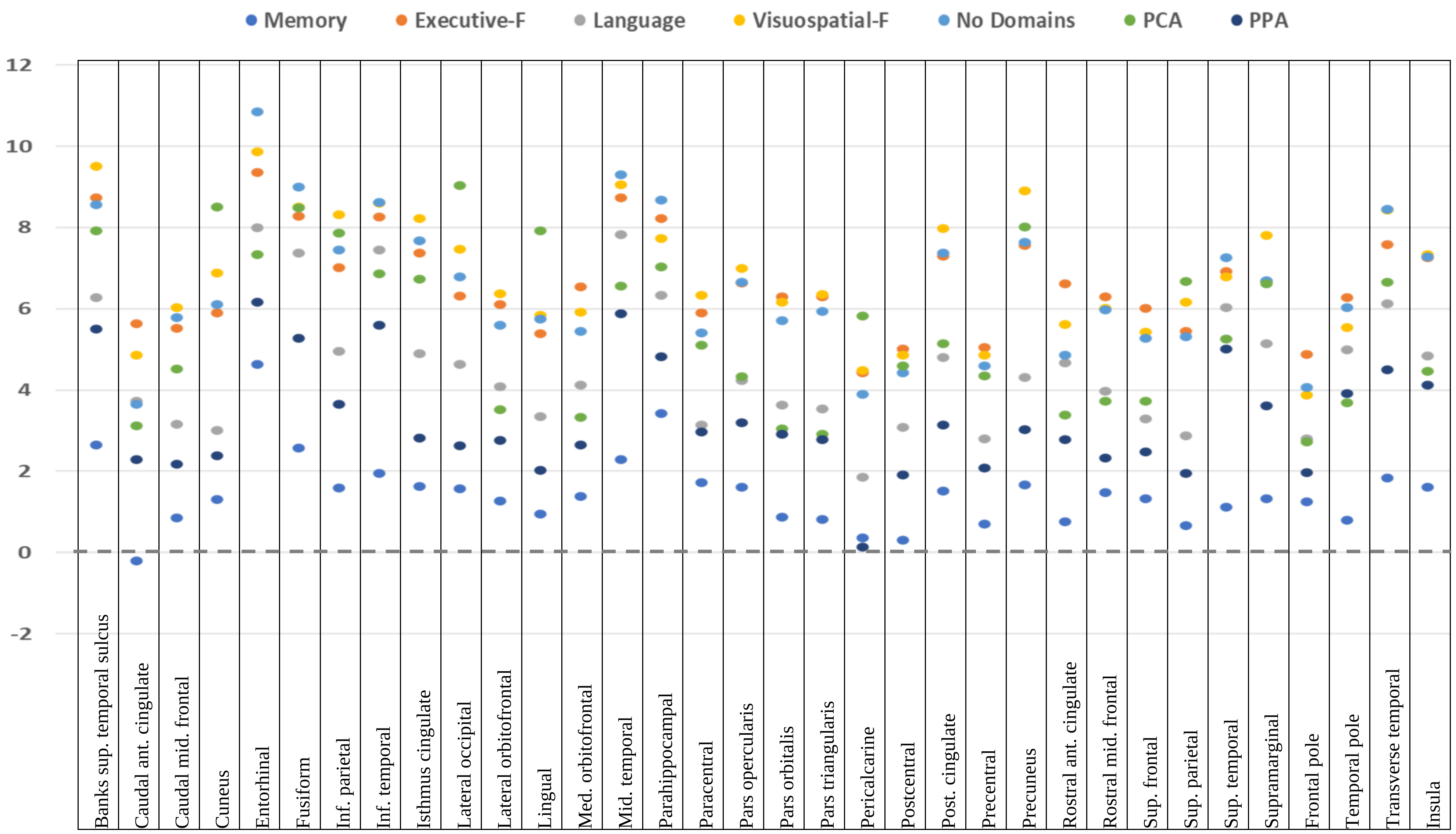

**Supplemental Figure 2. Atrophy within regions-of-interest across AD-subgroups**

Plotted values are T values within 34 regions-of-interest from the voxel-based contrasts indicating differences in gray matter volume between AD-subgroups and cognitively normal controls. Larger values indicate more atrophy.

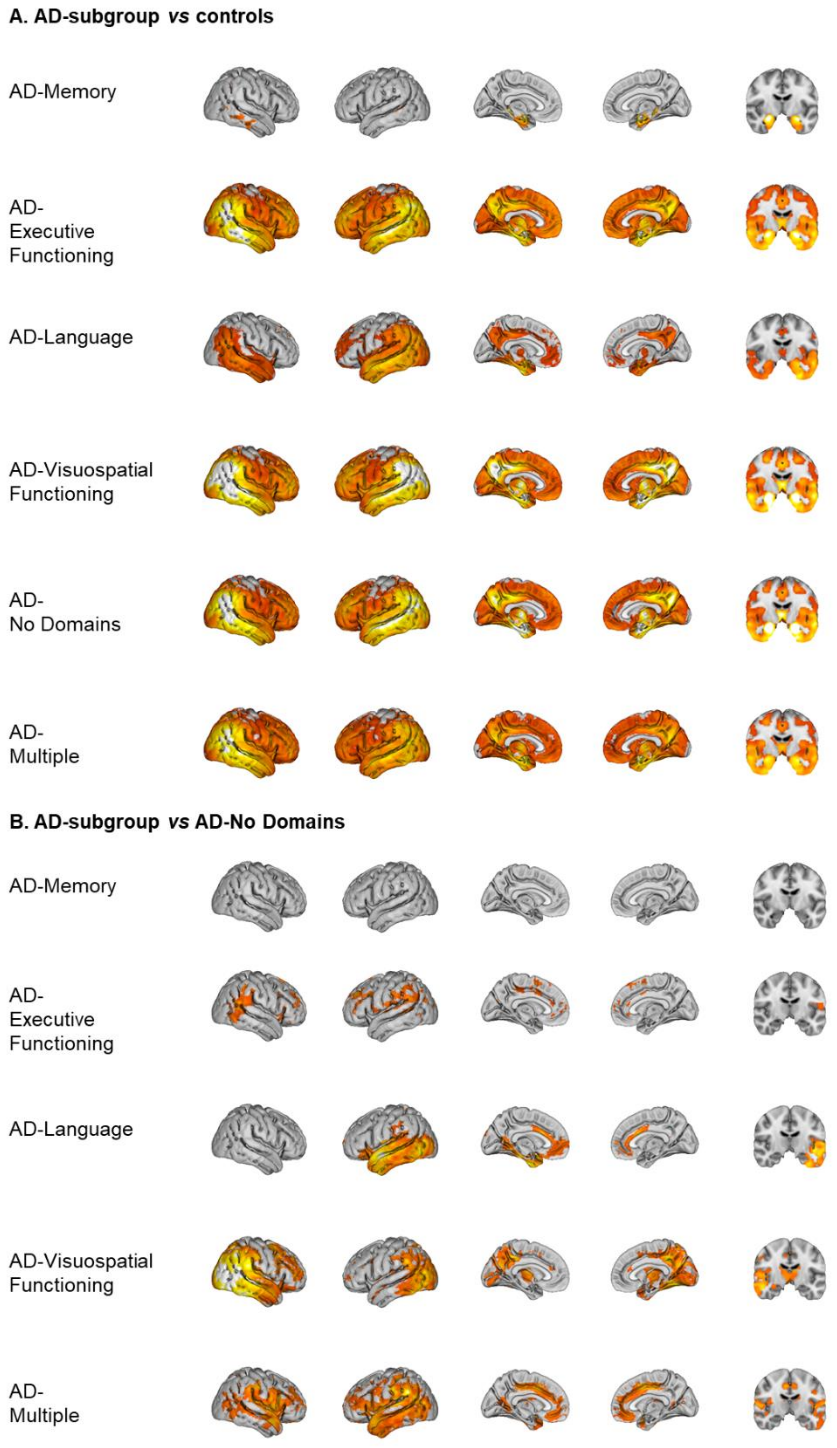

**Supplemental Figure 3. Regional atrophy across cognitively-defined subgroups**

Results from the voxel-based morphometry analyses. Only significant voxels are displayed at $p<0.05$ family-wise error corrected (panel A) and $p<0.001$ uncorrected (panel B). Colored voxels represent significant differences in gray matter volumes, adjusted for age, sex, scanner, intracranial volume (in panel A and B) and global atrophy (in panel B). Panel A displays results from voxelwise contrasts between subgroups and controls. Panel B displays results from voxelwise contrasts between subgroups and AD-No Domains

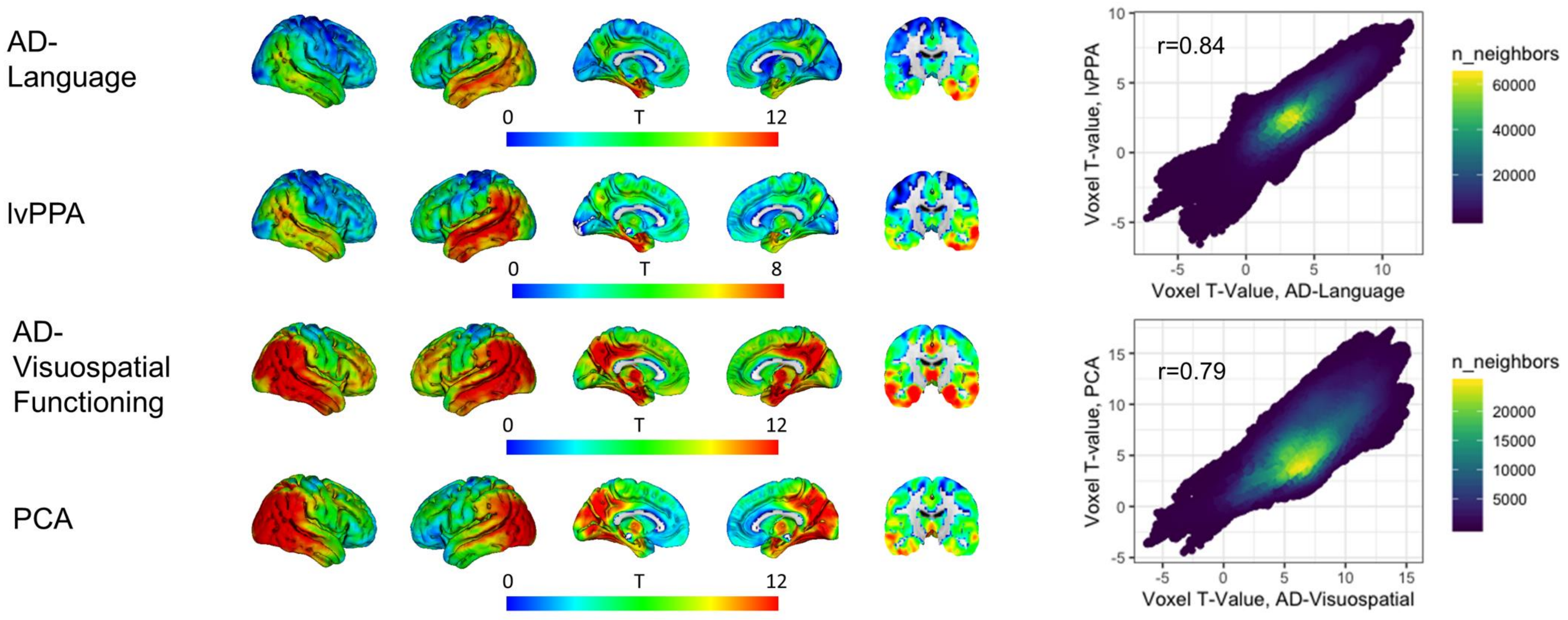

**Supplemental Figure 4. Spatial pattern of atrophy in AD-Visuospatial-Functioning and PCA, and AD-Language and lvPPA**

The point density plots display the spatial association between voxel-wise T values for contrasts against cognitively normal controls

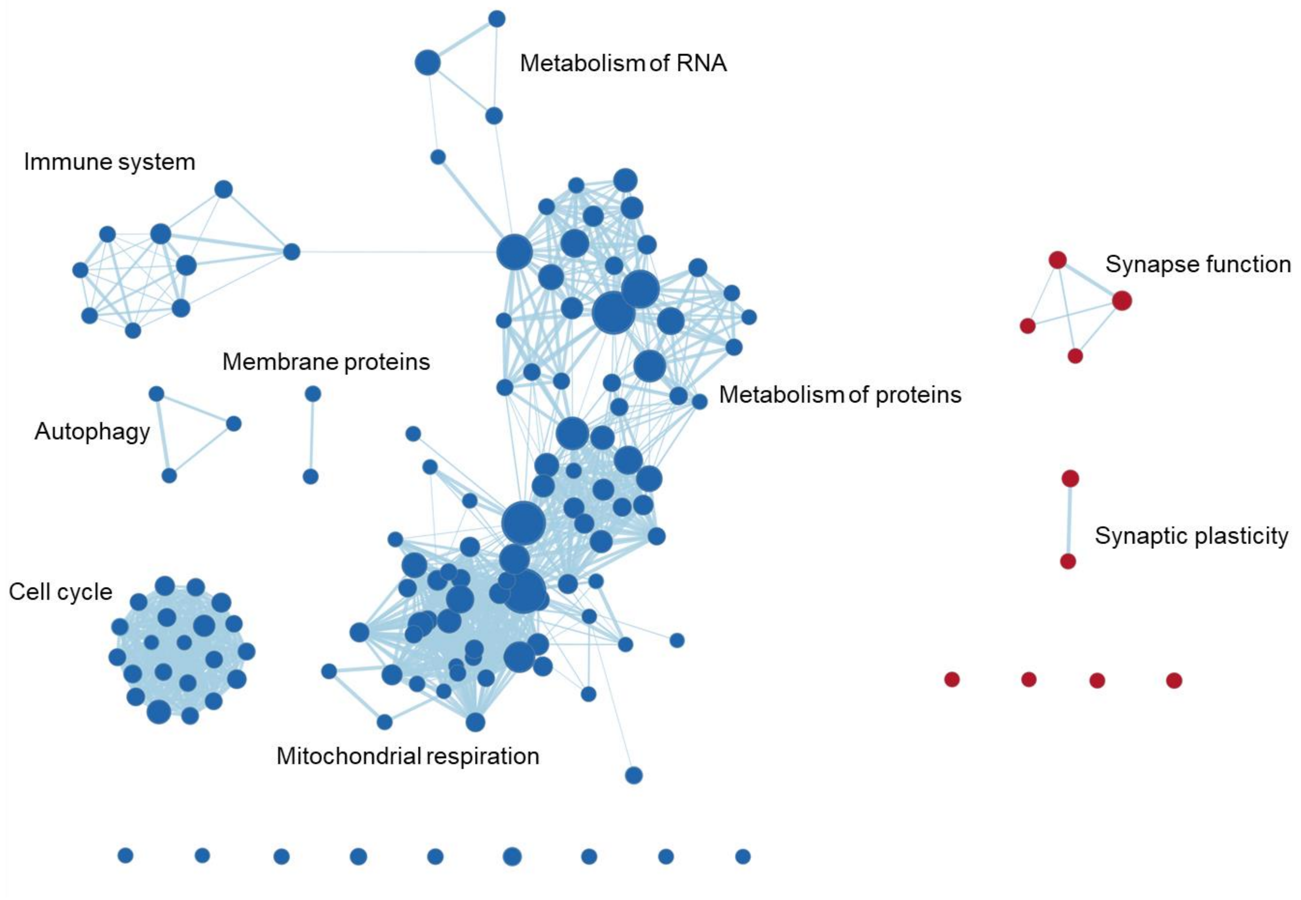

Metabolism of RNA
Immune system
Synapse function
Membrane proteins
Metabolism of proteins
Autophagy
Synaptic plasticity
Cell cycle
Mitochondrial respiration

AD-Memory

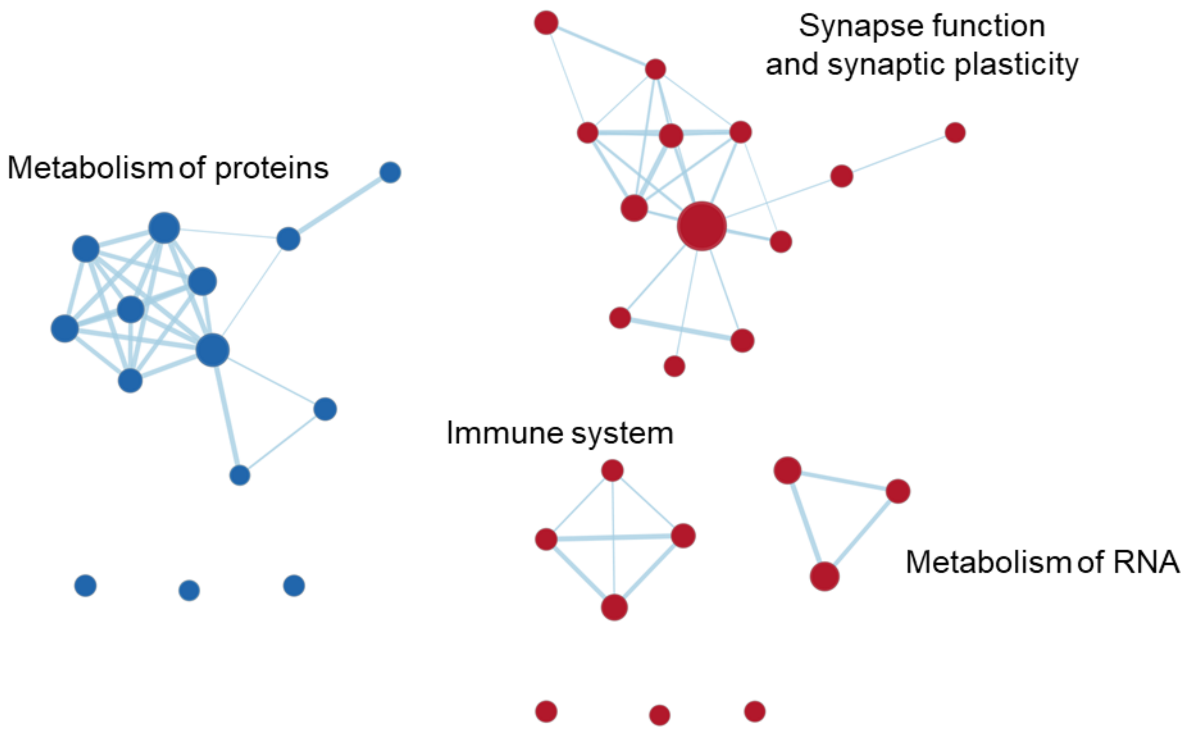

AD-Executive Functioning

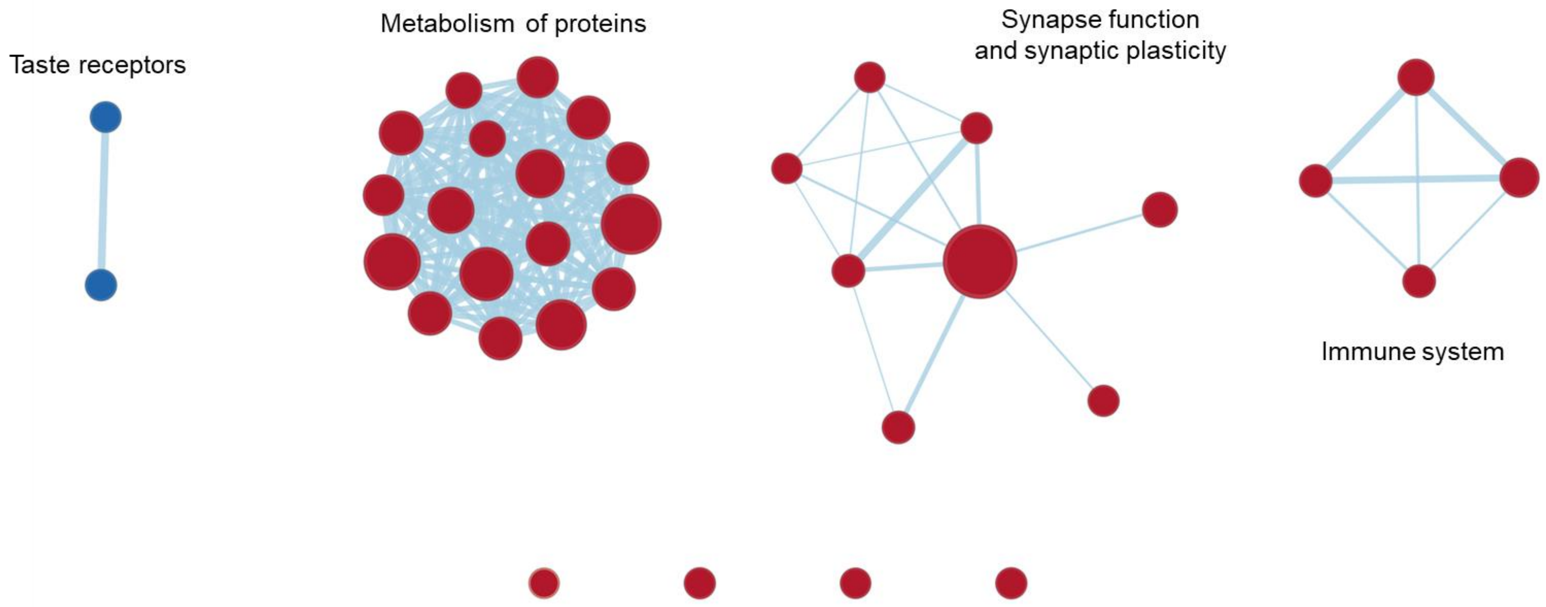
Taste receptors
Metabolism of proteins
Synapse function
and synaptic plasticity
Immune system

AD-Language

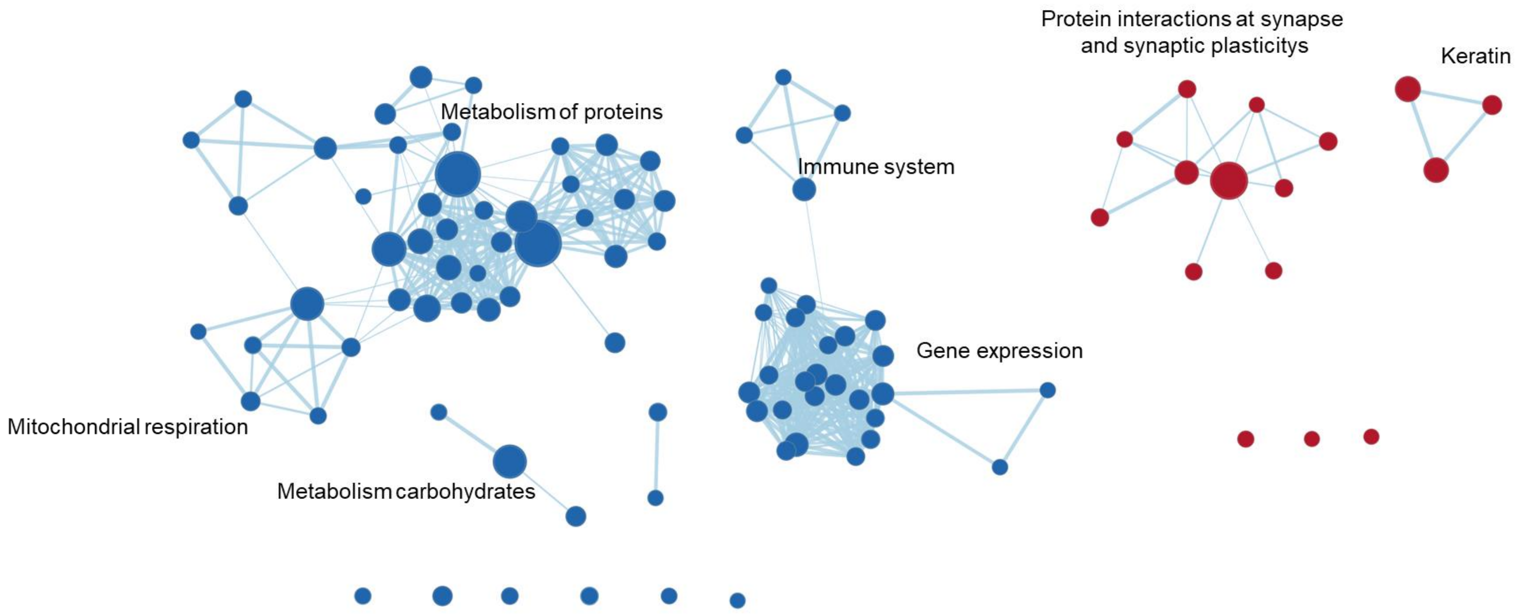

AD-Visuospatial Functioning

**Supplemental Figure 5. GSEA results displayed as clusters of genesets**

Displayed are the results from gene set enrichment analyses and the clusters represent groupings of gene sets, gene expression levels of which are spatially associated with regional atrophy patterns in AD-Memory, AD-Executive Functioning, AD-Language and AD-Visuospatial Functioning. Red nodes represent gene sets that are positively enriched (more atrophy -> higher gene expression) while blue nodes are negatively enriched. Clusters were named according to main biological functions that were commonly related to the gene sets in that grouping.

These figures display which genes (x-axis) overlapped between gene sets (y-axis) and drive the clustering results. Some gene families which showed a lot of overlap are displayed on the borders of the leading-edge results. Axis-labels are removed for visualization purposes

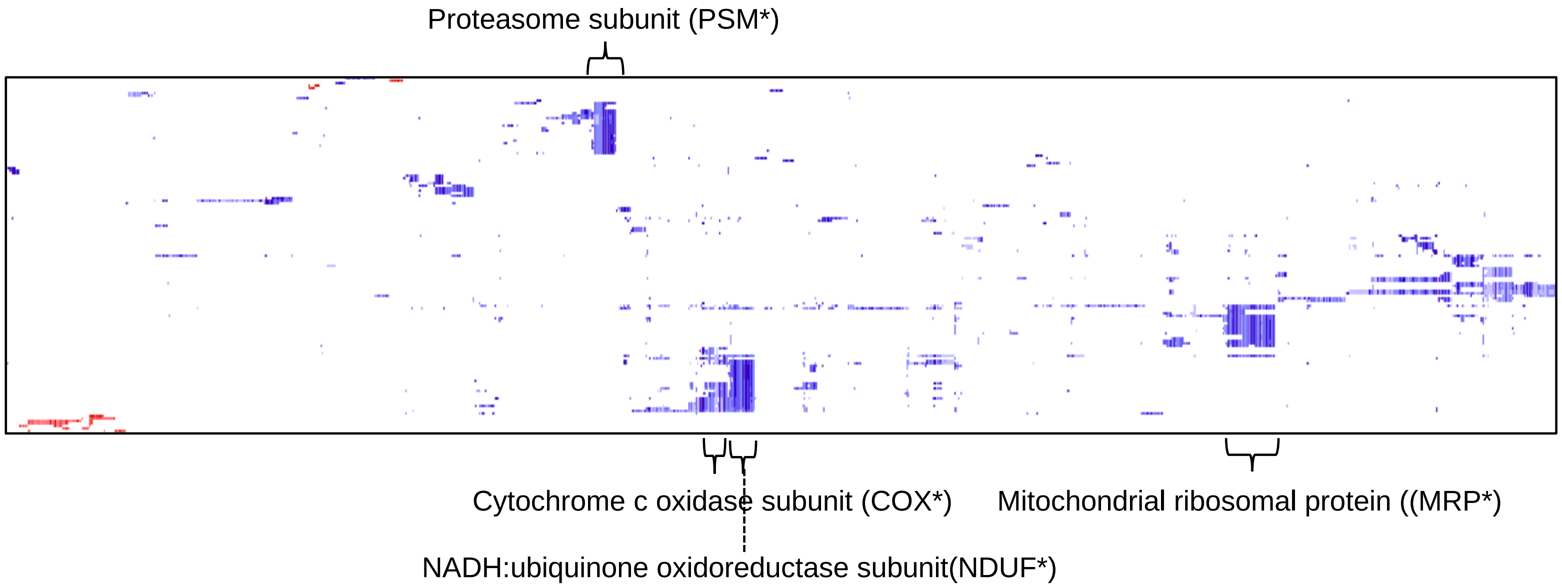

*AD-Memory***:** Four gene families were observed to show a high degree of overlap between gene sets that were negatively enriched: the mitochondrial ribosomal proteins (*MRP**; associated with protein synthesis within the mitochondrion) in the metabolism of proteins cluster, respiratory complex 1 (*NDUF**; involved in the respiratory chain) genes and cytochrome (*COX**; associated with ATP synthesis) genes in the mitochondrial respiration cluster, and proteasome (*PSM**; degrade proteins through proteolysis) in the cell cycle cluster.

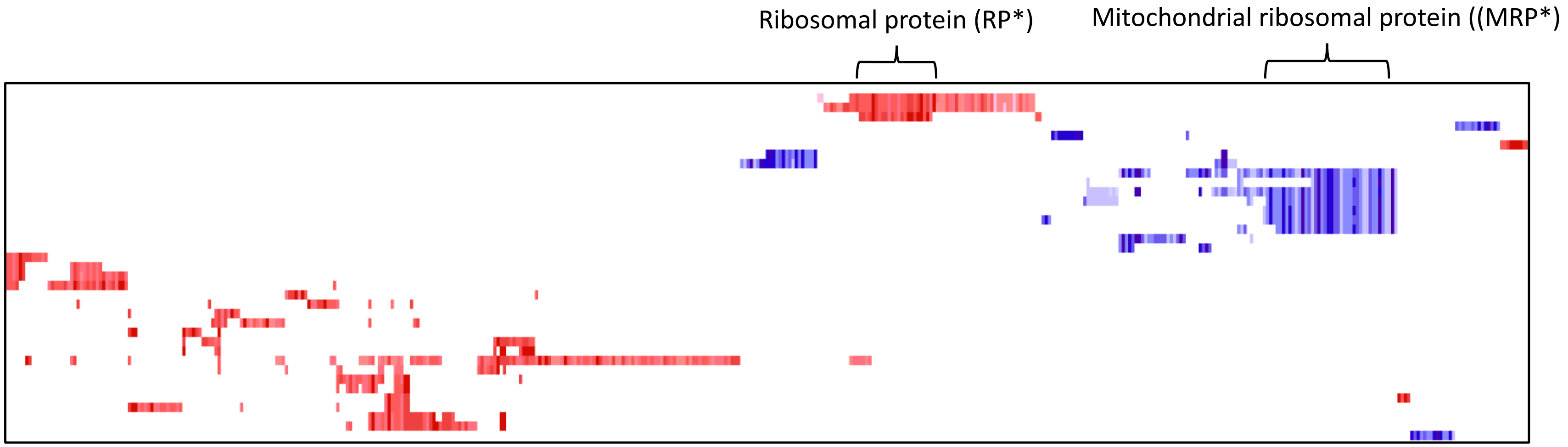

*AD-Executive:* Overlap between gene sets in the metabolism of protein cluster was again driven by the *MRP** family, while overlap in the metabolism of RNA cluster was driven by the ribosomal protein (*RP**; coding for the subunits that make up the ribosome) gene family.

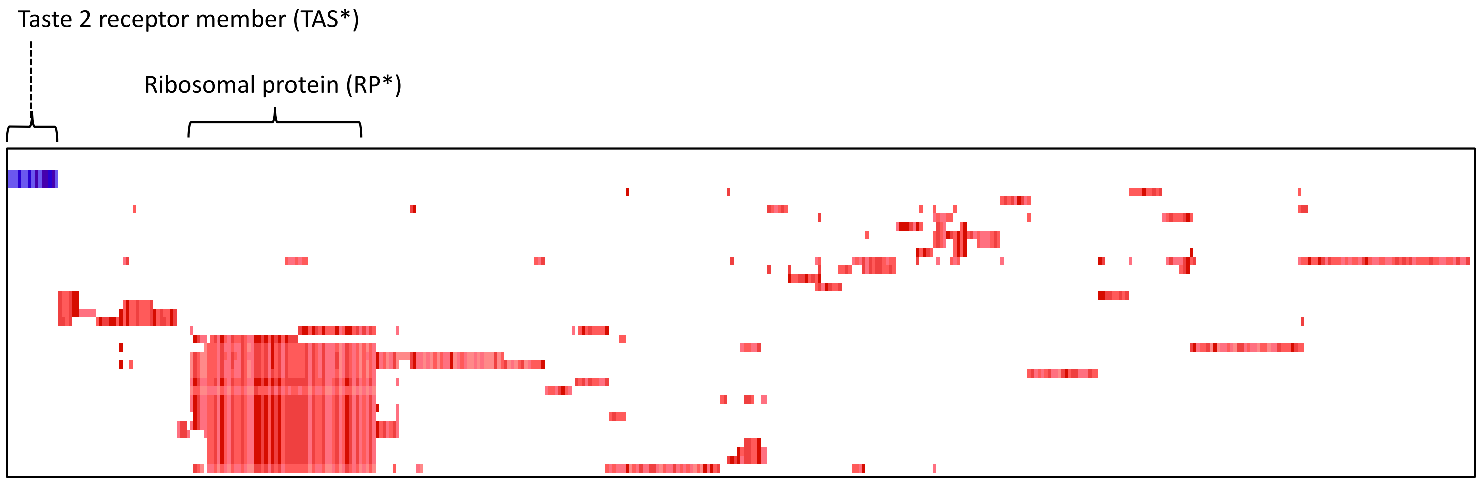

*AD-Language:* Overlap in the metabolism cluster was driven by the *RP** gene family, and in the taste receptor cluster by taste 2 receptor member (*TAS**; bitter taste receptors).

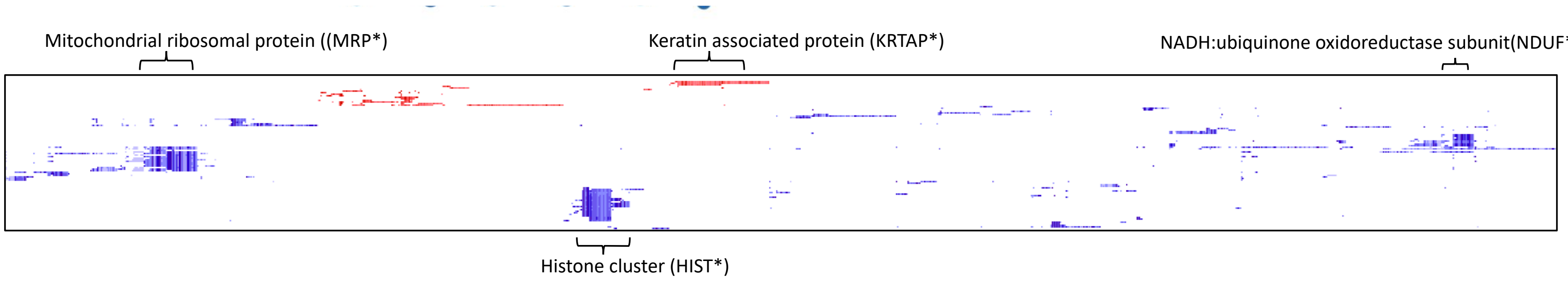

*AD-Visual:* Four gene families were observed to show a high degree of overlap between gene sets in the clusters, the mitochondrial ribosomal protein (*MRP**) family in the metabolism of protein cluster, respiratory complex 1 (*NDUF**) in the mitochondrial respiration cluster, histone cluster genes (*HIST**; making up chromatin and package DNA into nucleosomes and gene regulation) in the gene expression cluster and keratin associated proteins (*KRTAP**; keratin associated proteins) in the keratinization cluster.

**Supplemental Figure 6. Leading edge gene analysis across subgroup**

| | Cognitively defined Alzheimer's disease subgroup EOAD | | | | | | | |
|---|---|---|---|---|---|---|---|---|
| | **Total** | **AD-Memory** | **AD-Executive** | **AD-Language** | **AD-Visuospatial** | **AD-Multiple** | **AD-No Domains** | p for group effect |
| N (% of total) | 282 | 11 | 40 | 17 | 95 | 33 | 86 | |
| Age at AD diagnosis | 59.01(4.19) | 60.48(5.09) | 60.91(3.38) | 59.42(4.33) | 58.25(4.04) | 59.14(4.00) | 58.63(4.40) | 0,017[a] |
| Sex, male (%) | 129(45.7) | 5(45.5) | 20(50.0) | 9(52.9) | 42(44.2) | 21(63.6) | 32(37.2) | 0,185 |
| Edu (median [IQR])^ | 5.00[4.00,6.00] | 5.00[5.00,6.00] | 5.00[4.00,5.75] | 6.00[5.00,6.00] | 5.00[4.00,6.00] | 5.00[4.00,6.00] | 5.00[4.00,6.00] | 0,146 |
| *APOE*ε4 positive (%) | 185(65.6) | 10(90.9) | 26(65.0) | 10(58.8) | 61(64.2) | 20(60.6) | 58(67.4) | 0,542 |
| MMSE | 20.29(5.51) | 23.82(3.09) | 17.55(5.98) | 17.35(6.30) | 21.66(5.22) | 19.61(4.44) | 20.45(5.35) | <0.001[b] |
| CSF Amyloid-β (pg/ml) | 601.81(100.13) | 637.89(104.17) | 589.69(89.76) | 590.19(124.10) | 600.94(112.32) | 599.40(82.32) | 607.36(92.35) | 0,827 |
| CSF Total tau (pg/ml) | 707.93(410.11) | 796.33(464.35) | 657.00(385.80) | 760.56(434.64) | 760.14(447.48) | 630.67(403.00) | 682.28(370.39) | 0,554 |
| CSF P-tau (pg/ml) | 86.02(34.74) | 99.00(36.23) | 84.03(36.22) | 93.75(42.38) | 88.75(33.44) | 78.70(30.57) | 83.70(35.26) | 0,497 |
| Total gray matter to intracranial volume^^ | 0.38(0.04) | 0.44(0.03) | 0.37(0.04) | 0.39(0.04) | 0.39(0.04) | 0.37(0.04) | 0.39(0.04) | <0.001[c] |

**Supplemental Table 2. Demographic and clinical characteristics of the sample**

Values depicted are mean±SD, unless otherwise indicated. Differences were assessed using ANOVA with post-hoc independent samples t-tests and $\chi^2$- tests with post-hoc Fishers exact tests, where appropriate. The Kruskal-Wallis tests was used for the non-normally distributed education variable. The AD-Multiple group was excluded from the comparisons. *APOE* – Apolipoprotein E

a – AD-Executive Functioning > AD-No Domains and AD-Visuospatial Functioning

b – AD-Executive Functioning < AD-Memory, AD-No Domains and AD-Visuospatial Functioning | AD-Language < AD-Memory and AD-No Domains

c – AD-Executive Functioning < AD-Memory, AD-No Domains and AD-Visuospatial Functioning | AD-Language < AD-Memory

^ - Assessed using the qualitative Verhage scale (range: 1-7)

^^ - Ratio of gray matter volume to total intracranial volume that indicates whole brain gray matter atrophy, lower values indicate more atrophy

**Cognitively defined Alzheimer's disease subgroup LOAD**

| | **Total** | **AD-Memory** | **AD-Executive** | **AD-Language** | **AD-Visuospatial** | **AD-Multiple** | **AD-No Domains** | p for group effect |
|---|---|---|---|---|---|---|---|---|
| N (% of total) | 397 | 30 | 77 | 16 | 76 | 53 | 145 | |
| Age at AD diagnosis | 71.39(4.22) | 71.20(4.35) | 71.70(4.39) | 69.76(2.94) | 70.55(3.82) | 71.58(4.44) | 71.81(4.29) | 0,189 |
| Sex, male (%) | 189(47.6) | 12(40.0) | 40(51.9) | 9(56.2) | 37(48.7) | 27(50.9) | 64(44.1) | 0,742 |
| Edu (median [IQR])^ | 5.00[4.00,6.00] | 5.50[4.00,7.00] | 5.00[4.00,6.00] | 5.00[4.00,6.00] | 5.00[4.00,6.00] | 5.00[4.00,6.00] | 5.00[4.00,6.00] | 0,205 |
| Edu (median [IQR])^ | 282(71.0) | 27(90.0) | 49(63.6) | 7(43.8) | 52(68.4) | 37(69.8) | 110(75.9) | 0,01[a] |
| MMSE | 21.88(4.63) | 23.63(2.79) | 21.88(4.61) | 17.12(5.23) | 23.01(4.07) | 20.19(5.56) | 22.06(4.31) | <0.001[b] |
| CSF Amyloid-β (pg/ml) | 605.49(102.80) | 597.92(87.30) | 609.48(104.34) | 566.19(100.41) | 601.14(101.50) | 594.14(109.47) | 616.33(103.11) | 0,441 |
| CSF Total tau (pg/ml) | 751.22(436.93) | 840.96(427.27) | 716.28(435.25) | 801.50(454.58) | 705.90(383.30) | 711.04(330.77) | 788.25(498.47) | 0,568 |
| CSF P-tau (pg/ml) | 92.92(41.78) | 104.69(37.20) | 88.49(38.82) | 98.75(44.59) | 89.36(38.00) | 88.82(40.58) | 95.93(46.08) | 0,428 |
| Total gray matter to intracranial volume^^ | 0.38(0.04) | 0.39(0.03) | 0.37(0.04) | 0.37(0.04) | 0.38(0.03) | 0.37(0.04) | 0.38(0.04) | 0,03[c] |

**Supplemental Table 3. Demographic and clinical characteristics of the LOAD part of the sample**

Values depicted are mean±SD, unless otherwise indicated. Differences were assessed using ANOVA with post-hoc independent samples t-tests and $\chi^2$- tests with post-hoc Fishers exact tests, where appropriate. The Kruskal-Wallis tests was used for the non-normally distributed education variable. The AD-Multiple group was excluded from the comparisons. *APOE* – Apolipoprotein E.

a – AD-Memory > AD-Executive Functioning, AD-Language and AD-Visuospatial Functioning

b – AD-Language < AD-Executive Functioning, AD-No Domains and AD-Visuospatial Functioning

c – AD-Executive Functioning < AD-Visuospatial Functioning

^ - Assessed using the qualitative Verhage scale (range: 1-7)

^^ - Ratio of gray matter volume to total intracranial volume that indicates whole brain gray matter atrophy, lower values indicate more atrophy

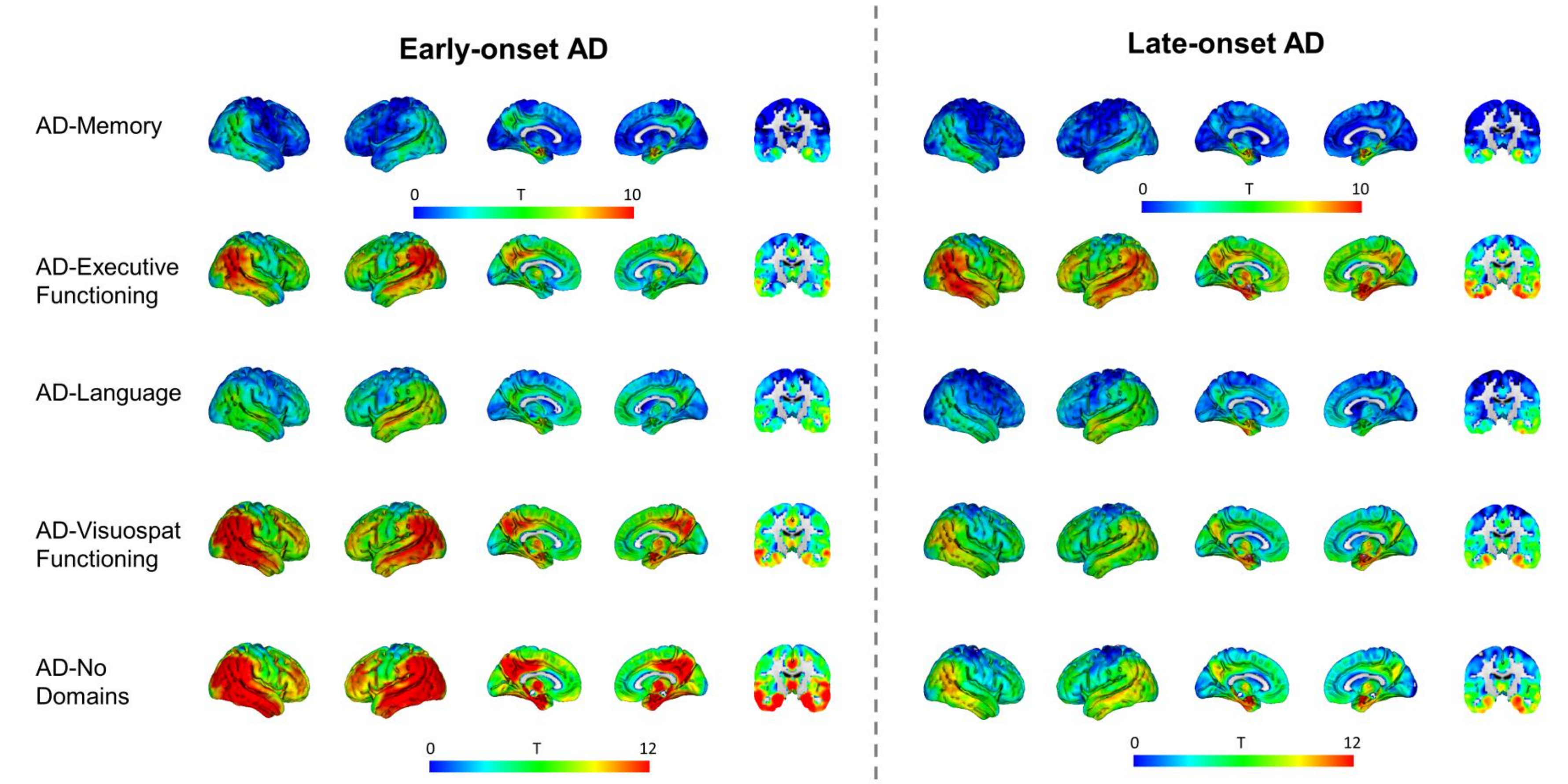

**Supplemental Figure 7. Regional atrophy across cognitively-defined subgroups stratified according to early and late-onset Alzheimer's disease**

Results from the voxel-based morphometry analyses, displayed as T-maps, representing differences in gray matter volumes between subgroups and controls adjusted for age, sex, scanner, and intracranial volume. Late-onset subjects are defined as aged > 60 at time of Alzheimer's disease diagnosis and early-onset as aged <60 years.

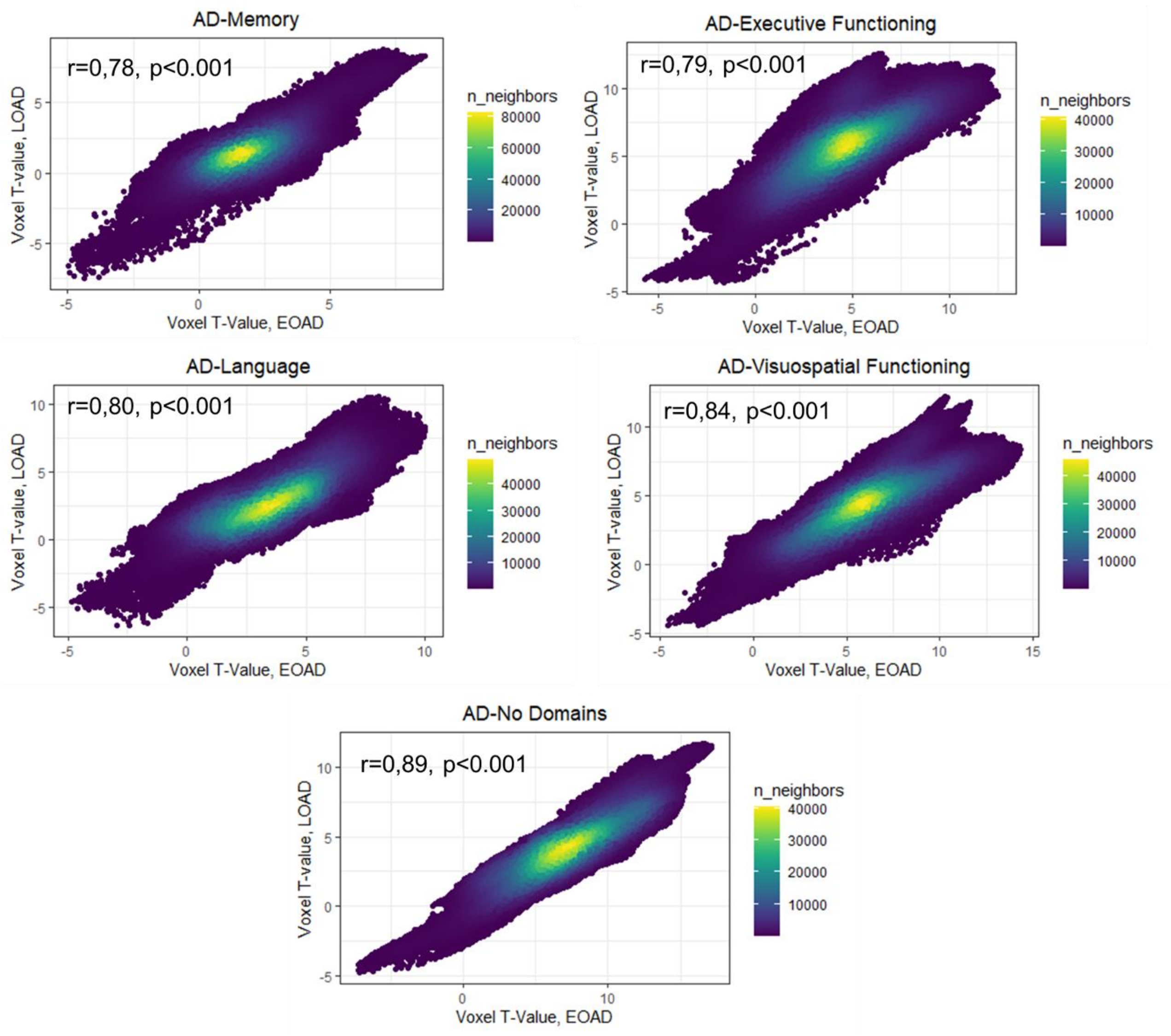

**Supplemental Figure 8. Spatial correlation between voxelwise T-maps *vs* controls of EOAD and LOAD groups.**

| **Memory** | **Executive** | **Language** | **Visual** |
| --- | --- | --- | --- |
| Rey auditory verbal learning test – immediate recall | Digit span - forward | Verbal fluency - animal naming | Number location |
| Rey auditory verbal learning test – delayed recall | Digit span - backward | Letter fluency - D,A,T | Dot counting |
| Verbal association test - A and B | Trial-making - A and B | Verbal association test - object naming | Fragmented letters |
| | Letter digit substitution test | Confrontational naming | MMSE - pentagon item |
| | Frontal assessment battery | Stroop - color | |
| | Stroop color - word | Stroop - word | |
| | | Comparative questions | |

**Supplemental Table 4. Categorization of neuropsychological tests**

Cognitive data was collected at date of AD dementia diagnosis and neuropsychological test scores were categorized into four domains according to consensus within an expert panel (P.K.C., E.T., and R.O.)

| | Cognitively defined Alzheimer's disease subgroup | | | | | | |
|---|---|---|---|---|---|---|---|
| | **Total** | **Memory** | **Executive** | **Language** | **Visual** | **Multiple** | **No domain** |
| Cognition | | | | | | | |
| Memory | 0.90 (1.03) | 0.11 (0.58) | 0.94 (1.02) | 0.55 (0.98) | 1.26 (1.07) | 1.21 (1.09) | 0.68 (0.92) |
| Executive | -0.07 (1.35) | 1.43 (0.75) | -1.13 (1.14) | -0.54 (0.96) | 0.41 (1.14) | -0.89 (1.50) | 0.20 (1.10) |
| Language | 0.76 (1.64) | 2.46 (0.98) | 0.53 (1.40) | -1.76 (1.10) | 1.15 (1.49) | 0.56 (2.17) | 0.72 (1.32) |
| Visual | -0.17 (1.24) | 1.25 (0.62) | 0.08 (1.15) | 0.05 (0.90) | -0.87 (1.13) | -0.96 (1.26) | 0.25 (1.00) |
| Average | 0.35 (1.05) | 1.31 (0.58) | 0.11 (1.04) | -0.43 (0.83) | 0.49 (1.05) | -0.02 (1.09) | 0.46 (0.96) |

**Supplemental Table 5. ACT-normalized cognitive domain Z-scores**

Values represent cognitive domain scores that are co-calibrated and z-normalized to the ACT cohort

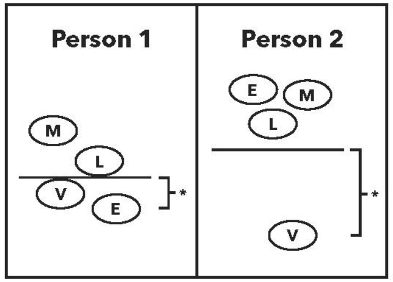

These ovals depict scores ranging from low (bottom) to high (top) for memory (M), executive functioning (E), language (L) and visuospatial functioning (V) at the time of Alzheimer's disease diagnosis. Person 1's scores are clustered closely together. Person 2's V score is much lower than their other scores. The average across domains is shown by the horizontal line. We consider differences from that average, shown by the brackets and asterisks, to place each person into a subgroup. Person 1: AD-No Domain. Person 2: AD-Visuospatial.

**Supplemental Figure 9. Graphical representation of how subgroup categorization was achieved**

| | Cognitively defined Alzheimer's disease subgroup | | | | | | |
|---|---|---|---|---|---|---|---|
| | **Total** | **Memory** | **Executive** | **Language** | **Visual** | **Multiple** | **No domain** |
| Scan (%) | | | | | | | |
| Philips PET/MR | 144 (21.2) | 12 (29.3) | 22 (18.8) | 6 (18.2) | 31 (18.1) | 16 (18.6) | 57 (24.7) |
| GE Signa | 396 (58.3) | 16 (39.0) | 68 (58.1) | 20 (60.6) | 107 (62.6) | 57 (66.3) | 128 (55.4) |
| Toshiba Titan | 139 (20.5) | 13 (31.7) | 27 (23.1) | 7 (21.2) | 33 (19.3) | 13 (15.1) | 46 (19.9) |

**Supplemental Table 6. MRI scanners used across cognitively-defined subgroups**

P for differences across groups = 0.270. Scanning parameters: GE SignaHDxt 3T (n=396, GE Healthcare, voxel size .94×.94×1mm, echo time 3ms, repetition time 7.8ms, flip angle 12°, field of view 240mm), Toshiba Vantage Titan 3T (n=139, Toshiba medical systems, voxel size 1×1×1mm, echo time 3.2ms, repetition time 9.5ms and flip angle 7°, field of view 256mm) and Philips Ingenuity TF PET–MRI 3T (n=144, Philips medical systems, voxel size .87×.87×1mm, echo time 3ms, repetition time 7ms, flip–angle 12°, field of view 250mm).

**Supplemental Text 1. Psychometric analyses**

What follows is a detailed description of how co-calibration of neuropsychological scores was achieved. First, we will describe how scores from ACT, ADNI, ROS/MAP and PITT were co-calibrated to form the legacy cohort. Then, we will describe the co-calibration of the scores from the ADC cohort to this legacy cohort.

*Confirmatory factor analyses in each study*

**Step 1: Domain assignment:** In each of the studies (Adult Changes in Thought [ACT], Alzheimer's Disease Neuroimaging Initiative [ADNI], the Religious Orders Study–Memory and Aging Project (ROS/MAP], and the University of Pittsburgh data set [PITT]), the expert panel (Dr. Trittschuh, Dr. Mez, and Dr. Saykin) assigned items from the neuropsychological battery to one of the four domains (memory, language, executive functioning, and visuospatial ability); other items did not map to any of these domains. The expert panel also assigned each of these items to sub-domains based on the cognitive processes involved in each task. We also noted methods effects where the same stimulus was used in multiple assessments. We also used a data-driven approach looking at patterns of responses among participants to identify alternate possible secondary domain structures.

**Step 2: Data quality control:** Each of the studies sent their neuropsychological data sets to our team and Ms. Sanders, our data manager, ran an initial quality control on these. Ms. Sanders prepared a data set which included item-level data for individuals at their first Alzheimer's disease diagnosis. Before running psychometric models, we performed additional recoding of the data. Some items such as Trails A and B were reverse coded. We checked each item to make sure lower values represent lower cognitive performance. We considered the distribution of each item among those with non-missing data and combined categories as needed. Our goals were a.) to avoid sparse categories (operationally defined as <5 responses for each study administering each item) and b.) to have a maximum of 10 categories, which is the maximum number of categories handled by Mplus v7.4. We treated each item as an ordinal indicator of the domain—the numerical value assigned to each category is irrelevant beyond its rank, e.g. calling the lowest category 3 points vs. 18 points makes no difference in how the item is treated or what the final score would be.

We also looked at informative missingness in each study and recoded relevant items accordingly. For example, some of the studies include multiple missing codes, where it was possible to identify refusal to respond to an item as opposed to the interviewer ran out of time and the item was never administered. The first of these—

refusal—we took as informative missing and assigned that code to the lowest response category, while the second of these—missing due to scheduling etc.—we took as non-informative missing and omitted that item from consideration.

**Step 3: Confirmatory factor analyses:** We then turned to confirmatory factor analysis modeling with Mplus using a Robust Weighted Least Squares including terms for the mean and the variance (WLSMV) estimator. We ran four models: a.) a single factor model, with no residual structure; b.) A theory-driven cognitive process bifactor model, using the a priori sub-domain assignments; c.) a theory-driven methods effects bifactor model, using the "methods effects" assignments; and d.) a data-driven bifactor model, using hierarchical clustering-assigned sub-domains. We consulted the expert panel on the sub-domain assignment of items in our data-driven approach to make sure these models made sense to our experts. Our overall strategy was that we would choose the single factor model if adding secondary factors did not markedly improve model fit and if adding secondary factors did not markedly impact any individual's score (see below).

Our goal with the three bifactor models (models b, c, and d) was to identify a single candidate bifactor model to compare with the single-factor model (model a). Our criteria for selecting the candidate bifactor model included fit statistics (see below) and concordance of model results with theory, such as all loadings on secondary factors being positive. The fit statistics we considered were the confirmatory fit index (CFI) where higher values indicate better fit; thresholds of 0.90 and 0.95 have been used in other settings as criteria for adequate or good fit; the Tucker-Lewis Index (TLI), which has similar criteria as the CFI; and the root mean squared error of approximation (RMSEA), where lower values indicate better fit, and thresholds of 0.08 and 0.05 have been used in other settings as criteria for adequate or good fit.

When comparing the single factor model with the best bifactor model, we a) looked at whether loadings on the primary factor were within 10% of each other across the two models and b) compared the scores for the single factor model vs. scores for the final candidate bifactor model. We used as our threshold a difference of 0.30 units. We chose this value based on the default stopping rule for computerized adaptive testing; this has been used for years as the default level of tolerable measurement differences in the setting of computerized adaptive tests. While arbitrary, this is a level of ambiguity that has been thought to be tolerable in a variety of situations. If there were a substantial number of people for whom the differences in scores were larger than 0.3 from each other, and if the bifactor model conformed to our theory better and had better fit statistics, we selected the final candidate bifactor model as our choice for modeling a domain.

*Co-calibration of the domains across ACT, ADNI, and ROS/MAP*

**Step 1: Identification of anchor items:** Co-calibration requires either the same people taking different tests or different tests sharing common items. Here we had common items. We identified candidate anchor items with identical content across tests administered in different studies and ensured that their relationship with the underlying ability tested was the same across studies by performing preliminary confirmatory factor analysis models within each study. These items were then used to anchor the scales in each domain to a common metric. We consulted the expert panel (Dr. Trittschuh) to make sure we chose the anchor items correctly.

**Step 2: Quality control for anchor items:** Anchor items were cleaned and recoded after merging in the items from all the studies making sure that the range of the anchor items were similar in each study. We carefully reviewed documentation from each study to ensure that the stimulus was precisely the same, that the response options were precisely the same or could be re-coded to be the same, and that we were mapping data from each study in a way that the same response would result in the same score regardless of which study the person was enrolled in.

A note regarding response options—in many cases the stimulus is fairly open-ended, such as “can you please draw from memory the figure you copied a while ago”, where the participant is handed a blank sheet of paper and a writing implement. The resulting drawing then gets scored based on how similar it was to the initial stimulus figure. The specific scoring applied to such a stimulus could vary across studies. One study could score such an item as correct vs. incorrect, while another could apply points for various aspects of the drawing. We reviewed the scoring documentation from both studies to determine what “correct” meant in the first study, and how many aspects of the drawing would need to be present for a “correct” score in that study. Then we would map all scores from the second study that would have resulted in a “correct” score in the first study to a “correct” score, and all other scores from the second study to an “incorrect” score. In this way, the resulting score is invariant to which study the person is participating in, as each response would be consistently scored regardless of study.

**Step 3: Confirmatory factor analyses:** We co-calibrated each of the four domains (memory, executive functioning, language, and visuospatial ability) by incorporating the components of the best model in each study (i.e., the final single-factor or bifactor model selected as described above) into one mega-calibration model. One particularly tricky aspect of co-calibrating scores using bifactor models is how to handle secondary domains. Some anchor items had loadings on the primary domain (e.g. memory) and also on a secondary domain. That structure by itself does not lead to conceptual problems. However, item representation of the secondary domain may vary across studies, with variable numbers of items, and potential missing data and identifiability issues. To address this we used robust maximum likelihood (MLR) estimation that is robust to missing data, and assigned all subdomain indicators across studies to the same subdomain. Unlike running a CFA model with the WLSMV estimator, a CFA model with MLR estimator does not output fit statistics like CFI/TLI/RMSEA. For our purposes, these secondary domains were nuisances. We performed a number of sensitivity analyses to reassure ourselves that scores on the primary domain were minimally impacted by various ways of specifying the mean and variance on secondary domains. In the final models we selected, we specified a mean of 0 and a variance of 1 for each secondary domain factor, regardless of the number of studies that included items that loaded on that factor.

Once we had fit the final mega-calibration model for each domain, we extracted factor scores for the primary factor (e.g. memory). The resulting scores are on the same metric with a mean of 0 and variance of 1. We used all participants with relevant data to fit data for each domain, so different the scale for each domain was based on models that included different specific people, since some people were missing for some domains. We therefore picked a reference population for standardizing scores for each domain. We used ACT for this, as it was a community-based prospective cohort study, and had a very large sample (n=825) of people with sufficient cognitive data to generate all of our scores. We applied the same standardization to all participants for each study.

Thus, a score of 0, regardless of study, reflects the mean for people with Alzheimer's disease in the ACT study; and a score of -1, regardless of study, reflects 1 SD below the mean for people with Alzheimer's disease in the ACT study.

For future data sets in our pipeline such as University of Pittsburgh (PITT) and the Amsterdam Dementia Cohort (ADC), we used these estimated thresholds and loadings of items from the mega-calibration models to obtain scores for individuals. New items (not part of previously co-calibrated studies) were freely estimated while already seen items will have their parameters fixed based on these mega-calibration models.

Confirmatory factor analysis model considerations in co-calibration models

**1.** For all CFA models, we categorized items to ≤ 10 categories. For co-calibration purpose, we had to re-categorize some of the items even though they already had ≤ 10 categories. This was because some studies had more granular data (more categories) for anchor items compared to other studies. In these cases, after we estimated item parameters from the co-calibration model, we re-estimated parameters of the anchor item(s) in the most granular form in the given study. For example, the item “q20mme” was an anchor item for visuospatial ability administered in ACT, ROS/MAP, and ADNI. The item asks individuals to copy intersecting pentagons. In ROS/MAP and ADNI, this item is coded as 0/1 (incorrect/correct) while in ACT it is coded 0–10 (four points for aspects of the left pentagon, four points for aspects of the right pentagon, two points for aspects of the intersection). For co-calibration purpose, we dichotomized this item to 0/1. We consulted scoring algorithms for each of the studies to determine that only scores of 10/10 from the ACT study would have received scores of 1 from ROS/MAP or ADNI; any drawing receiving a score of 9 or fewer from ACT would have scored a 0 in the other studies.

After using re-coded items for co-calibration, we fixed all of the other items to their values from the co-calibration run and freely estimated parameters for re-coded anchors in their most granular form. This approach enabled us to obtain more precise scores in studies that incorporated more granular scoring rules, while still using all items administered across studies to co-calibrate metrics across studies.

**2.** The base co-calibration exercise for each of the four domains was performed across ACT, ROS/MAP, and ADNI. PITT and ADC data were subsequently added with the following steps. For each domain, we identified anchor items and fixed their item parameters to those estimated previously in the base co-calibration models; unique items administered in the new study that were not administered to people in previously co-calibrated studies were freely estimated.

In these models,

a) The mean and variance for the primary factor were freely estimated.

b) If every item in a sub-domain in the new data had parameters available from the co-calibration model, we fixed those item parameters to their previously identified values, and allowed the mean and variance to be freely estimated in the new data.

If no item from a sub-domain had parameters available, then we freely estimated each of the sub-domain loadings, fixing the mean and variance of the subdomain factor to 0 and 1.

If there was a mix of previously specified and new items in a subdomain, we fixed the parameters for the previously specified items, and allowed the mean and variance of the factor and the loadings for new items to be freely estimated in the new data.

**NOTE:** Detailed overview and all code snippets can be obtained from authors on request.

Neuropsychological items by domain for each study and fit statistics from CFA models

MEMORY

ACT: Final model was a theory driven methods-effects bifactor model with CFI = 0.923, TLI = 0.914, and RMSEA = 0.052. The following items were included in the CFA analysis (Supplemental Table 6).

**Supplemental Table 6. Items and secondary structure for memory for the ACT study**

| Study | Variable | Description | Secondary Structure |
|---|---|---|---|
| ACT | mat_mem | Mattis Dementia Rating Scale Memory score | |
| ACT | w_in_c1 | Word list learning trial 1 total score | F1 |
| ACT | w_in_c2 | Word list learning trial 2 total score | F1 |
| ACT | w_in_c3 | Word list learning trial 3 total score | F1 |
| ACT | w_rcl_c | Word List Recall—correct | F1 |
| ACT | w_rcg_t | Word Recognition—total correct | F1 |
| ACT | cp_re_ci | Constructional Praxis Delay—circle | F2 |
| ACT | cp_re_di | Constructional Praxis Delay—diamond | F2 |
| ACT | cp_re_re | Constructional Praxis Delay—rectangles | F2 |
| ACT | cp_re_cu | Constructional Praxis Delay—cube | F2 |
| ACT | w_lm_ima | Logical Mem I—immediate recall total story A | F3 |
| ACT | w_lm_imb | Logical Mem I—immediate recall total story B | F4 |
| ACT | w_lm_dea | Logical Mem II—delayed recall total story A | F3 |
| ACT | w_lm_deb | Logical Mem II—delayed recall total story B | F4 |
| ACT | w_vp_ine | Verbal Paired Associates I easy | F5 |
| ACT | w_vp_inh | Verbal Paired Associates I hard | F6 |
| ACT | w_vp_ree | Verbal Paired Associates II easy | F5 |
| ACT | w_vp_reh | Verbal Paired Associates II hard | F6 |
| ACT-CASI | rgs1 | repeat words | F7 |
| ACT-CASI | rc1a | Word recall—something to wear—1 | F7 |

| | | | |
|---|---|---|---|
| ACT-CASI | rc1b | Word recall—a color—1 | F7 |
| ACT-CASI | rc1c | Word recall—personal quality—1 | |
| ACT-CASI | yr | What is today's date?—year | F8 |
| ACT-CASI | mo | What is today's date?—month | F8 |
| ACT-CASI | casi_dat | What is today's date?—day | F8 |
| ACT-CASI | day | What day of week? | F8 |
| ACT-CASI | casi_ssn | What season is it? | F8 |
| ACT-CASI | spa | What state and city? | F8 |
| ACT-CASI | spb | What is this place? | F8 |
| ACT-CASI | rc2a | Word recall—something to wear—2 | F7 |
| ACT-CASI | rc2b | Word recall—a color—2 | F7 |
| ACT-CASI | rc2c | Word recall—personal quality—2 | F7 |
| ACT-CASI | rcobj | Recall of 5 objects | |

ADNI: Final model was a data driven bifactor model with CFI = 0.951, TLI = 0.946, and RMSEA = 0.036. The following items were included in the CFA analysis (Supplemental Table 7).

**Supplemental Table 7. Items and secondary structure for memory for the ADNI study**

| Study | Variable | Description | Secondary Structure |
|---|---|---|---|
| ADNI | limmtotal | Logical Memory—Immediate Recall | F1 |
| ADNI | ldeltotal | Logical Memory—Delayed Recall | F1 |
| ADNI | avtot1* | AVLT Trial 1 Total | F2 |
| ADNI | avtot2* | Trial 2 Total | F2 |
| ADNI | avtot3* | Trial 3 Total | F2 |
| ADNI | avtot4* | Trial 4 Total | F2 |
| ADNI | avtot5* | Trial 5 Total | F2 |
| ADNI | avtot6* | Trial 6 Total | F3 |
| ADNI | avtotb* | List B Total | F2 |

| | | | |
|---|---|---|---|
| ADNI | avdel30min* | 30 Minute Delay Total | F3 |
| ADNI | avdeltot* | Recognition Score | F4 |
| ADNI | q1score | ADAS Word Recall—score | F2 |
| ADNI | q4score | ADAS Delayed Word Recall | F4 |
| ADNI | q7score | ADAS Orientation—score | F5 |
| ADNI | q8score | ADAS Word Recognition—score | |
| ADNI | mmdate | What is today's date? | F5 |
| ADNI | mmyear | What is the year? | F5 |
| ADNI | mmmonth | What is the month? | F5 |
| ADNI | mmday | What day of the week is today? | F5 |
| ADNI | mmseason | What season is it? | |
| ADNI | mmhospit | What is the name of this hospital (clinic, place)? | |
| ADNI | mmfloor | What floor are we on? | |
| ADNI | mmcity | What town or city are we in? | |
| ADNI | mmarea | What county (district, borough, area) are we in? | |
| ADNI | mmstate | What state are we in? | |
| ADNI | mmball | Ball | F6 |
| ADNI | mmflag | Flag | F6 |
| ADNI | mmtree | Tree | F6 |
| ADNI | mmballdl | Ball delayed | F7 |
| ADNI | mmflagdl | Flag delayed | F7 |
| ADNI | mmtreedl | Tree delayed | F7 |
| ADNI | imm1sum | Immediate recall of the MoCA list (#1) | F2 |
| ADNI | imm2sum | Immediate recall of the MoCA list(#2) | F2 |
| ADNI | delsum | Delayed recall of the MoCA list | |

MoCA (blue) items were only administered in ADNI GO/2 while orange items were in all ADNI waves (1/GO/2).

ADNI administered two versions (different word lists) of RAVLT (avtot1–avdeltot) and three different versions of ADAS-Cog items (q*) across waves. We ran the model separately for the two versions. The ADAS-Cog versions were found to be equivalent while the RAVLT versions were not. For determining secondary factor structures and extracting model fit statistics, we considered all RAVLT versions to be equivalent. The different versions of RAVLT were taken into account in the final co-calibration phase.

There were additional MoCA items, which were the same (theoretically) as corresponding items from the Mini-Mental State Examination (MMSE). We excluded MoCA items if those items were already asked as part of the neuropsychological battery.

ROS/MAP: Final model was a data driven bifactor model with CFI = 0.941, TLI = 0.929, and RMSEA = 0.063. The following items were included in the CFA analysis (Supplemental Table 8):

**Supplemental Table 8. Items and secondary structure for memory for the ROS and MAP studies**

| Study | Variable | Description | Comments | Secondary Structure |
|---|---|---|---|---|
| ROS/MAP | Q1mme | What is the year? | | F1 |
| ROS/MAP | Q2mme | What is the season of the year? | | |
| ROS/MAP | Q3mme | What is the date? | | F1 |
| ROS/MAP | Q4mme | What is the day of the week? | | F1 |
| ROS/MAP | Q5mme | What is the month? | | F1 |
| ROS/MAP | Q6mme | What state are we in? | | F2 |
| ROS/MAP | Q8mme | What city are we in? | | F2 |
| ROS/MAP | Q7mme | What county are we in? | | F2 |
| ROS/MAP | Q9mme | What room are we in? | | F2 |
| ROS/MAP | Q10amme | What is the address of this place? | | F2 |
| ROS/MAP | Q10bmme | Street Name | | F2 |
| ROS/MAP | atb1 | Apple, table, penny (immediate) | 3 items collapsed | |
| ROS/MAP | story | Logical memory | | F3 |
| ROS/MAP | WordT1 | Word list learning Trial 1 | 10 items collapsed | F4 |
| ROS/MAP | WordT2 | Word list learning Trial 2 | “ | F4 |

| | | | | |
|---|---|---|---|---|
| ROS/MAP | WordT3 | Word list learning Trial 3 | " | F4 |
| ROS/MAP | WordRec | Which one of these words is from that list? (Word list recognition) | 10 items collapsed | F6 |
| ROS/MAP | Recall | Word list recall | 10 items collapsed | F6 |
| ROS/MAP | ebmt | East Boston immediate recall | 12 items collapsed | F5 |
| ROS/MAP | atb2 | apple, table, penny (delayed) | 3 items collapsed | |
| ROS/MAP | ebdr | East Boston delayed recall | 12 items collapsed | F5 |
| ROS/MAP | Delay | Tell me the story again | | F3 |

**Supplemental Table 9. Co-calibration of memory across ACT, ADNI, ROS/MAP**

| Study | Variable | Secondary structure | Comments |
|---|---|---|---|
| ACT, ADNI, ROS/MAP | mmyear | F2 | Q1mme in ROS/MAP; yr in ACT |
| ACT, ADNI, ROS/MAP | mmseason | | Q2mme in ROS/MAP; casi_ssn in ACT |
| ACT, ADNI, ROS/MAP | mmdate | F2 | Q3mme in ROS/MAP; casi_dat in ACT |
| ACT, ADNI, ROS/MAP | mmday | F2 | Q4mme in ROS/MAP; day in ACT |
| ACT, ADNI, ROS/MAP | mmmonth | F2 | Q5mme in ROS/MAP; mo in ACT |
| ACT, ADNI, ROS/MAP | mmctst | F6 | Collapsed (Q6mme Q8mme) in ROSMAP and (mmcity mmstate) in ADNI to create a single variable; spa in ACT |
| ACT, ADNI | limmtotal | F3 | limmtotal in ADNI; w_lm_ima in ACT |
| ACT, ADNI | ldeltotal | F3 | limmtotal in ADNI; w_lm_dea in ACT |
| ROS/MAP | Q7mme | F6 | |
| ROS/MAP | Q9mme | F6 | |
| ROS/MAP | Q10amme | F6 | |
| ROS/MAP | Q10bmme | F6 | |
| ROS/MAP | atb1 | | |
| ROS/MAP | story | F9 | |

| | | |
|---|---|---|
| ROS/MAP | WordT1 | F7 |
| ROS/MAP | WordT2 | F7 |
| ROS/MAP | WordT3 | F7 |
| ROS/MAP | ebmt | F8 |
| ROS/MAP | WordRec | F10 |
| ROS/MAP | atb2 | |
| ROS/MAP | Recall | F10 |
| ROS/MAP | ebdr | F8 |
| ROS/MAP | Delay | F9 |
| ACT | mat_mem | |
| ACT | w_in_c1 | F11 |
| ACT | w_in_c2 | F11 |
| ACT | w_in_c3 | F11 |
| ACT | w_rcl_c | F11 |
| ACT | w_rcg_t | F11 |
| ACT | cp_re_ci | F12 |
| ACT | cp_re_di | F12 |
| ACT | cp_re_re | F12 |
| ACT | cp_re_cu | F12 |
| ACT | w_lm_imb | F14 |
| ACT | w_lm_deb | F14 |
| ACT | w_vp_ine | F15 |
| ACT | w_vp_inh | F16 |
| ACT | w_vp_ree | F15 |
| ACT | w_vp_reh | F16 |
| ACT-CASI | rgs1 | |
| ACT-CASI | rc1a | F13 |
| ACT-CASI | rc1b | F13 |
| ACT-CASI | rc1c | F13 |

| | | | |
|---|---|---|---|
| ACT-CASI | spb | F6 | |
| ACT-CASI | rc2a | F13 | |
| ACT-CASI | rc2b | F13 | |
| ACT-CASI | rc2c | F13 | |
| ACT-CASI | rcobj | | |
| ADNI | avtot1 | F1 | Each RAVLT item was split into two items to account for two versions of RAVLT used in ADNI at specific waves where both versions of the same item were loaded into the same secondary structure |
| ADNI | avtot2 | F1 | |
| ADNI | avtot3 | F1 | |
| ADNI | avtot4 | F1 | |
| ADNI | avtot5 | F1 | |
| ADNI | avtot6 | F5 | |
| ADNI | avtotb | F1 | |
| ADNI | avdel30min | F5 | |
| ADNI | avdeltot | F4 | |
| ADNI | q1score | F1 | |
| ADNI | q4score | F4 | |
| ADNI | q7score | F2 | |
| ADNI | q8score | | |
| ADNI | mmhospit | | |
| ADNI | mmfloor | | |
| ADNI | mmarea | | |
| ADNI | bft1 | | Immediate—ball, flag, tree collapsed |
| ADNI | bft2 | | Delayed—ball, flag, tree collapsed |
| ADNI | imm1sum | F1 | |
| ADNI | Imm2sum | F1 | |
| ADNI | delsum | | |

Executive functioning

ACT: Final model was a data driven bifactor model with CFI = 0.948, TLI = 0.929, and RMSEA = 0.064. The following items were included in the CFA analysis (Supplemental Table 10):

**Supplemental Table 10. Items and secondary structure for executive functioning for the ACT study**

| Study | Variable | Description | Comments | Secondary Structure |
|---|---|---|---|---|
| ACT | mat_attn | Mattis Dementia Rating Scale, Attention score | | |
| ACT | mat_conc | Mattis Dementia Rating Scale, Concentration score | | |
| ACT | mat_ip | Mattis Dementia Rating Scale, initiation / perseveration score | | |
| ACT | tr_a_tm | Trails A | | F1 |
| ACT | tr_b_tm | Trails B | | F1 |
| ACT | clockdr | Clock | | |
| ACT-CASI | dbsum | repeat numbers backward 1–3 | Repeat numbers backward—3 trials collapsed | F2 |
| ACT-CASI | subtra | Subtraction 1–3 | Subtraction—3 trials collapsed | F2 |
| ACT-CASI | sim | similarities | | |
| ACT-CASI | jgmt | judgement | | |

ADNI: Final model was a theory driven methods-effects bifactor model with CFI = 0.951, TLI = 0.946, and RMSEA = 0.041. The following items were included in the CFA analysis (Supplemental Table 11):

**Supplemental Table 11. Items and secondary structure for executive functioning for the ADNI study**

| Study | Variable | Description | Comments | Secondary Structure |
|---|---|---|---|---|
| ADNI | clockcirc | Approximately circular face | | |
| ADNI | clocksym | Symmetry of number placement | | F2 |

| | | | | |
|---|---|---|---|---|
| ADNI | clocknum | Correctness of numbers | | F2 |
| ADNI | clockhand | Presence of the two hands | | |
| ADNI | clocktime | Presence of the two hands, set to ten after eleven | | |
| ADNI | dspanbac | Backward Total Correct | | F4 |
| ADNI | traascor | Part A Time to Complete | | F3 |
| ADNI | trabscor | Part B Time to complete | | F3 |
| ADNI | digitscor | Digit Symbol Total Correct | | F1 |
| ADNI | dspanfor | Digit Span Forward Total Correct | | F4 |
| ADNI | q13score | Number cancellation task | | F1 |
| ADNI | absmeas | Abstraction: watch-ruler | | |
| ADNI | abstran | Abstraction: train-bicycle | | |
| ADNI | trails | MoCA Trails | | |
| ADNI | digback | Digits Backward | 5 trials collapsed | |
| ADNI | serial | Serial 7 total | | |
| ADNI | digfor | Digits Forward | | |
| ADNI | letters | List of Letters/Tapping: # Errors | | |

ROS/MAP: Final model was a theory driven methods-effects model with CFI = 0.975, TLI = 0.960, and RMSEA = 0.064. The following items were included in the CFA analysis (Supplemental Table 12):

**Supplemental Table 12. Items and secondary structure for memory for the ROS and MAP studies**

| **Study** | **Variable** | **Description** | **Comments** | **Secondary structure** |
|---|---|---|---|---|
| ROS/MAP | AA | Which piece would complete the pattern… | 4 A patterns merged | |
| ROS/MAP | BB | Which piece would complete the pattern… | 8 B patterns merged | |
| ROS/MAP | Q12bmme | Spell WORLD backwards | | |

| | | | | |
|---|---|---|---|---|
| ROS/MAP | DigBak | digits backward | combined 12 items | |
| ROS/MAP | cts_sdmt | symbol digits modality (oral) | | F1 |
| ROS/MAP | cts_nccrtd | Number comparison | | F1 |
| ROS/MAP | DigFor | digits forward | combined 12 items | |

**Supplemental Table 13. Items and secondary structure for memory**

| Study | Variable | Description | Secondary structure |
|---|---|---|---|
| ACT, ADNI | traascor | Trails A | F3 |
| ACT, ADNI | trabscor | Trails B | F3 |
| ADNI, ROS/MAP | dspanfor | Digit Span Forward: Total Correct | F4 |
| ADNI, ROS/MAP | dspanbac | Digit Span Backward: Total Correct | F4 |
| ROS/MAP | AA | Which piece would complete the pattern… | |
| ROS/MAP | BB | Which piece would complete the pattern… | |
| ROS/MAP | Q12bmme | Spell WORLD backwards | |
| ROS/MAP | cts_sdmt | symbol digits modality (oral) | F6 |
| ROS/MAP | cts_nccrtd | Number comparison | F6 |
| ACT | mat_attn | Mattis Dementia Rating Scale | |
| ACT | mat_conc | Mattis Dementia Rating Scale | |
| ACT | mat_ip | Mattis Dementia Rating Scale | |
| ACT | clockdr | Clock | |
| ACT-CASI | dbsum | repeat numbers backward | F5 |
| ACT-CASI | subtra | subtraction | F5 |
| ACT-CASI | sim | similar types | |
| ACT-CASI | jgmt | judgement | |
| ADNI | clockcirc | Approximately circular face | |
| ADNI | clocksym | Symmetry of number placement | F2 |

| | | | |
|---|---|---|---|
| ADNI | clocknum | Correctness of numbers | F2 |
| ADNI | clockhand | Presence of the two hands | |
| ADNI | clocktime | Presence of the two hands, set to 10 after 11 | |
| ADNI | digitscor | Digit Symbol Total Correct | F1 |
| ADNI | q13score | Number cancellation task | F1 |
| ADNI | absmeas | Abstraction: watch–ruler | |
| ADNI | abstran | Abstraction: train–bicycle | |
| ADNI | trails | Trails | |
| ADNI | digback | Digits Backward | |
| ADNI | serial | Serial 7 | |
| ADNI | digfor | Digits Forward | |
| ADNI | letters | List of Letters/Tapping: # Errors | |

Language

ACT: Final model was a data driven bifactor model with CFI = 0.956, TLI = 0.943, and RMSEA = 0.055. The following items were included in the CFA analysis (Supplemental Table 14):

**Supplemental Table 14. Items and secondary structure for language for the ACT study**

| Study | Variable | Description | Secondary Structure |
|---|---|---|---|
| ACT | bnt_adpr* | Boston Naming Test – 10-item version | F1 |
| ACT | bnt_cer * | Boston Naming Test – 15-item version | F1 |
| ACT | v_flu_t | Verbal Fluency | |
| ACT-CASI | animal | animals with 4 legs | |
| ACT-CASI | rpta | repeat phrase 1 | F2 |
| ACT-CASI | rptb | repeat phrase 2 | F2 |
| ACT-CASI | cas_read | read and follow a command | |
| ACT-CASI | cas_writ | write a sentence | |

| | | | |
|---|---|---|---|
| ACT-CASI | cmd | obey oral commands | |
| ACT-CASI | body | identify parts of body | |
| ACT-CASI | obja | identify objects—1 | F3 |
| ACT-CASI | objb | identify objects—2 | F3 |

* ACT administers all 15 items from the CERAD version of the Boston Naming Test (bnt_cer) and another 8 distinct items from a long version of the Boston Naming Test (bnt_adpr).

ADNI: Final model was a theory driven methods-effects bifactor model with CFI = 0.951, TLI = 0.946, and RMSEA = 0.041. The following items were included in the CFA analysis (Supplemental Table 15):

**Supplemental Table 15. Items and secondary structure for language for the ADNI study**

| Study | Variable | Description | Secondary structure |
|---|---|---|---|
| ADNI | catanimsc | Category Fluency (Animals) —Total Correct | F1 |
| ADNI | catvegesc | Category Fluency (VegSupplemental Tables) —Total Correct | |
| ADNI | bnttotal | Total Number Correct (1+3) | F1 |
| ADNI | q2score | ADAS Commands | |
| ADNI | q5score | ADAS Naming | F1 |
| ADNI | q6score | Ideational Praxis—score | |
| ADNI | mmwatch | Show wrist watch, ask: What is this? | |
| ADNI | mmpencil | Show pencil, ask: What is this? | |
| ADNI | mmrepeat | Say: Repeat after me: no ifs, ands, or buts. | |
| ADNI | mmhand | Takes paper in right hand | |
| ADNI | mmfold | Folds paper in half | |
| ADNI | mmonflr | Puts paper on floor | |
| ADNI | mmread | Present the piece of paper which reads—CLOSE YOUR EYES—and say: Read this and | |
| ADNI | mmwrite | Give the participant a blank piece of paper and say: Write a sentence. | |
| ADNI | camel | Camel | |

| | | |
|---|---|---|
| ADNI | lion | Lion |
| ADNI | rhino | Rhinoceros |
| ADNI | repeat1 | Repeat Sentence. |
| ADNI | repeat2 | Repeat Sentence. |
| ADNI | ffluency | Letter Fluency—F: Total number of correct words |

ROS/MAP: Final model was a data driven model with CFI = 0.932, TLI = 0.924, and RMSEA = 0.036. The following items were included in the CFA analysis (Supplemental Table 16):

**Supplemental Table 16. Items and secondary structure for language for the ROS and MAP studies**

| Study | Variable | Description | Secondary Structure |
|---|---|---|---|
| ROS/MAP | Q12amme | Spell WORLD forwards | |
| ROS/MAP | Q14mme | [SHOW WRIST WATCH] What is this called? | |
| ROS/MAP | Q15mme | [SHOW PENCIL] What is this called? | |
| ROS/MAP | Q16mme | Repeat a phrase | |
| ROS/MAP | Q17mme | Read the words on this card, then do what it says | |
| ROS/MAP | paper | Takes piece of paper | |
| ROS/MAP | folds | Folds paper in half | |
| ROS/MAP | places | Places paper in lap | |
| ROS/MAP | Q19mme | Write any complete sentence | |
| ROS/MAP | dnaming | What is the name of this object? | |
| ROS/MAP | clothing | all of the things that belong in that category | |
| ROS/MAP | animals | all of the things that belong in that category | F1 |
| ROS/MAP | fruits | all of the things that belong in that category | F1 |
| ROS/MAP | sink1 | Will a board sink in water? | |
| ROS/MAP | sink2 | Will a stone sink in water? | |
| ROS/MAP | hammer1 | Is a hammer good for cutting wood? | |

| | | |
|---|---|---|
| ROS/MAP | hammer2 | Can you use a hammer to pound nails? |
| ROS/MAP | flour1 | Do two pounds of flour weigh more than one? |
| ROS/MAP | flour2 | Is one pound of flour heavier than two? |
| ROS/MAP | boots1 | Will water go through a good pair of rubber boots? |
| ROS/MAP | boots2 | Will a good pair of rubber boots keep water out? |

**Supplemental Table 17. Co-calibration of language across ACT, ADNI, ROS/MAP**

| Study | Variable | Description | Secondary structure |
|---|---|---|---|
| ACT, ADNI, ROS/MAP | read | Read the words on this card, then do it | |
| ACT, ADNI, ROS/MAP | cmd | Paper, fold, place on floor combined | |
| ACT, ADNI, ROS/MAP | catanim | Category Fluency (Animals)—Total Correct | F3 |
| ACT, ROS/MAP | bnt_name | Boston Naming: Name of this object? | F2 |
| ADNI, ROS/MAP | watch | [SHOW WRIST WATCH] What is this called? | |
| ADNI, ROS/MAP | pencil | [SHOW PENCIL] What is this called? | |
| ADNI, ROS/MAP | repeat | I would like you to repeat a phrase after me | |
| ADNI, ROS/MAP | write | Write any complete sentence on this piece of | |
| ROS/MAP | Q12amme | Spell WORLD forwards | |
| ROS/MAP | clothing | all of the things that belong in that category | |
| ROS/MAP | fruits | all of the things that belong in that category | F3 |
| ROS/MAP | sink1 | Will a board sink in water? | |
| ROS/MAP | sink2 | Will a stone sink in water? | |
| ROS/MAP | hammer1 | Is a hammer good for cutting wood? | |
| ROS/MAP | hammer2 | Can you use a hammer to pound nails? | |
| ROS/MAP | flour1 | Do two pounds of flour weigh more than one? | |
| ROS/MAP | flour2 | Is one pound of flour heavier than two? | |
| ROS/MAP | boots1 | Will water go through a good pair of rubber boots? | |
| ROS/MAP | boots2 | Will a good pair of rubber boots keep water out? | |

| | | | |
|---|---|---|---|
| ACT | bnt_adpr | Boston Naming Test | F2 |
| ACT-CASI | animal | animals with 4 legs | F3 |
| ACT-CASI | rpta | repeat something | |
| ACT-CASI | rptb | repeat something | |
| ACT-CASI | cas_writ | write something | |
| ACT-CASI | body | identify part of body | |
| ACT-CASI | obja | identify object—1 | F1 |
| ACT-CASI | objb | identify object—2 | F1 |
| ADNI | catvegesc | Category Fluency (VegSupplemental Tables) —Total Correct | F3 |
| ADNI | bnttotal | Total Number Correct (1+3) | F3 |
| ADNI | q2score | ADAS Commands | |
| ADNI | q5score | ADAS Naming | F3 |
| ADNI | q6score | Ideational Praxis—score | |
| ADNI | camel | Camel | |
| ADNI | lion | Lion | |
| ADNI | rhino | Rhinoceros | |
| ADNI | repeat1 | Repeat Sentence. | |
| ADNI | repeat2 | Repeat Sentence. | |
| ADNI | ffluency | Letter Fluency—F: Total # of correct words | |

Visuospatial FUNCTIONING

ACT: Final model was a data driven bifactor model with CFI = 0.993, TLI = 0.987, and RMSEA = 0.031. The following items were included in the CFA analysis (Supplemental Table 18):

**Supplemental Table 18. Items and secondary structure for visuospatial functioning for the ACT study**

| **Study** | **Variable** | **Description** | **Secondary Structure** |
|---|---|---|---|

| | | | |
|---|---|---|---|
| ACT | mat_cons | Mattis Dementia Rating Scale—constructional praxis score | |
| ACT | cp_in_ci | Constructional Praxis—circle | F1 |
| ACT | cp_in_di | Constructional Praxis—diamond | F1 |
| ACT | cp_in_re | Constructional Praxis—rectangles | |
| ACT | cp_in_cu | Constructional Praxis—cube | |
| ACT-CASI | draw | Copy interlocking pentagons | |

ADNI: Final model was a single factor model with CFI = 0.958, TLI = 0.937, and RMSEA = 0.060. The following items were included in the CFA analysis (Supplemental Table 19):

**Supplemental Table 19. Items and secondary structure for visuospatial functioning for the ADNI study**

| Study | Variable | Description | Secondary Structure |
|---|---|---|---|
| ADNI | copycirc | Clock copy: Approximately circular face | |
| ADNI | copysym | Symmetry of number placement | |
| ADNI | copynum | Correctness of numbers | |
| ADNI | copyhand | Presence of the two hands | |
| ADNI | copytime | Presence of the two hands, set to ten after eleven | |
| ADNI | q3score | Constructional Praxis—score | |
| ADNI | mmdraw | Present the participant with the Cnstrn Stimulus page. Say: Copy this design | |

ROS/MAP: Final model was a single factor model with CFI = 0.948, TLI = 0.940, and RMSEA = 0.040. The following items were included in the CFA analysis (Supplemental Table 20):

**Supplemental Table 20. Items and secondary structure for visuospatial functioning for the ROS and MAP studies**

| Study | Variable | Description | Secondary Structure |
|---|---|---|---|

| | | |
|---|---|---|
| ROS/MAP | Q20mme | Please copy the drawing on this piece of paper |
| ROS/MAP | Line1…15 | Which two lines point in the same direction…? |

Co-calibration of visuospatial ability across ACT, ADNI, ROS/MAP

Single factor models were selected for ADNI and ROS/MAP, while a bifactor model with a single residual correlation was chosen for the ACT study. The pair of items with a residual correlation from ACT was unique to ACT and not present in either of the other studies. Our final model was thus a bifactor model that included only a single residual correlation for the pair of items from the ACT study; all other items, including all of the items administered in ROS/MAP and ADNI, only had loadings on the general factor.

**Supplemental Table 21: Co-calibration of visuospatial ability across ACT, ADNI, and ROS/MAP.**

| Study | Variable | Description | Secondary Structure |
|---|---|---|---|
| ACT, ADNI, ROS/MAP | Q20mme | Please copy the drawing on this piece of paper | |
| ROS/MAP | Line1–15 | Which two lines point in the same direction… | |
| ACT | mat_cons | Mattis Dementia Rating Scale | |
| ACT | cp_in_ci | Constructional Praxis—circle | F1 |
| ACT | cp_in_di | Constructional Praxis—diamond | F1 |
| ACT | cp_in_re | Constructional Praxis—rectangles | |
| ACT | cp_in_cu | Constructional Praxis—cube | |
| ADNI | copycirc | Clock copy: Approx circular face | |
| ADNI | copysym | Symmetry of number placement | |
| ADNI | copynum | Correctness of numbers | |
| ADNI | copyhand | Presence of the two hands | |
| ADNI | copytime | Presence of the two hands, set to ten after eleven | |
| ADNI | q3score | Constructional Praxis—score | |

**Addition of University of Pittsburgh study to the pipeline**

As detailed above, items where the PITT item was the same as an item with parameters from the ACT/ADNI/ROS-MAP analyses, we used those previously co-calibrated parameters. For items administered only to PITT participants, we freely estimated item parameters from the data set.

**Supplemental Table 22. Memory specification for the PITT dataset**

| Study | Variable | Description | Secondary Structure |
|---|---|---|---|
| ROS/MAP, PITT | wrec | Word recognition trial 1 | F2 |
| ROS/MAP, PITT | wrec2 | Word recognition trial 2 | F2 |
| ROS/MAP, PITT | wrec3 | Word recognition trial 3 | F2 |
| ROS/MAP, PITT | wrecde | Word recognition—delayed | |
| PITT | targets | Word recognition—target correct | |
| PITT | foils | Word recognition—foils correct | |
| PITT | reyim | Rey figure—immediate recall | F3 |
| PITT | reyde | Rey figure—delayed | F3 |
| ACT, ADNI, PITT | logimem | Logical Memory A1 | F1 |
| ACT, ADNI, PITT | memunits | Logical Memory A2 | F1 |
| ACT, PITT | mattism | Mattis DRS—memory | |

**Supplemental Table 23. Executive function specification for the PITT dataset**

| Study | Variable | Description | Secondary Structure |
|---|---|---|---|
| ROS/MAP, ADNI, PITT | spansb | Digit span—backwards | F1 |
| ACT, ADNI, PITT | trailas | Trail A—time | F2 |
| ACT, ADNI, PITT | trailbs | Trail B—time | F2 |
| PITT | mbar | Abstract reasoning | |
| ACT, PITT | mattisip | Mattis DRS—initiation/perseveration | |

| | | | |
|---|---|---|---|
| ACT, PITT | mconcep | Mattis DRS—conceptualization | |
| ROS/MAP, ADNI, PITT | spansf | Digit span—forward | F1 |
| ACT, PITT | mattisa | Mattis DRS—attention | |
| PITT | stpcw | Stroop—Color,Word | |

**Supplemental Table 24. Language specification for the PITT dataset**

| Study | Variable | Description | Secondary Structure |
|---|---|---|---|
| ACT, ADNI, ROS/MAP, PITT | fluen | Fluency test—animals | F1 |
| PITT | fluenb | Fluency test—birds | F1 |
| PITT | fluend | Fluency test—dogs | F1 |
| ADNI, PITT | fluenf | Fluency test—letter F | F3 |
| PITT | fluena | Fluency test—letter A | F3 |
| PITT | fluens | Fluency test—letter S | F3 |
| PITT | stpw | Stroop—Word | F2 |
| PITT | stpc | Stroop—Color | F2 |
| ADNI, PITT | veg | Category fluency—veg | F1 |
| ADNI, PITT | boston | Boston Naming Test total | F1 |

**Supplemental Table 25. Visuospatial functioning specification for the PITT dataset.**

| Study | Variable | description | Secondary Structure |
|---|---|---|---|
| ACT, ADNI, ROS/MAP, PITT | pentagon | draw intersecting pentagons | |
| PITT | reyco | Rey figure—copy | |
| PITT | blkdsn | Block design | |
| ACT, PITT | mconst | Mattis DRS—construction | |

**Addition of Amsterdam Dementia Cohort (ADC) study to the pipeline**

As detailed above, items where the ADC item was the same as an item with parameters from the ACT/ADNI/ROS-MAP/PITT analyses, we used those previously co-calibrated parameters. For items administered only to ADC participants, we freely estimated item parameters from the data set.

**Supplemental Table 26 Memory specification for the ADC dataset**

| Study | Variable | Description | Secondary Structure |
|---|---|---|---|
| ADNI, ADC | ravtot1 | Rey AVLT Trial 1 Total | F1 |
| ADNI, ADC | ravtot2 | Rey AVLT Trial 2 Total | F1 |
| ADNI, ADC | ravtot3 | Rey AVLT Trial 3 Total | F1 |
| ADNI, ADC | ravtot4 | Rey AVLT Trial 4 Total | F1 |
| ADNI, ADC | ravtot5 | Rey AVLT Trial 5 Total | F1 |
| ADC | ravdel | Rey AVLT – delayed recall | |
| ADC | vat_a1 | Verbal association Test A1 | F2 |
| ADC | vat_a2 | Verbal association Test A2 | F2 |
| ADC | vat_b1 | Verbal association Test B1 | F3 |
| ADC | vat_b2 | Verbal association Test B2 | F3 |

ADNI administered two versions (different word lists) of RAVLT (A & B) while ADC administers a Dutch version of the RAVLT. We performed a series of psychometric analyses of RAVLT items looking at thresholds and loadings using ADC only item parameters and using co-calibrated parameters. We concluded that RAVLT Trials 1 to 5 items can be considered “anchor” items compared to ADNI RAVLT version A.

**Supplemental Table 27. Executive function specification for the ADC dataset**

| Study | Variable | Description | Secondary Structure |
|---|---|---|---|
| ACT, ADNI, PITT | trailas | Trail A—time | F1 |

| | | | |
|---|---|---|---|
| ACT, ADNI, PITT | trailbs | Trail B—time | F1 |
| ADC | dspanfor | Digit span—forward | F2 |
| ADC | dspanbac | Digit span—backwards | F2 |
| ADC | ldst | Letter digit substitution test | |
| ADC | fabat | Frontal assessment battery | |
| ADC | stroopcw | Stroop—Color,Word | |

**Supplemental Table 28. Language specification for the ADC dataset**

| Study | Variable | Description | Secondary Structure |
|---|---|---|---|
| ACT, ADNI, ROS/MAP, PITT, ADC | catanim | Fluency test—animals | |
| ADC | vatn | Verbal association test naming | F1 |
| ADC | naming | Confrontational naming | F1 |
| ADNI, PITT, ADC | ffluency | Fluency test—letter D | F3 |
| PITT, ADC | fluena | Fluency test—letter A | F3 |
| PITT, ADC | fluent | Fluency test—letter T | F3 |
| ADC | stroopw | Stroop—Word | F2 |
| ADC | stroopc | Stroop—Color | F2 |
| ADC | compq | Comparative questions | |

After literature search, we found that the Dutch version of letter fluency items starting with letters D/A/T are equivalent to F/A/S in English. We performed sensitivity analyses to make sure that a) frequencies of these items are similar across the Dutch and English version and b) item parameters (thresholds and loadings) are similar across the two languages. We concluded that these three items can be considered "anchor" items.

**Supplemental Table 29. Visuospatial functioning specification for the ADC dataset.**

| Study | Variable | description | Secondary Structure |
|---|---|---|---|

| | | |
|---|---|---|
| ACT, ADNI, ROS/MAP, PITT, ADC | mmdraw | draw intersecting pentagons |
| ADC | notcount | Dot counting |
| ADC | fraglett | Fragmented letters |
| ADC | numloc | Number location |

**Cohorts**

**The ACT/eMERGE Studies (ACT)** The ACT cohort is an urban and suburban elderly population from a stable HMO that includes 2,581 cognitively intact subjects age ≥ 65 who were enrolled between 1994 and 1998. An additional 811 subjects were enrolled in 2000-2002 using the same methods except oversampling clinics with more minorities. More recently, a Continuous Enrollment strategy was initiated in which new subjects are contacted, screened and enrolled to keep 2000 active at-risk person-years accruing in each calendar year. This resulted in an enrollment of 4,146 participants as of May 2009. All clinical data are reviewed at a consensus conference. Dementia onset is assigned half way between the prior biennial and the exam that diagnosed dementia. Enrollment for the eMERGE Study began in 2007. A waiver of consent was obtained from the IRB to enroll deceased ACT participants.

**The ADNI Study (ADNI 1/GO/2)** ADNI is a longitudinal, multi-site observational study including people with Alzheimer's disease, people with mild cognitive impairment (MCI), and elderly individuals with normal cognition assessing clinical and cognitive measures, MRI and PET scans (FDG and 11C PIB) and blood and CNS biomarkers. For this study, ADNI contributed data on 607 Alzheimer's disease cases and 325 healthy controls with Alzheimer's disease -free status confirmed as of most recent follow-up. Alzheimer's disease subjects were between the ages of 65–90, had an MMSE score of 20–26 inclusive, met NINCDS/ADRDA criteria for probable Alzheimer's disease, and had an MRI consistent with the diagnosis of Alzheimer's disease. Control subjects had MMSE scores between 28 and 30 and a Clinical Dementia Rating of 0 without symptoms of depression, MCI or other dementia and no current use of psychoactive medications. According to the ADNI protocol, subjects were ascertained at regular intervals over 3 years, but for the purpose of our analysis we only used the final ascertainment status to classify case-control status.

Data used in the preparation of this article were obtained from ADNI database (http://adni.loni.ucla.edu). The ADNI was launched in 2003 by the National Institute on Aging (NIA), the National Institute of Biomedical Imaging and Bioengineering (NIBIB), the Food and Drug Administration (FDA), private pharmaceutical companies and non-profit organizations, as a $60 million, 5-year public-private partnership. The primary goal of ADNI has been to test whether serial magnetic resonance imaging (MRI), positron emission tomography (PET), other biological markers, and clinical and neuropsychological assessment can be combined to measure the progression of mild cognitive impairment (MCI) and early Alzheimer's disease Alzheimer's disease. Determination of sensitive and specific markers of very early Alzheimer's disease progression is intended to aid researchers and clinicians to develop new treatments and monitor their effectiveness, as well as lessen the time and cost of clinical trials. The Principal Investigator of this initiative is Michael W. Weiner, MD, VA Medical Center and University of California—San Francisco. ADNI is the result of efforts of many co-investigators from a broad range of academic institutions and private corporations, and subjects have been recruited from over 50 sites across the U.S. and Canada. The initial goal of ADNI was to recruit 800 subjects but ADNI has been followed by ADNI-GO and ADNI-2. To date these three protocols have recruited over 1500 adults, ages 55 to 90, to participate in the research, consisting of cognitively normal older individuals, people with early or late MCI, and people with early Alzheimer's disease. The follow up duration of each group is specified in the protocols for ADNI-1, ADNI-2 and ADNI-GO. Subjects originally recruited for ADNI-1 and ADNI-GO had the option to be followed in ADNI-2. For up-to-date information, see www.adni-info.org.

**The ROS/MAP Studies** ROS/MAP are two community-based cohort studies. The ROS has been on-going since 1993, with a rolling admission. Through July of 2010, 1,139 older nuns, priests, and brothers from across the United States initially free of dementia who agreed to annual clinical evaluation and brain donation at the time of death completed their baseline evaluation. The MAP has been on-going since 1997, also with a rolling admission. Through July of 2010, 1,356 older persons from across northeastern Illinois initially free of dementia who agreed to annual clinical evaluation and organ donation at the time of death completed their baseline evaluation.

**University of Pittsburgh (PITT)** The University of Pittsburgh data set contains 1,271 Caucasian Alzheimer's disease cases (of which 277 were autopsy-confirmed) recruited by the University of Pittsburgh Alzheimer's Disease Research Center, and 841 Caucasian, cognitively normal elderly controls ages 60 and older (2 were autopsy-confirmed). All Alzheimer’s disease cases met NINCDS/ADRDA criteria for probable or definite Alzheimer’s disease. Additional details of the cohort used for GWAS have been previously published. We limited analyses from the Pittsburgh site to those with a Clinical Dementia Rating (CDR) of 0.5 or 1.0, since stability of cognitively-defined Alzheimer’s disease subgroups in more advanced degrees of severity has not been established.

**Amsterdam Dementia Cohort (ADC).** The Alzheimer center of the Amsterdam UMC - VU University Medical Center opened in 2000 and was initiated to combine both patient care and research. Together, to date (2018), all patients forming the Amsterdam Dementia Cohort number almost 6,000 individuals. This cohort focusses on these research lines: 1) early diagnosis, 2) heterogeneity, and 3) vascular factors. Among the most important research efforts that have also impacted patients' lives and/or the research field, we count the development of novel, easy to use diagnostic measures such as visual rating scales for MRI and the Amsterdam IADL Questionnaire, insight in different subgroups of AD, and findings on incidence and clinical sequelae of microbleeds https://www.alzheimercentrum.nl/.
